# Reference genome and transcriptome informed by the sex chromosome complement of the sample increases ability to detect sex differences in gene expression from RNA-Seq data

**DOI:** 10.1101/668376

**Authors:** Kimberly C. Olney, Sarah M. Brotman, Jocelyn P. Andrews, Valeria A. Valverde-Vesling, Melissa A. Wilson

**Affiliations:** School of Life Sciences, Arizona State University, Tempe AZ 85282 USA; Center for Evolution and Medicine, Arizona State University, Tempe AZ 85282 USA; Center for Mechanisms of Evolution, The Biodesign Institute, Arizona State University, Tempe AZ 85282 USA; Department of Genetics, University of North Carolina, Chapel Hill NC 27599 USA; College of Osteopathic Medicine of the Pacific, Western University of Health Sciences, Pomona CA 91766 USA

**Keywords:** RNA-Seq, sex chromosomes, differential expression, transcriptome, mapping, alignment, pseudo-alignment, quantification

## Abstract

**Background:** Human X and Y chromosomes share an evolutionary origin and, as a consequence, sequence similarity. We investigated whether sequence homology between the X and Y chromosomes affects alignment of RNA-Seq reads and estimates of differential expression. We tested the effects of using reference genomes and reference transcriptomes informed by the sex chromosome complement of the sample’s genome on measurements of RNA-Seq abundance and sex differences in expression.

**Results:** The default genome includes the entire human reference genome (GRCh38), including the entire sequence of the X and Y chromosomes. We created two sex chromosome complement informed reference genomes. One sex chromosome complement informed reference genome was used for samples that lacked a Y chromosome; for this reference genome version, we hard-masked the entire Y chromosome. For the other sex chromosome complement informed reference genome, to be used for samples with a Y chromosome, we hard-masked only the pseudoautosomal regions of the Y chromosome, because these regions are duplicated identically in the reference genome on the X chromosome. We analyzed transcript abundance in the whole blood, brain cortex, breast, liver, and thyroid tissues from 20 genetic female (46, XX) and 20 genetic male (46, XY) samples. Each sample was aligned twice; once to the default reference genome and then independently aligned to a reference genome informed by the sex chromosome complement of the sample, repeated using two different read aligners, HISAT and STAR. We then quantified sex differences in gene expression using featureCounts to get the raw count estimates followed by Limma/Voom for normalization and differential expression. We additionally created sex chromosome complement informed transcriptome references for use in pseudo-alignment using Salmon. Transcript abundance was quantified twice for each sample; once to the default target transcripts and then independently to target transcripts informed by the sex chromosome complement of the sample.

**Conclusions:** We show that regardless of the choice of read aligner, using an alignment protocol informed by the sex chromosome complement of the sample results in higher expression estimates on the pseudoautosomal regions of the X chromosome in both genetic male and genetic female samples, as well as an increased number of unique genes being called as differentially expressed between the sexes. We additionally show that using a pseudo-alignment approach informed on the sex chromosome complement of the sample eliminates Y-linked expression in female XX samples.

**Author summary:** The human X and Y chromosomes share an evolutionary origin and sequence homology, including regions of 100% identity; this sequence homology can result in reads misaligning between the sex chromosomes, X and Y. We hypothesized that misalignment of reads on the sex chromosomes would confound estimates of transcript abundance if the sex chromosome complement of the sample is not accounted for during the alignment step. For example, because of shared sequence similarity, X-linked reads could misalign to the Y chromosome. This is expected to result in reduced expression for regions between X and Y that share high levels of homology. For this reason, we tested the effect of using a default reference genome versus a reference genome informed by the sex chromosome complement of the sample on estimates of transcript abundance in human RNA-Seq samples from whole blood, brain cortex, breast, liver, and thyroid tissues of 20 genetic female (46, XX) and 20 genetic male (46, XY) samples. We found that using a reference genome with the sex chromosome complement of the sample resulted in higher measurements of X-linked gene transcription for both male and female samples and more differentially expressed genes on the X and Y chromosomes. We additionally investigated the use of a sex chromosome complement informed transcriptome reference index for alignment free quantification protocols. We observed no Y-linked expression in female XX samples only when the transcript quantification was performed using a transcriptome reference index informed on the sex chromosome complement of the sample. We recommend that future studies requiring aligning RNA-Seq reads to a reference genome or pseudo-alignment with a transcriptome reference should consider the sex chromosome complement of their samples prior to running default pipelines.

## Background

Sex differences in aspects of human biology, such as development, physiology, metabolism, and disease susceptibility are partially driven by sex specific gene regulation (Arnold et al., 2012; Khramtsova et al., 2018; Raznahan et al., 2018; Traglia et al., 2017). There are reported sex differences in gene expression across human tissues(Gershoni and Pietrokovski, 2017; Goldstein et al., 2014; Shi et al., 2016) and while some may be attributed to hormones and environment, there are documented genome-wide sex differences in expression based solely on the sex chromosome complement (Arnold and Chen, 2009). However, accounting for the sex chromosome complement of the sample in quantifying gene expression has been limited due to shared sequence homology between the sex chromosomes, X and Y, that can confound gene expression estimates.

The X and Y chromosomes share an evolutionary origin: mammalian X and Y chromosomes originated from a pair of indistinguishable autosomes ∼180-210 million years ago that acquired the sex-determining genes (Charlesworth, 1991; Lahn and Page, 1999; Ross et al., 2005). The human X and Y chromosomes formed in two different segments: a) one that is shared across all mammals called the X-conserved region (XCR) and b) the X-added region (XAR) that is shared across all eutherian animals (Ross et al., 2005). The sex chromosomes, X and Y, previously recombined along their entire lengths, but due to recombination suppression from Y chromosome-specific inversions (Lahn and Page, 1999; Pandey et al., 2013), now only recombine at the tips in the pseudoautosomal regions (PAR) PAR1 and PAR2 (Charlesworth, 1991; Lahn and Page, 1999; Ross et al., 2005). PAR1 is ∼2.78 million bases (Mb) and PAR2 is ∼0.33 Mb; these sequences are 100% identical between X and Y (Aken et al., 2017; Charchar et al., 2003; Ross et al., 2005) (Figure 1A). The PAR1 is a remnant of the XAR Ross et al. 2005) and shared among eutherians, while the PAR2 is recently added and human-specific (Charchar et al., 2003). Other regions of high sequence similarity between X and Y include the X-transposed-region (XTR) with 98.78% homology (Veerappa et al., 2013) (Figure 1A). The XTR formed from an X chromosome to Y chromosome duplication event following the human-chimpanzee divergence (Ross et al., 2005; Skaletsky et al., 2003). Thus, the evolution of the X and Y chromosomes has resulted in a pair of chromosomes that are diverged, but still share some regions of high sequence similarity.

**Figure 1.**
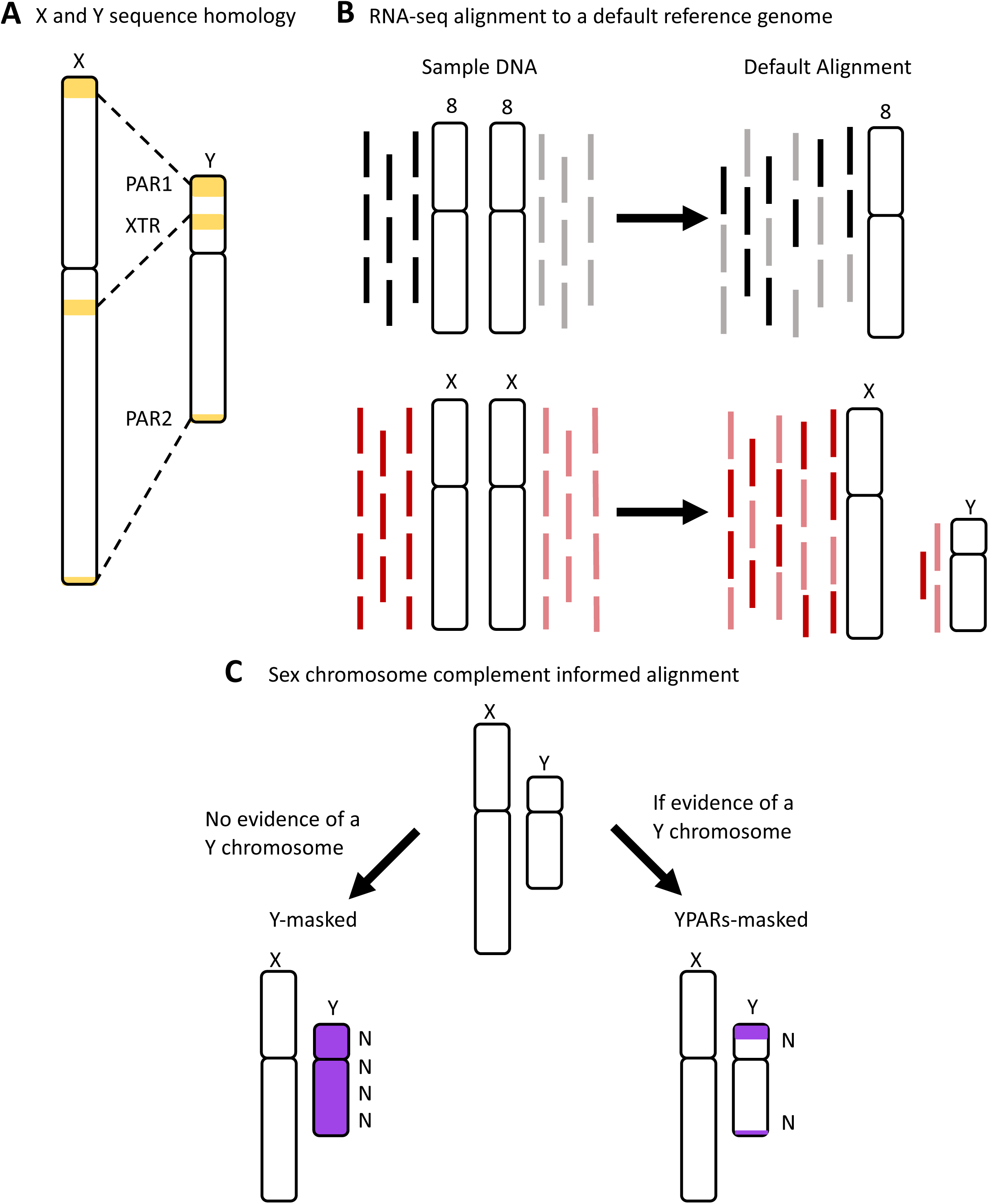
Homology between the human X and Y chromosomes where misaligning could occur. **A)** High sequence homology exists between the human X and Y chromosomes in three regions: 100% sequence identity for the pseudoautosomal regions (PARs), PAR1 and PAR2, and ∼99% sequence homology in the X-transposed region (XTR). The X chromosome PAR1 is ∼2.78 million bases (Mb) extending from X:10,001 to 2,781,479 and the X chromosome PAR2 is ∼0.33 Mb extending from X:155,701,383 to 156,030,895. The X chromosome PAR1 and PAR2 are identical in sequence to the Y chromosome PAR1 Y:10,001 – 2,781,479 and PAR2 Y:56,887,903 – 57,217,415. **B)** Using a standard alignment approach will result in reads misaligning between regions of high sequence homology on the sex chromosomes. **C)** Using a reference genome that is informed by the genetic sex of the sample may help to reduce misaligning between the X and Y chromosomes. In humans, samples without evidence of a Y chromosome should be aligned to a Y-masked reference genome and samples with evidence of a Y should be aligned to a YPARs-masked reference genome.

To infer which genes or transcripts are expressed, RNA-Seq reads can be aligned to a reference genome. The abundance of reads mapped to a transcript is reflective of the amount of expression of that transcript. RNA-Seq methods rely on aligning reads to an available high quality reference genome sequence, but this remains a challenge due to the intrinsic complexity in the transcriptome of regions with a high level of homology (Piskol et al., 2013). By default, the GRCh38 version of the human reference genome includes both the X and Y chromosomes, which is used to align RNA-Seq reads from both male XY and female XX samples. It is known that sequence reads from DNA will misalign along the sex chromosomes affecting downstream analyses (Webster et al., 2019). However, this has not been tested using RNA-Seq data and the effects on differential expression analysis are not known. Considering the increasing number of human RNA-Seq consortium datasets (e.g., the Genotype-Tissue Expression project (GTEx) (GTEx Consortium, 2015), The Cancer Genome Atlas (TCGA) (Cancer Genome Atlas Research Network et al., 2013), Geuvadis project (Lappalainen et al., 2013), and Simons Genome Diversity Project (Mallick et al., 2016)), there is an urgent need to understand how aligning to a default reference genome that includes both X and Y may affect estimates of gene expression on the sex chromosomes (Khramtsova et al., 2018; Tukiainen et al., 2016). We hypothesize that regions of high sequence similarity will result in misaligning of RNA-Seq reads and reduced expression estimates (Figure 1A & B).

Here, we tested the effect of sex chromosome complement informed read alignment on the quantified levels of gene expression and the ability to detect sex-biased gene expression. We utilized data from the GTEx project, focusing on five tissues, whole blood, brain cortex, breast, liver, and thyroid, which are known to exhibit sex differences in gene expression (Gershoni and Pietrokovski 2017; R. Li and Singh 2014; de Perrot et al. 2000; Melé et al. 2015; Mayne et al. 2016). Many genes have been reported to be differentially expressed between male and female brain samples (Gershoni and Pietrokovski, 2017; Goldstein et al., 2014; Shi et al., 2016) and differential expression in blood samples between males and females has also been documented (Gershoni and Pietrokovski, 2017; Goldstein et al., 2014). An analysis of all GTEx tissue samples reported that breast mammary gland tissues are the most sex differentially expressed tissue (Gershoni and Pietrokovski, 2017). It has also been reported that there are sex disparities in thyroid cancer (Rahbari et al., 2010) and liver cancer (Natri et al., 2019; Naugler et al., 2007) suggesting possible sex differences in gene expression. We used whole blood, brain cortex, breast, liver, and thyroid tissues from 20 genetic male (46, XY) and 20 genetic female (46, XX) individuals for a total of 200 samples evenly distributed among tissues. Male and female samples, for each tissue, were age-matched between the sexes and only included samples of age 55 to 70. We aligned all samples to a default reference genome that includes both the X and Y chromosomes and to a reference genome that is informed on the sex chromosome complement of the genome: Male XY samples were aligned to a reference genome that includes both the X and Y chromosome, where the Y chromosome PAR1 and PAR2 are hard-masked with Ns (Figure 1C) so that reads will align uniquely to the X PAR sequences. Conversely, female XX samples were aligned to a reference genome where the entirety of the Y chromosome is hard-masked (Figure 1C). We tested two different read aligners, HISAT (Kim et al., 2015) and STAR (Dobin et al., 2013), to account for variation between alignment methods and measured differential expression using Limma/Voom (Law et al., 2014). We found that using a sex chromosome complement informed reference genome for aligning RNA-Seq reads increased expression estimates on the pseudoautosomal regions of the X chromosome in both male XY and female XX samples and uniquely identified differentially expressed genes.

We additionally investigated the effect of transcriptome references on pseudo-alignment methods. We quantified abundance using Salmon (Patro et al., 2017) in male and female brain cortex samples twice, once to a default reference transcriptome index that includes both the X and Y chromosome linked transcripts and to a reference transcriptome index that is informed on the sex chromosome complement of the sample. We found that using a sex chromosome complement informed reference transcriptome index for RNA-Seq pseudo-alignment quantification eliminated Y-linked expression estimates in female XX samples, that were observed in the default approach.

Regardless of alignment or pseudo-alignment approach, we recommended carefully considering the annotations of the sex chromosomes in the references used, as theses will affect quantifications and differential expression estimates, especially of sex chromosome linked genes.

## Methods

### Building sex chromosome complement informed reference genomes

All GRCh38.p10 unmasked genomic DNA sequences, including autosomes 1-22, X, Y, mitochondrial DNA (mtDNA), and contigs were downloaded from ensembl.org release 92 (Aken et al., 2017). The default reference genome here includes all 22 autosomes, mtDNA, the X chromosome, the Y chromosome, and contigs. For the two sex chromosome complement informed reference assemblies, we included all 22 autosomes, mtDNA, and contigs from the default reference and a) one with the Y chromosome either hard-masked for the “Y-masked reference genome” or b) one with the pseudoautosomal regions, PAR1 and PAR2, hard-masked on the Y chromosome for “YPARs-masked reference genome” (Figure 1C). Hard-masking with Ns will force reads to not align to those masked regions in the genome. Masking the entire Y chromosome for the sex chromosome complement informed reference genome, Y-masked, was accomplished by changing all the Y chromosome nucleotides [ATGC] to N using a sed command in linux. YPARs-masked was created by hard-masking the Y PAR1: 6001-2699520 and the Y PAR2: 154931044-155260560 regions. The GRCh38.p10 Y PAR1 and Y PAR2 chromosome start and end location was defined using Ensembl GRCh38 Y PAR definitions (Aken et al., 2017). After creating the Y chromosome PAR1 and PAR2 masked fasta files, we concatenated all the Y chromosome regions together to create a YPARs-masked reference genome. After creating the GRCh38.p10 default reference genome and the two sex chromosome complement informed reference genomes, we indexed the reference genomes and created a dictionary for each using HISAT version 2.1.0 (Kim et al., 2015) hisat2-build -f option and STAR version 2.5.2 (Dobin et al., 2013), using option --genomeDir and --sjdbGTFfile. Reference genome indexing was followed by picard tools version 1.119 CreateSequenceDictionary (2020), which created a dictionary for each reference genome (Pipeline available on GitHub, https://github.com/SexChrLab/XY_RNAseq).

### Building sex chromosome complement informed transcriptome index

Ensembl’s GRCh38.p10 cDNA reference transcriptome fasta consists of transcript sequences resulting from Ensemble gene predictions. Ensembl’s cDNA was downloaded from ensembl.org release 92 (Aken et al., 2017). The default transcriptome reference includes 199,234 transcripts which includes autosomal, mtDNA, X chromosome, Y chromosome and contig transcripts. The default Ensembl cDNA does not contain Y chromosome PAR linked transcript sequences, it only contains the X chromosome PAR sequence transcripts. For the sex chromosome complement informed reference transcriptome index, we included all 22 autosomes, mtDNA, X, and contigs from the default cDNA transcriptome and we hard-masked all available Y chromosome linked transcript sequences. Hard-masking the Y chromosome linked transcripts was accomplished by changing all the Y chromosome nucleotides [ATGC] to N using a sed command in linux. After downloading the GRCh38.p10 default reference transcriptome and creating the Y-masked sex chromosome complement informed reference transcriptome fasta files, we then generated a decoy-aware transcriptome for each transcriptome reference. For generating the default decoy-aware reference transcriptome, we used the default genome as the decoy sequence. This was accomplished by concatenating the default genome fasta to the end of the default transcriptome fasta to populate the decoy file with the chromosome names, as suggested by Salmon (Patro et al., 2017). The default transcriptome fasta and the default decoy file were then used to create the mapping-based index using the Salmon version 1.2.0 index function (Patro et al., 2017). The Y-masked decoy-aware transcriptome fasta was generated by concatenating the Y-masked genome fasta to the end of the Y-masked transcriptome fasta to populate the decoy file with the chromosome names. The Y-masked transcriptome fasta and the decoy file were then used as inputs for generating the Y-masked mapping-based index using the salmon index function. For both the default and the Y-masked mapping-based index, a k-mer of 31 was used as this was suggested to work well for reads of 75bp.

In addition to the Ensembl reference, we investigated the effects of a sex chromosome complement reference transcriptome index using the gencode transcript reference fasta GRCh38.p12 that contains 206,694 transcripts which includes autosomal, mtDNA, X, Y and contigs. The gencode transcriptome reference includes both the X and Y PAR transcripts (J et al., 2012). Following the same parameters for the Ensembl decoy-aware transcriptome, we created two gencode sex chromosome complement decoy-aware transcriptome references, in addition to a default gencode decoy-aware transcriptome reference. The pipeline is available on GitHub, https://github.com/SexChrLab/XY_RNAseq.

### RNA-Seq samples

From the Genotyping-Tissue Expression (GTEx) Project data, we downloaded SRA files for whole blood, brain cortex, breast, liver, and thyroid tissues from 20 separate genetic female (46, XX) and 20 separate genetic male (46, XY) individuals (Consortium, 2015; GTEx Consortium, 2015) that were age matched between the sexes and ranged from age 55 to 70 (Additional file 1 & 2). Age matching exactly was accomplished using the matchit function in the R package MatchIt (Ho et al. 2011). The GTEx data is described and available through dbGaP under accession phs000424.v6.p1; we received approval to access this data under dbGaP accession #8834. GTEx RNA-Seq samples were sequenced to 76bp reads and the median coverage was ∼82 million (M) reads (Consortium, 2015). Although information about the genetic sex of the samples was provided in the GTEx summary downloads, it was additionally investigated by examining the gene expression of select genes that are known to be differentially expressed between the sexes or are known X-Y homologous genes: *DDX3X, DDX3Y, PCDH11X, PCDH11Y, USP9X, USP9Y, ZFX, ZFY, UTX, UTY, XIST*, and *SRY* (Figure 2; Additional file 3 & 4).

**Figure 2.**
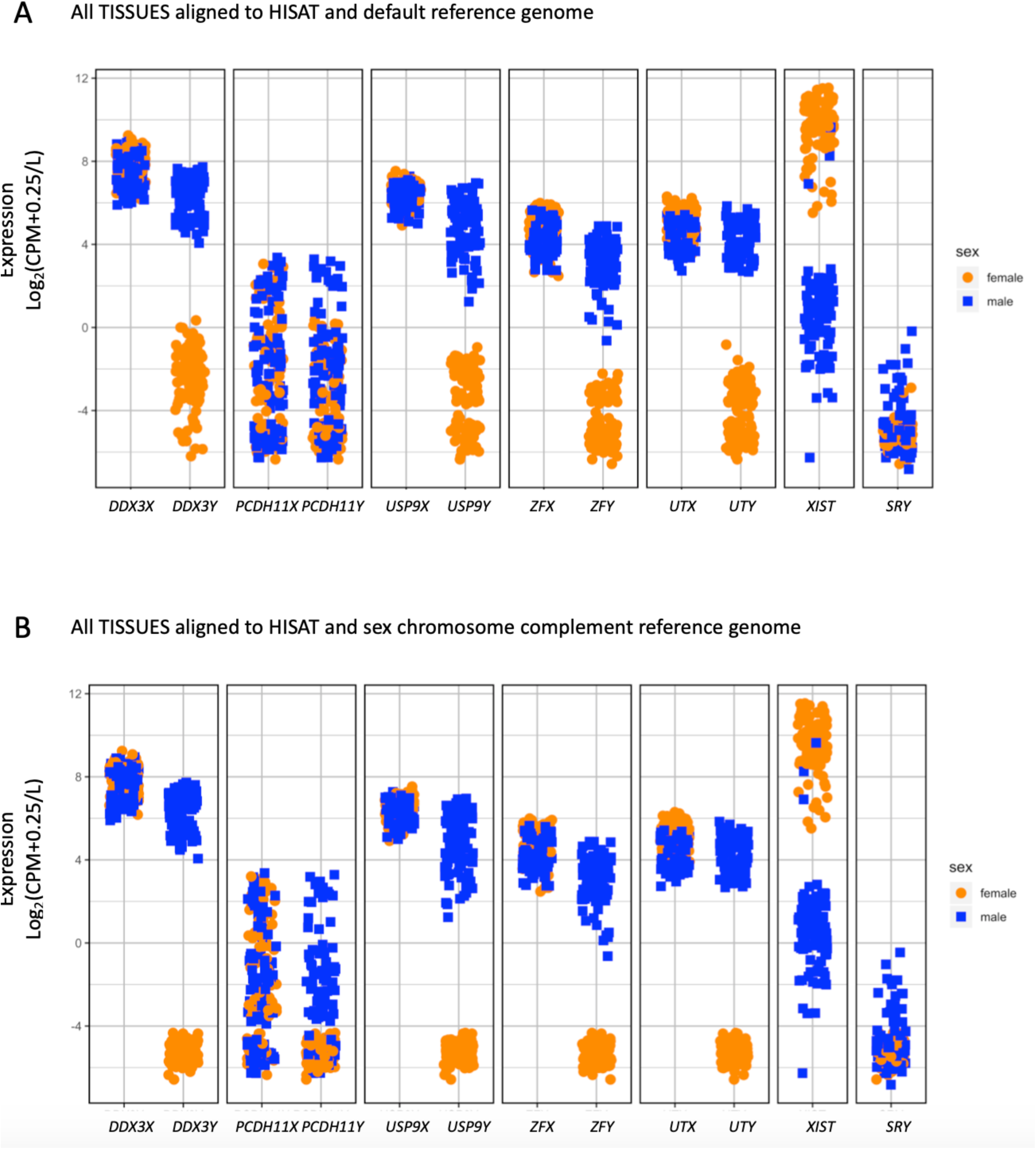
Genetic sex of RNA-Seq samples. We investigated gene expression, log2(CPM+0.25/L), of XY homologous genes (*DDX3X/Y, PCDH11X/Y, USP9X/Y, ZFX/Y, UTX/Y*), and *XIST*, and *SRY* in all samples from all tissues analyzed here from genetic males (blue squares) and genetic females (orange circles) **A)** when aligned to a default reference genome, and **B)** when aligned to a sex chromosome complement informed reference genome, using HISAT as the read aligner.

### RNA-Seq trimming and quality filtering

RNA-Seq sample data was converted from sequence read archive (sra) format to the paired-end FASTQ format using the SRA toolkit (Leinonen et al., 2011). Quality of the samples’ raw sequencing reads was examined using FastQC (Andrews) and MultiQC. Subsequently, adapter sequences were removed using Trimmomatic version 0.36 (Bolger et al., 2014). More specifically, reads were trimmed to remove bases with a quality score less than 10 for the leading strand and less than 25 for the trailing strand, applying a sliding window of 4 with a mean PHRED quality of 30 required in the window and a minimum read length of 40 bases.

### RNA-Seq read alignment

Following trimming, paired RNA-Seq reads from all samples were aligned to the default reference genome. Unpaired RNA-Seq reads were not used for alignment. Reads from the female (46, XX) samples were aligned to the Y-masked reference genome and reads from male (46, XY) individuals were aligned to the YPARs-masked reference genome. Read alignment was performed using HISAT version 2.1.0 (Kim et al., 2015), keeping all parameters the same, only changing the reference genome used, as described above. Read alignment was additionally performed using STAR version 2.5.2 (Dobin et al., 2013), where all samples were aligned to a default reference genome and to a reference genome informed on the sex chromosome complement, keeping all parameters the same (Pipeline available on GitHub, https://github.com/SexChrLab/XY_RNAseq).

### Processing of RNA-Seq alignment files

Aligned RNA-Seq samples from HISAT and STAR were output in Sequence Alignment Map (SAM) format and converted to Binary Alignment Map (BAM) format using bamtools version 2.4.0 (Li et al., 2009). Summaries on the BAM files including the number of reads mapped were computed using bamtools version 2.4.0 package (Barnett et al., 2011). RNA-Seq BAM files were indexed, sorted, duplicates were marked, and read groups added using bamtools, samtools, and Picard (Barnett et al., 2011; Li et al., 2009, 2020). All RNA-Seq BAM files were indexed using the default reference genome using Picard ReorderSam (2020), this was done so that all samples would include all chromosomes in the index files. Aligning XX samples to a Y-masked reference genome using HISAT indexes would result in no Y chromosome information in the aligned BAM and BAM index bai files. For downstream analysis, some tools require that all samples have the same chromosomes, which is why we hard-masked rather than removed. Reindexing the BAM files to the default reference genome does not alter the read alignment, and thus does not alter our comparison between default and sex chromosome complement informed alignment.

### Gene expression level quantification

Read counts for each gene across all autosomes, sex chromosomes, mtDNA, and contigs were generated using featureCounts version 1.5.2 (Liao et al., 2014) for all aligned and processed RNA-Seq BAM files. Female XX samples when aligned to a sex chromosome complement informed reference genome will show zero counts for Y-linked genes, but will still include those genes in the raw counts file. This is an essential step for downstream differential expression analysis between males and females to keep the total genes the same between the sexes for comparison. Only rows that matched gene feature type in Ensembl Homo_sapiens.GRCh38.89.gtf gene annotation (Aken et al., 2017) were included for read counting. There are 2,283 genes annotated on the X chromosome and a total of 56,571 genes across the entire genome for GRCh38 version of the human reference genome (Aken et al., 2017). Only primary alignments were counted and specified using the --primary option in featureCounts.

### RNA-seq quantification for transcriptome index

Transcript quantification for trimmed paired RNA-seq brain cortex samples were estimated twice, once to a default decoy-aware reference transcriptome index and once to a sex chromosome complement informed decoy-aware reference transcriptome index using Salmon with the – validateMappings flag. Salmon’s –validateMappings adopts a scheme for finding protentional mapping loci of a read using a chain algorithm introduced in minimap2 (Li, 2018). Transcript quantification for female (46, XX) samples was estimated using a Y-masked reference transcriptome index and male (46, XY) transcript quantification was estimated using a Y PAR masked reference transcriptome index when the Y PAR sequence information was available for the transcriptome build. This was repeated for both the Ensembl and the gencode cDNA transcriptome builds, keeping all parameters the same, only changing the reference transcriptome index used, as described above.

### DGEList object

Differential expression analysis was performed using the limma/voom pipeline (Law et al., 2014) which has been shown to be a robust differential expression software package (Costa-Silva et al., 2017; Seyednasrollah et al., 2015) for both reference-based and pseudo-alignment quantification. Quantified read counts from each sample for the reference-based quantification were generated from featureCounts were combined into a count matrix, each row representing a unique gene ID and each column representing the gene counts for each unique sample. This was repeated for each tissue type and read into R using the DGEList function in the R limma package (Love et al., 2014). A sample-level information file related to the genetic sex of the sample, male or female, and the reference genome used for alignment, default or sex chromosome complement informed, was created and corresponds to the columns of the count matrix described above.

Pseudo-aligned transcript read counts from each brain cortex sample quantified using Salmon were combined into a count matrix using tximport (Soneson et al., 2015) with each row representing a unique transcript ID and each column representing the transcript counts for each unique sample. To create length scaled transcripts per million (TPM) values to pass into limma, tximport function lengthScaledTPM was employed (Soneson et al., 2015). The reference assembly annotation file was read into R using tximport function makeTxDbFromGFF. Following this, a key of the transcript ID corresponding to the gene ID was created was created using the keys function (Soneson et al., 2015). Gene level TPM values were then generated using the tx2gene function. This was repeated for the Ensembl and the gencode default and sex chromosome complement informed transcriptome quantification estimates.

### Multidimensional Scaling

Multidimensional Scaling (MDS) was performed using the DGEList-object containing gene expression count information for each sample. MDS plots were generated using the plotMDS function in in the R limma package (Law et al., 2014). The distance between each pair of samples is shown as the log_2_ fold change between the samples. The analysis was done for each tissue separately using all shared common variable genes for dimensions (dim) 1 & 2 and dim 2 & 3. Samples that did not cluster with reported sex or clustered in unexpected ways in either dim1, 2 or 3 were removed from all downstream analysis (Additional file 5). MDS plots for each tissue containing the samples that were used for quality control are located in Additional file 6. Briefly, one male XY whole blood did not cluster with any of the other samples and was removed. One female XX breast sample clustered with the opposite sex and was thus removed. In brain cortex, three male XY brain cortex samples didn’t cluster neatly with the other male XY samples in dim 1 & 2 were thus removed. Another male brain cortex sample, although clustered with other male samples, had the lowest number of sequencing remaining after trimming for quality, 23.9M, and thus was also removed. To keep the number of samples in each sex roughly equal, four female XX brain cortex samples were randomly selected for removal. For liver and thyroid tissue, no samples appeared to cluster in any unexpected ways and thus no liver or thyroid tissue samples were removed. For all aligners the first component of variation in the MDS plot is explained by the sex of the sample (Figure 3).

**Figure 3.**
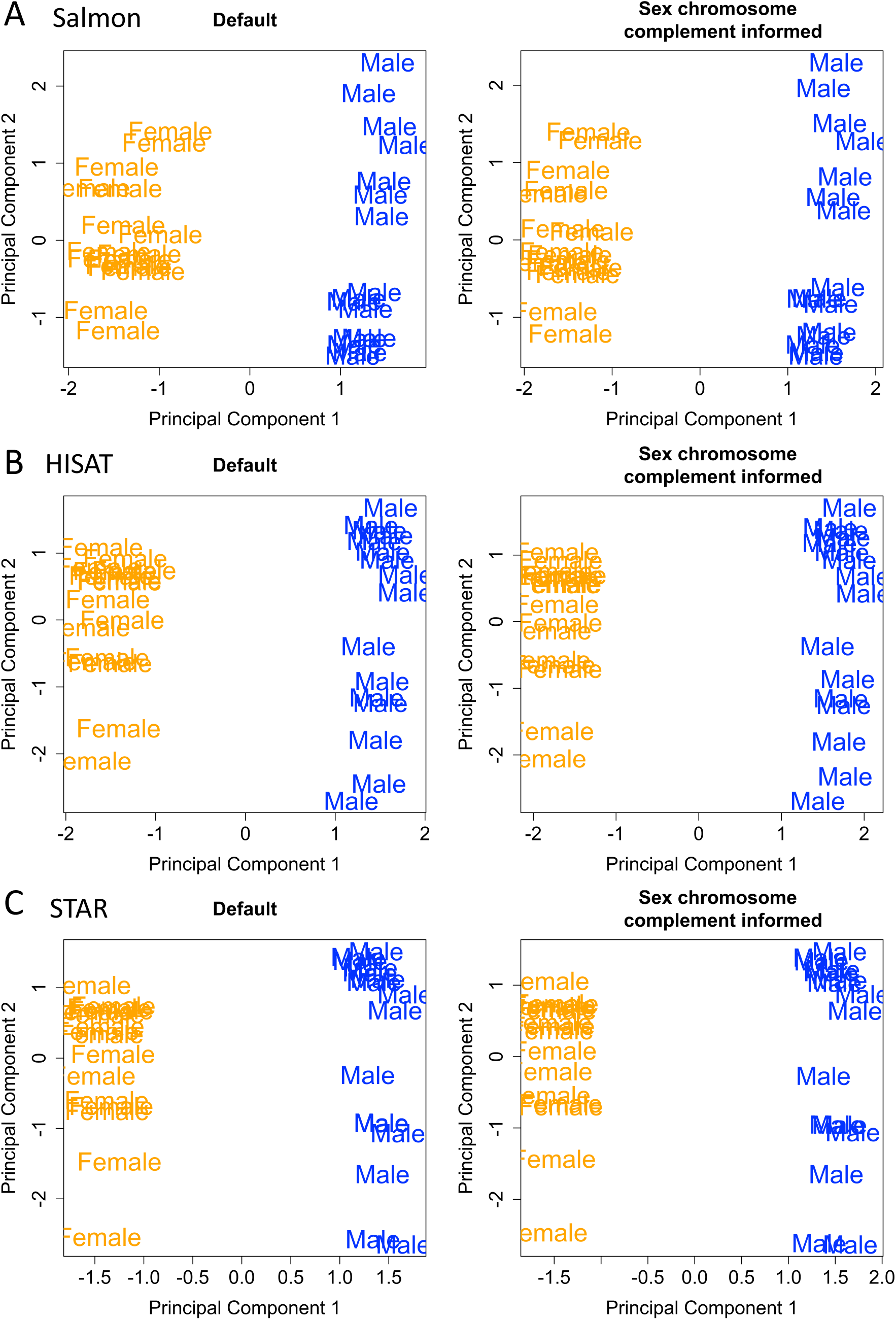
Multidimensional scaling for the top 100 most variable genes. We investigated multidimensional scaling for the top 100 most common variable genes in brain cortex samples. **A)** Salmon pseudo-alignment with Ensembl transcriptome reference **B)** HISAT read aligner and **C)** STAR read aligner when quantifying using both the default and the sex chromosome complement informed reference. The most variation in the data is explained by the sex of the sample.

### Differential expression

Using edgeR (Robinson et al., 2010), raw counts were normalized to adjust for compositional differences between the RNA-Seq libraries using the voom normalize quantile function, which normalizes the reads by the method of trimmed mean of values (TMM) (Law et al., 2014). Counts were then transformed to log_2_(CPM+0.25/L), where CPM is counts per million, L is library size, and 0.25 is a prior count to avoid taking the log of zero (Robinson et al., 2010). For this dataset, the average library size is about 79.76 million, therefore L is 79.76. Thus, the minimum log_2_(CPM+0.25/L) value for each sample, representing zero transcripts, is log_2_(0+0.25/15) = −8.32.

A mean minimum of 1 CPM, or the equivalent of 0 in log_2_(CPM+2/L), in at least one sex per tissue comparison was required for the gene to be kept for downstream analysis. A CPM value of 1 was used in our analysis to separate expressed genes from unexpressed genes, meaning that in a library size of ∼79.76 million reads, there are at least an average of 79 counts in at least one sex. After filtering for a minimum CPM, 53,804 out of the 56,571 quantified genes were retained for the whole blood samples, 53,822 for brain cortex, 54,184 for breast, 53,830 for liver, and 53,848 for thyroid. A linear model was fitted to the DGEList-object, which contains the filtered and normalized gene counts for each sample, using the limma lmfit function which will fit a separate model to the expression values for each gene (Law et al., 2014).

For differential expression analysis a design matrix containing the genetic sex of the sample (male or female) and which reference genome the sample was aligned to (default or sex chromosome complement informed) was created for each tissue type for contrasts of pairwise comparisons between the sexes. Pairwise contrasts were generated using limma makecontrasts function (Law et al., 2014). We identified genes that exhibited significant expression differences defined using an Benjamini-Hochberg adjusted p-value cutoff that is less than 0.01 (1%) to account for multiple testing in pairwise comparisons between conditions using limma/voom decideTests vebayesfit (Law et al., 2014). A conservative adjusted p-value cutoff of less than 0.01 was chosen to be highly confident in the genes that were called as differentially expressed when comparing between reference genomes used for alignment. Pipeline available on GitHub, https://github.com/SexChrLab/XY_RNAseq.

### GO analysis

We examined differences and similarities in gene enrichment terms between the differentially expressed genes obtained from the differential expression analyses of the samples aligned to the default and sex chromosome complement informed reference genomes, to investigate if the biological interpretation would change depending on the reference genome the samples were aligned to. We investigated gene ontology enrichment for lists of genes that were identified as showing overexpression in one sex versus the other sex for whole blood, brain cortex, breast, liver, and thyroid samples (adjusted p-value < 0.01). We used the GOrilla webtool, which utilizes a hypergeometric distribution to identify enriched GO terms (Eden et al., 2009). A modified Fisher exact p-value cutoff < 0.001 was used to select significantly enriched terms (Eden et al., 2009).

## Results

### RNA-Seq reads aligned to autosomes do not vary much between reference genomes

We compared total mapped reads when reads were aligned to a default reference genome and to a reference genome informed on the sex chromosome complement. Reads mapped across the whole genome, including the sex chromosomes, decreased when samples were aligned to a reference genome informed on the sex chromosome complement, paired t-test p-value < 0.05 (Additional files 7 – 9). This was true regardless of the read aligner used, HISAT or STAR, or of the sex of the sample, XY or XX. To test the effects of realignment on an autosome, we selected chromosome 8, because of its similar size to chromosome X. Overall, there is a slight mean increase in reads mapping to chromosome 8 when samples are aligned to a sex chromosome complement informed reference genome compared to aligning to a default reference genome (Additional file 9). For female XX samples, the mean increase in reads mapping for chromosome 8 was 42.2 reads for whole blood, 50.25 for brain cortex, 109.9 for breast, 68.5 for liver, and 98.2 for thyroid (Additional file 9), which was significant using a paired t-test, p-value < 0.05 in all tissues (Additional file 9). Male XY samples also showed a mean increase in reads mapping for chromosome 8. The mean increase in reads mapping to chromosome 8 for male whole blood samples was 0.84, 2.38 for brain cortex, 5.58 for breast, 3.2 for liver, and 5 for thyroid (Additional file 9). There was a significant increase, p-value < 0.05 paired t-test, for reads mapping to chromosome 8 for male brain cortex, breast, liver, and thyroid samples. There was no significant increase in reads mapping for male whole blood for chromosome 8 (Additional file 9).

### Reads aligned to the X chromosome increase in both XX and XY samples when using a sex chromosome complement informed reference genome

We found that when reads were aligned to a reference genome informed by the sex chromosome complement for both male XY and female XX tissue samples, reads on the X chromosome increased by ∼0.12% when aligned using HISAT. For all tissues and both sexes we observe an average increase of 1,991 reads on chromosome X. We observe an increase in reads mapping to the X chromosome for all tissues and for each sex, which was significant using a paired t-test, p-value < 0.05 (Additional file 9). Reads on the Y chromosome decreased 100% (67,033 reads on average) across all female XX samples and by ∼57.32% (69,947 reads on average) across all male XY samples when aligned using HISAT (Additional file 7 & 9). Similar increases in X chromosome and decreases in Y chromosome reads when aligned to a sex chromosome complement informed reference were observed when STAR was used as the read aligner for both male XY and female XX samples (Additional file 8 & 9).

### Aligning to a sex chromosome complement informed reference genome increases the X chromosome PAR1 and PAR2 expression

We next explored the effect of changes in read alignment on gene expression. There was an increase in pseudoautosomal regions, PAR1 and PAR2, expression when reads were aligned to a reference genome informed on the sex chromosome complement for both male XY and female XX samples (Additional file 10 & 11). We found an average of 2.73 log_2_ fold increase in expression in PAR1 expression for female XX brain cortex samples and 2.75 log_2_ fold increase in expression in PAR1 for male XY brain cortex samples using HISAT (Figure 4). The X-transposed region, XTR, in female XX brain cortex samples showed a 1.22 log_2_ fold increase in expression and no change in male XY brain cortex samples. PAR2 showed an average of 2.13 log_2_ fold increase for female XX brain cortex samples and 2.19 log_2_ fold increase in PAR2 for male XY brain cortex samples using HISAT, with similar results for STAR read aligner (Additional file 10 & 11). Complete lists of the log_2_(CPM+0.25/L) values for each X chromosomal gene and each gene within the whole genome for male XY and female XX samples are in Additional file 12 available on Dryad for download under https://doi.org/10.5061/dryad.xksn02vbv.

**Figure 4.**
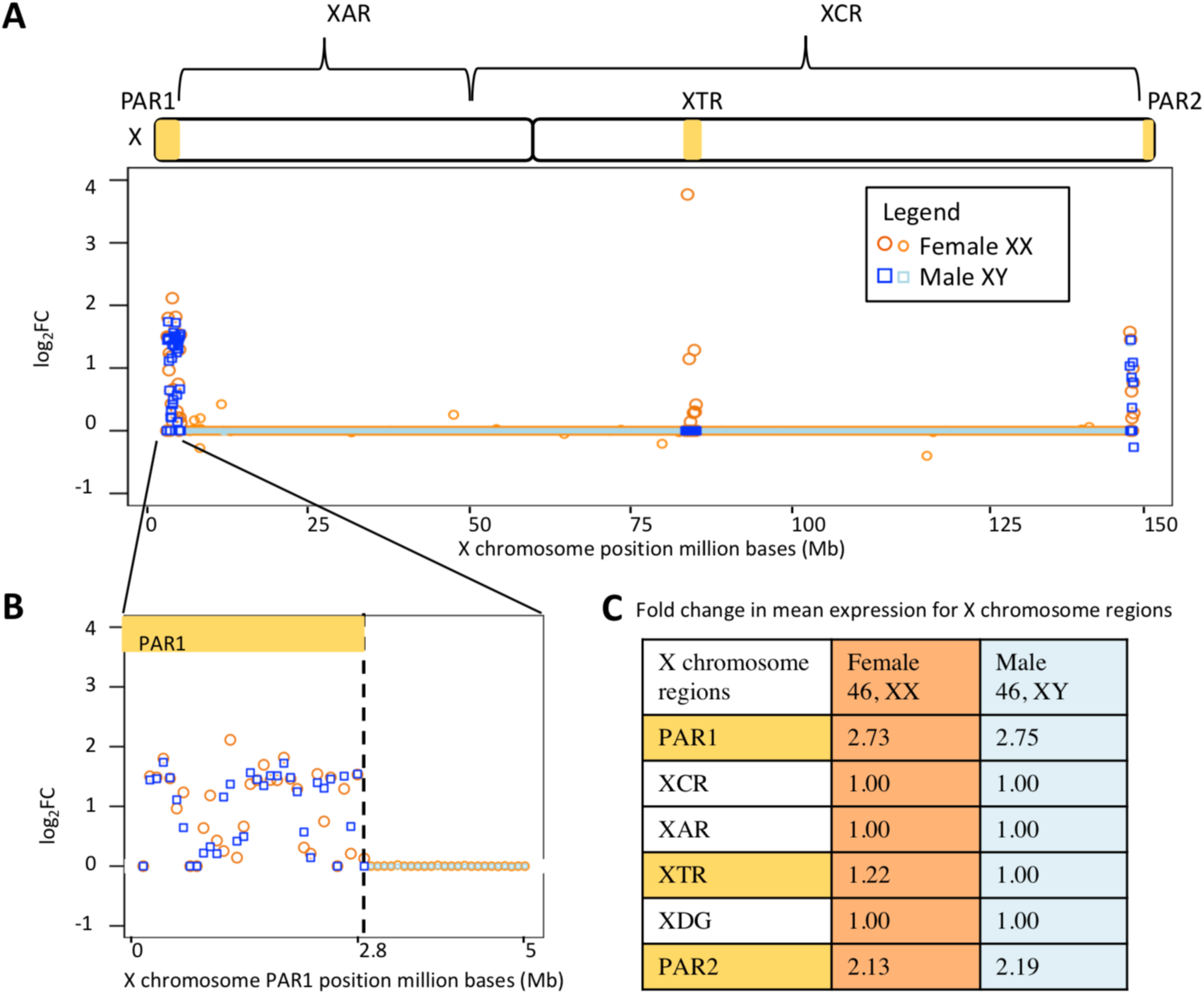
X chromosome RNA-Seq alignment differences in brain cortex. We plot log2 fold change (FC) across **A)** the entire X chromosome and **B)** the first 5 million bases (Mb) and show **C)** average fold change in large genomic regions on the X chromosome between aligning brain cortex using HISAT to the default genome and aligning to a sex chromosome complement informed reference genome. For log2 FC, a value less than zero indicates that the gene showed higher expression when aligned to a default reference genome, while values above zero indicate that the gene shows higher expression when aligned to a reference genome informed by the sex chromosome complement of the sample. Samples from genetic females are plotted in orange circles, while samples from males are plotted in blue squares. Darker shades indicate which gene points are in PAR1, XTR, and PAR2 while lighter shades are used for genes outside of those regions.

### Regions outside the PARs and XTR show little difference in expression between reference genomes

Intriguingly, regions outside the PARs on the X chromosome were less affected by the choice of reference genome. Across the entire X-conserved region, we observed practically no change in estimates of gene expression between the default and sex chromosome complement informed references (e.g., a 0.99 log_2_ fold in male thyroid samples, and 1.00 log_2_ fold change in female brain cortex samples, essentially showing no difference (Additional file 10 & 11)). Additionally, X and Y homologous genes (*AMELX, ARSD, ARSE, ARSF, CASK, GYG2, HSFX1, HSFX2, NLGN4X, OFD1, PCDH11X, PRKX, RBMX, RPS4X, SOX3, STS, TBL1X, TGIF2LX, TMSB4X, TSPYL2, USP9X, VCX, VCX2, VCX3A, VCX3B, ZFX*) showed little to no increase in expression when aligned to a sex chromosome complement informed reference genome compared to aligning to a default reference genome (Additional file 13). *PCDH11X* showed the highest increase in expression for all tissues regardless of read aligner. The log_2_ fold increase in expression for *PCDH11X* for female samples when aligned using HISAT was 0.4, 0.28, 0.33, 0.16, and 0.16 for whole blood, brain cortex, breast, liver, and thyroid, respectively. Other X and Y homologous genes sometimes increased in expression depending on the tissue and sometimes there was no change in expression (Additional file 13). Next to *PCDH11X*, the most increase in expression in an X and Y homologous genes was *VCX3B, NLGN4X*, and *VCX3A. NLGN4X* in whole blood showed a 0.14 log_2_ fold increase when aligned using HISAT. *VCX3B* showed a 0.2 log_2_ fold increase in brain, *NLGN4X* showed a 0.04 log_2_ fold increase in breast, *VCX3A* showed a 0.07 log_2_ fold increase in liver, and *VCX3B* showed a 0.04 log_2_ fold increase in thyroid, when aligned using HISAT (Additional file 13).

### A sex chromosome complement informed reference genome increases the ability to detect sex differences in gene expression

We next investigated how this would affect gene differential expression between the sexes. Generally, we find that more genes are differentially expressed on the sex chromosomes between the sexes when the sex chromosome complements are taken into account. The number of differentially expressed genes on the autosomes remained the same or increased. At a conservative Benjamini-Hochberg adjusted p-value of < 0.01 and aligning with HISAT, we find 4 new genes (3 Y-linked and 1 X-linked) that are only called as differentially expressed between the sexes in the brain cortex when aligned to reference genomes informed on the sex chromosome complement (Figure 5; Additional file 14). We observed similar trends in changes for differential expression between male XY and female XX for whole blood, breast, liver, and thyroid samples using either HISAT or STAR as the aligner (Additional file 14). For example, in whole blood, 3 additional genes are called as being differentially expressed between the sexes using HISAT, while 1 additional gene is called differentially expressed when aligned using STAR. Additionally, when taking sex chromosome complement into account, the number of genes called as differentially expressed between the sexes for the breast samples increased by 13 genes (8 autosomal, 3 X-linked and 2 Y-linked) using HISAT and by 8 genes using STAR (6 autosomal and 2 X-linked) (Additional file 14 & 15). For all tissues, no genes were uniquely called as being differentially expressed between the sexes when aligned to a default reference genome compared to a reference genome informed on the sex chromosome complement (Additional file 14 & 15). Rather, only when samples were aligned to a sex chromosome complement did we observe an increase in the genes called as being differentially expressed (Figure 5; Additional file 14 & 15).

**Figure 5.**
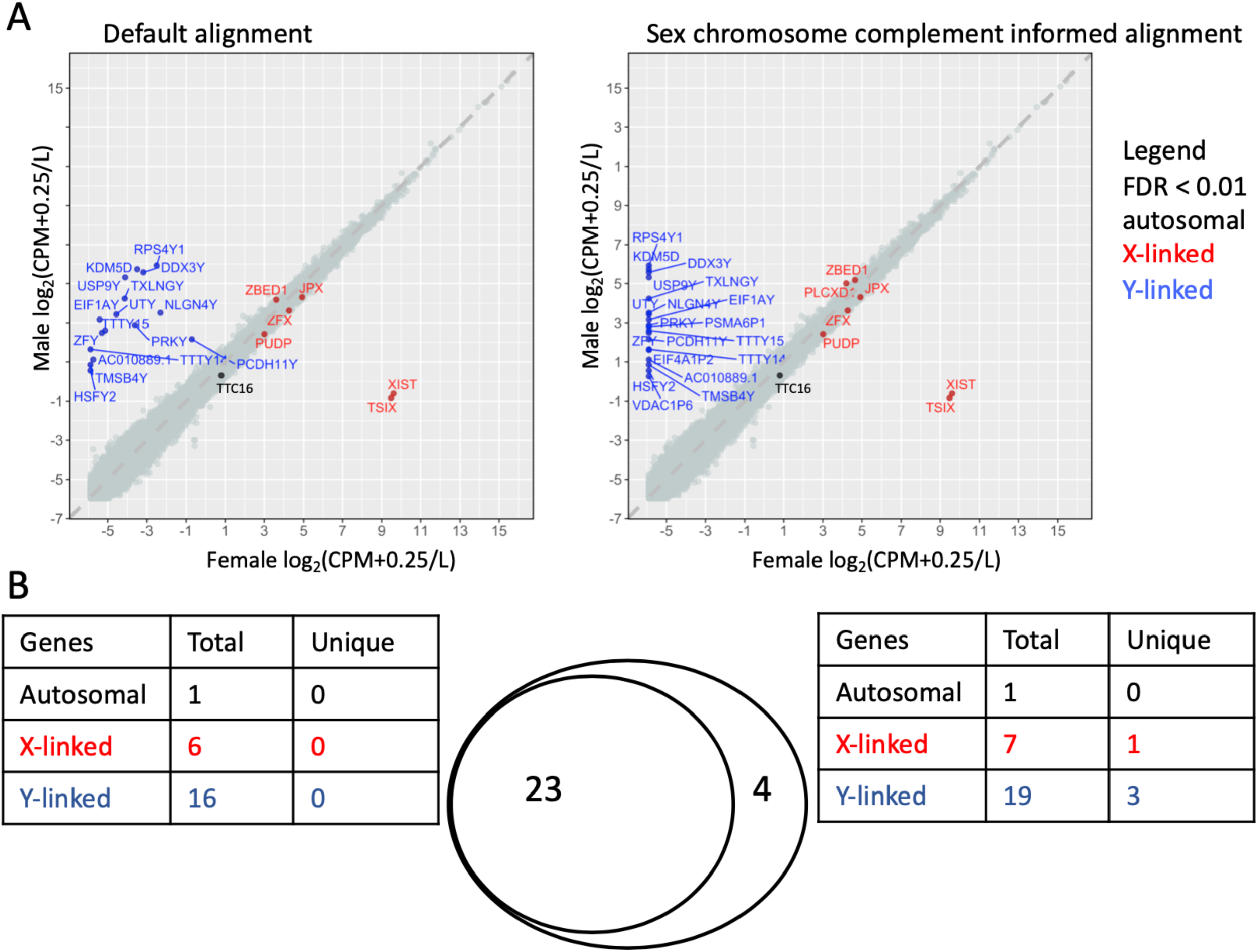
Sex chromosome complement informed alignment calls more sex-linked genes as being differentially expressed. **A)** Sex differences in gene expression, log_2_(CPM+0.25/L), between the twenty samples from genetic males and females are shown when aligning all samples to the default reference genome (left) and a reference genome informed on the sex chromosome complement (right) for brain cortex. Each point represents a gene. Genes that are differentially expressed, adjusted p-value < 0.01 are indicated in black for autosomal genes, blue for Y-linked genes, and red for X-linked genes. **B)** We show overlap between genes that are called as differentially expressed when all samples are aligned to the default genome, and genes that are called as differentially expressed when aligned to a sex chromosome complement informed genome. When samples were aligned to a reference genome informed on the sex chromosome complement, 27 genes were called as differentially expressed between the sexes, of which 4 were uniquely called in the sex chromosome complement informed alignment. There were no genes that were uniquely called as differentially expressed when aligned to a default reference genome.

### Increase in gene enrichment pathways when samples are aligned to a sex chromosome complement informed reference genome

A sex chromosome complement informed reference genome increases the ability to detect genes as differentially expressed between the sexes and thus alters gene enrichment results. When the thyroid samples were aligned using a sex chromosome complement informed reference genome using HISAT, genes up-regulated in male XY samples still show enrichment for positive regulation of transcription from RNA polymerase II (found when aligning to a default reference genome), but additionally find postsynaptic membrane assembly, postsynaptic membrane organization, and vocalization behavior (Additional file 16). These additional GO enrichments in the male XY thyroid samples involve *NRXN1* and *NLGN4Y* genes, both of these genes are located on the Y chromosome. GO enrichment analysis of genes that are more highly expressed in female liver compared to male liver samples, when samples were aligned to a default reference genome using HISAT, were genes involved in modification histone lysine demethylation (Additional file 16). However, when these samples were aligned to a sex chromosome complement informed reference genome, genes upregulated in females were enriched for histone lysine demethylation as well as negative regulation of endopeptidase activity, negative regulation of peptidase activity, cytoplasmic actin-based contraction involved in cell motility (Additional file 16). These additional GO enrichments in the female XX liver samples include the involvement of *KDM6A, DDX3X*, and *VIL1. KDM6A, DDX3X* <u>are X-linked and</u> *<u>VIL1</u>* <u>is on chromosome 2.</u> Whole blood, brain cortex, male liver, and female thyroid samples showed no difference in GO enrichment pathways when using a default reference genome compared to a sex chromosome complement reference genome for alignment when using HISAT with similar results for STAR as the read aligner (Additional file 17). Thus, while there won’t always be a difference, aligning to a sex chromosome complement informed reference genome can increase ability to detect enriched pathways.

### Using sex-linked genes alone is inefficient for determining the sex chromosome complement of a sample

The sex of each sample used in this analysis was provided in the GTEx manifest. We investigated the expression of genes that could be used to infer the sex of the sample. We studied X and Y homologous genes (*DDX3X/Y, PCDH11X/Y, USP9X/Y, ZFX/Y, UTX/Y*), *XIST*, and *SRY* gene expression in male and female whole blood, brain cortex, breast, liver, and thyroid (Figure 2; Additional file 3 & 4). Both males and females are expected to show expression for the X-linked homologs, whereas only XY samples should show expression of the Y-linked homologs. Further, *XIST* expression should only be observed in XX samples and *SRY* should only be expressed in samples with a Y chromosome. Using the default reference genome for aligning samples, we observed a small number of reads aligning to the Y-linked genes in female XX samples, but also observed clustering by sex for *DDX3Y, USP9Y, ZFY*, and *UTY* gene expression (Figure 2). Male XY samples showed expression for *DDX3X, DDX3Y, USP9X, ZFX*, and *UTX* (greater than 5 log_2_(CPM+2/L). Female XX samples showed expression for *XIST* (greater than 4.0 log_2_(CPM+2/L) and male XY samples showed little to no expression for *XIST* (less than 0 log_2_(CPM+2/L) with the exception of 2 male whole blood samples and 1 male liver sample, which showed greater than 5 log_2_(CPM+2/L) expression). In contrast to the default reference genome, when aligned to a sex chromosome complement informed reference genome, samples cluster more distinctly by sex for *DDX3Y, USP9Y*, ZFY, and *UTY*, all showing at least a 4 log_2_(CPM+2/L) difference between the sexes (Figure 2; Additional file 3 & 4). *SRY* is predominantly expressed in the testis (Albrecht et al., 2003; Turner et al., 2011) and typically one would expect *SRY* to show male-specific expression. In our set, we did not observe *SRY* expressed in any sample, and so it could not be used to differentiate between XX and XY samples (Figure 2, Additional file 3 & 14). In contrast, the X-linked gene *XIST* was differentially expressed between genetic males and genetic females in both genome alignments (default and sex chromosome complement informed) for the whole blood, brain cortex, breast, liver, and thyroid samples with the exception of 3 male XY samples. *XIST* expression is important in the X chromosome inactivation process (Carrel and Willard, 2005) and serves to distinguish samples with one X chromosome from those with more than one X chromosome (Tukiainen et al., 2016). However, this does not inform about whether the sample has a Y chromosome or not. For X-Y homologous genes, we do not find sex differences in read alignment with either default or sex chromosome complement informed for the X-linked homolog. When aligned to a default reference genome, female XX samples showed some expression for homologous Y-linked genes, but only presence/absence of Y-linked reads alone is insufficient to determine sex chromosome complement of the sample (Figure 2, Additional file 3).

### No Y-linked transcript expression in female XX samples when quantification was estimated using a transcriptome index informed on the sex chromosome complement

A pseudo-alignment shows similar effects of the reference to that of an alignment approach (Figure 5, Additional files 18 & 19). We observed no Y-linked expression in female XX samples when transcript quantification was estimated using a Y-masked sex chromosome complement reference transcriptome index. This was true for both the Ensembl and gencode pseudo-alignment with a sex chromosome complement reference transcriptome index (Additional files 18 & 19). Interestingly, there was a large difference between the Ensembl and gencode reference files. The transcript IDs in the transcriptome cDNA fasta and the transcript IDs in the annotation file are not one-to-one for the Ensembl assembly (Zhao and Zhang, 2015). There are 190,432 transcript sequences in the Ensembl cDNA fasta file but there are 199,234 transcripts in the Ensembl annotation file. Notably, Ensembl’s cDNA reference transcriptome fastas does not contain known transcripts such as the XIST transcripts (Eyras et al., 2004). The Ensembl reference transcriptome fasta also does not contain the Y PARs transcript sequences, it only contains the X PAR transcript sequences. In contrast, the gencode cDNA reference transcriptome fasta and annotation file both contain 206,694 sequences, including the Y PARs. Regardless of using an Ensembl or gencode transcriptome, female XX sample show Y-linked expression when using a default refence transcriptome index for pseudo-alignment, however the changes necessary for making a sex chromosome complement informed reference are different for the two builds.

## Discussion

For accuracy, the sex chromosome complement of the sample should be taken into account when aligning RNA-Seq reads to reduce misaligning sequences. Neither Ensembl or Gencode human reference genomes are correct for aligning both XX and XY samples. The Ensembl GRCh38 human reference genome includes all 22 autosomes, mtDNA, the X chromosome, the Y chromosome with the Y PARs masked, and contigs (Aken et al., 2017). The Gencode hg19 human reference genome includes everything with no sequences masked (Harrow et al., 2012).

Measurements of X chromosome expression increase for both male XY and female XX whole blood, brain cortex, breast, liver, and thyroid samples when aligned to a sex chromosome complement informed reference genome versus aligning to a default reference genome (Figure 4). While we see increases in measured expression for PAR1 and PAR2 genes in both males and females, we only observe a difference in measured XTR expression in females. This is because while the PARs are 100% identical between the X and Y and so one copy (here we mask the Y-linked copy) should be masked, the XTR is not hard-masked in the YPARs-masked reference genome. The XTR is not identical between the X and Y; it shares 98.78% homology between X and Y but no longer recombines between X and Y (Veerappa et al., 2013) (Figure 1A) and because of this divergence, is therefore not hard-masked when aligning male XY samples. Tukiainen et al., (2016) and others have shown that PAR1 genes have a male bias in expression (Tukiainen et al., 2016). Our findings here support this regardless if the samples were aligned to a default or a sex chromosome complement reference genome (Additional file 11 & 12). Differential expression results changed when using a sex chromosome complement informed alignment compared to using a default alignment. When aligned to a default reference genome, due to sequence similarity, some reads from female XX samples aligned to the Y chromosome (Figure 2; Figure 5). However, when aligned to a reference genome informed by the sex chromosome complement, female XX samples no longer showed Y-linked gene expression, and more Y-linked genes were called as being differentially expressed between the sexes (Figure 2; Figure 5; Additional file 12 & 15). This suggests that if using a default reference genome for aligning RNA-Seq reads, one would miss some Y-linked genes as differentially expressed between the sexes (Figure 5). Furthermore, these Y-linked genes serve in various important biological processes, thus altering the functional interpretation of the sex differences (Additional file 16 & 17). Only when samples were aligned to a sex chromosome complement reference genome did we observe more genes called as differentially expressed between the sexes (Additional file 14). An increase in genes called differentially expressed additionally alters the GO analysis results (Additional file 16 & 17). When samples were aligned to a default reference genome we sometimes missed GO pathways or misinterpreted which were the top pathways.

The choice of read aligner has long been known to give slightly differing results of differential expression due to the differences in the alignment algorithms (Conesa et al., 2016; Costa-Silva et al., 2017). Differences between HISAT and STAR could be contributed to differences in default parameters for handling multi-aligning reads (Kim et al., 2015). We show that regardless of choice of read aligner, HISAT or STAR, we observe similar results. Sample size has also long been known to alter differential expression analysis (Ching et al., 2014; Lamarre et al., 2018; Zhao et al., 2018). We therefore additionally replicated our findings in a smaller sample size of 3 male XY compared to 3 female XX samples for whole blood and brain cortex tissue and where the samples were randomly selected and confirmed the results from the larger sample size (Additional file 20).

In addition to reference-based quantification, we tested whether quantifying sex-linked reads with a pseudo-aligner would be affected by using a sex chromosome complement reference. Previous studies have shown that reference-based alignment is not necessary for high-quality estimation of transcript levels (Zielezinski et al., 2017). However, we observed expression estimates for Y-linked transcripts in female XX samples when using a default reference transcriptome index for pseudo-alignment quantification estimates. In contrast, when a sex chromosome complement informed reference transcriptome index was used, we observed no Y-linked expression in female XX samples. Salmon, and other alignment-free tools such as Kallisto (Bray et al., 2015) and Sailfish (R et al., 2014), build an index of k-mers from a reference transcriptome. The k-mer transcriptome index is used to group pseudoalignments belonging to the same set of transcripts to directly estimate the expression of each transcript. A k-mer alignment free approach is faster and less demanding than alignment protocols (Zielezinski et al., 2017); however, a sex chromosome complement informed transcriptome index should be carefully considered because even a k-mer approach is not sensitive to regions that are 100% identical in sequence. Additionally, alignment-free methods are not as robust in quantifying expression estimates for small RNAs and lowly-expressed genes (Wu et al., 2018).

The choice of reference transcriptome or reference genome can also give slightly differing results of differential expression due to the difference in which transcripts are included in the transcriptome (Zhao and Zhang, 2015). The Ensembl cDNA does not include the Y PAR linked transcripts whereas the gencode transcriptome fasta includes both the X and Y PARs. The Ensembl transcriptome does not include non-coding RNAs, such as *XIST* transcripts. The *XIST* gene is called as being up-regulated in the female XX samples for all tissues and all comparisons except for when transcript expression was estimated using the Ensembl reference transcriptome (Additional file 15, 18, & 19). Given the current builds, for RNA-seq projects interested in sex chromosome linked transcript expression, we suggest that researchers use a gencode sex chromosome complement informed reference transcriptome index.

Ideally, one would use DNA to confirm presence or absence of the Y chromosome, but if DNA sequence was not generated, one would need to confirm the genetic sex of the sample by assessing expression estimates for X-linked and Y-linked genes. To more carefully investigate the ability to use gene expression to infer sex chromosome complement of the sample, we examined the gene expression for a select set of X-Y homologous genes, as well as *XIST* and *SRY* that are known to be differentially expressed between the sexes (Figure 2, Additional file 13). The samples broadly segregated by sex for Y-linked gene expression using default alignment. However, the pattern was messy for each individual Y-linked gene. Thus, if inferring sex from RNA-Seq data, we recommend using the estimated expression of multiple X-Y homologous genes and *XIST* to infer the genetic sex of the sample. Samples should be aligned to a default reference genome first to look at the expression for several Y-specific genes to determine if the sample is XY or XX. Then samples should be realigned to the appropriate sex chromosome complement informed reference genome. Independently assessing sex chromosome complement of samples becomes increasingly important as karyotypically XY individuals are known to have lost the Y chromosome in particular tissues sampled, as shown in Alzheimer Disease (Dumanski et al., 2016), age-related macular degeneration (Grassmann et al., 2019), and in the blood of aging individuals (Forsberg, 2017), but should not have *XIST* expression. However, *XIST* may not be a sufficient marker alone to infer sex chromosome complement, especially in cancer in samples from XX individuals, where the inactive X can become reactivated (Chaligné et al., 2015). Self-reported sex may not match the sex chromosome complement of the samples, even in karyotypic individuals.

## Conclusion

Here we show that aligning RNA-Seq reads to a sex chromosome complement informed reference genome will change the results of the analysis compared to aligning reads to a default reference genome. We previously observed that a sex chromosome complement informed alignment is important for DNA as well (Webster et al., 2019). A sex chromosome complement informed approach is needed for a sensitive and specific analysis of gene expression on the sex chromosomes (Khramtsova et al., 2018). A sex chromosome complement informed reference alignment resulted in increased expression of the PARs of the X chromosome for both male XY and female XX samples. We further found different genes called as differentially expressed between the sexes and identified sex differences in gene pathways that were missed when samples were aligned to a default reference genome.

## Perspectives and Significance

The accurate alignment and pseudo-alignment of the short RNA-Seq reads to the reference genome or reference transcriptome is essential for drawing reliable conclusions from differential expression data analysis on the sex chromosomes. We strongly urge studies using RNA-Seq to carefully consider the genetic sex of the sample when quantifying reads, and provide a framework for doing so in the future (https://github.com/SexChrLab/XY_RNAseq).

## Author contributions

KCO: Supervision, Formal Analysis, Investigation, Visualization, Writing – Original Draft Preparation, Writing – Review and Editing

SMB: Formal Analysis, Investigation, Writing – Original Draft Preparation, Writing – Review and Editing

JPA: Formal Analysis, Investigation, Writing – Review and Editing

VAVV: Investigation, Writing – Review and Editing

MAW: Conceptualization, Supervision, Visualization, Resources, Project Administration,

Writing – Original Draft Preparation, Writing – Review and Editing, Funding Acquisition

## Acknowledgements

We thank Heini Natri for comments on the manuscript.

## Availability of Data and Material

The RNA-Seq datasets analyzed during the current study are available from the GTEx project through dbGaP under accession phs000424.v6.p1; we received approval to access this data under dbGaP accession #8834. All codes used are available on GitHub: https://github.com/SexChrLab/XY_RNAseq.

## Ethics Approval and Consent to Participate

Not applicable.

## Consent for Publication

Not applicable.

## Additional files

**Additional file 1. Sample IDs.** RNA-Seq whole blood, brain cortex, breast, liver, and thyroid tissue samples from 20 genetic female (46, XX) and 20 genetic male (46, XY) individuals were downloaded from the Genotype-Tissue Expression (GTEx) project (GTEx Consortium, 2015) for a total of 200 RNA-Seq tissue samples.

**Additional file 2. Histogram of sample reported age.** For each tissue, whole blood, brain cortex, breast, liver, and thyroid, male XY and female XX samples were age matched perfectly between age 55 to 70. Females are shown in blue and males are shown in lime green. Since the samples were aged perfectly the histogram bars show only the overlap of female and male samples is a mix color of the blue and lime green.

**Additional file 3. Genetic sex of RNA-Seq samples when aligned using STAR.** Gene expression log_2_(CPM+0.25/L) for select XY homologous genes (*DDX3X/Y, PCDH11X/Y, USP9X/Y, ZFX/Y, UTX/Y*) and *XIST* and *SRY* when reads were aligned to a default reference genome **A)**, and for **B)** when reads were aligned to a sex chromosome complement informed reference using STAR. Male XY whole blood, brain cortex, breast, liver, and thyroid samples are shown in blue squares and female XX in orange circles.

**Additional file 4. Genetic sex of RNA-Seq samples per tissue.** Gene expression log_2_(CPM+0.25/L) for select XY homologous genes (*DDX3X/Y, PCDH11X/Y, USP9X/Y, ZFX/Y, UTX/Y*) and *XIST* and *SRY* when reads were aligned to a default reference genome **A)**, and for **B)** when reads were aligned to a sex chromosome complement informed reference using HISAT and **C)** and **D)**, for when the reads were aligned using STAR. Male XY whole blood, brain cortex, breast, liver, and thyroid samples are shown in blue squares and female XX in orange circles.

**Additional file 5. List of samples that were removed from downstream analysis.** Samples that did not cluster with the reported sex or clustered in unexpected ways were removed from the differential expression analysis. One male XY whole blood, 4 female XX and 4 male XY brain cortex, and one female XX breast sample were removed.

**Additional file 6. Multidimensional Scaling plots.** We investigated multidimensional scaling for all shared common variable genes for dimensions 1 and 2, and for dimensions 2 and 3 in each tissue. The most variation in each tissue is explained by the aligner **C**.**aligner**. The second most variation in each tissue is explained by the sex of the sample **A.sex**.

**Additional file 7. HISAT mapped reads bar plot.** Mean difference in expression for average total reads mapped for each tissue and each sex when aligned to a sex chromosome informed versus a default reference genome. Paired t-test to test for significant difference in total reads mapped for the whole transcriptome, chromosome 8, and chromosome X. Nonparametric Wilcox single rank sum test was used to test for significant difference in total reads mapped on the Y chromosome for male samples in each tissue separately. Red * indicate a significant, p-value < 0.05, difference in average mapped reads, NS is no significant differences.

**Additional file 8. STAR mapped reads bar plot.** Mean difference in expression for average total reads mapped for each tissue and each sex when aligned to a sex chromosome informed versus a default reference genome. Paired t-test to test for significant difference in total reads mapped for the whole transcriptome, chromosome 8, and chromosome X. Nonparametric Wilcox single rank sum test was used to test for significant difference in total reads mapped on the Y chromosome for male samples in each tissue separately. Red * indicate a significant, p-value < 0.05, difference in average mapped reads, NS is no significant differences.

**Additional file 9. Paired t-test for mapped reads in default compared to sex chromosome complement reference genome.** Mean difference in expression for average total reads mapped for each tissue and each sex when aligned to a sex chromosome informed versus a default reference genome. Paired t-test to test for significant difference in total reads mapped for the whole transcriptome (WT), chromosome 8, and chromosome X. Nonparametric Wilcox single rank sum test was used to test for significant difference in total reads mapped on the Y chromosome for male samples in each tissue separately.

**Additional file 10. X chromosome expression differences between default and sex chromosome complement informed alignment.** X chromosome gene expression differences between default and sex chromosome complement informed alignment. Increase in expression when aligned to a sex chromosome complement informed reference genome is a log_2_ fold change (FC) > 0. A decrease in expression when aligned to a sex chromosome complement informed reference genome is log_2_ FC < 0. Female XX samples are indicated by red and pink circles for PAR1, XTR, PAR2 genes, and for all other X chromosome genes respectively. Blue and light blue squares represent male XY samples. Blue squares indicate which gene points are in PAR1, XTR, and PAR2, and light blue squares are for genes outside of those regions. Differences in X chromosome expression between reference genomes default and sex chromosome complement for male XY and female XX samples aligned using HISAT for the whole X chromosome and the first 5Mb are shown for the whole blood (**A** and **B**, respectively), brain cortex (**E** and **F**, respectively), breast (**I** and **J**, respectively), liver (**M** and **N**, respectively), and thyroid (**Q** and **R**, respectively). Differences in X chromosome expression between reference genomes for male XY and female XX samples aligned using STAR for the whole X chromosome and the first 5Mb are shown for the whole blood (**C** and **D**, respectively), brain cortex (**G** and **H**, respectively), breast (**K** and **L**, respectively), liver (**O** and **P**, respectively), and thyroid (**S** and **T**, respectively).

**Additional file 11. X chromosome regions mean and median expression values.** X chromosome regions PAR1, PAR2, XTR, XDG, XAR, XCR mean and median CPM expression for male XY and female XX samples for each tissue separately when aligned to a default or sex chromosome complement informed reference genome using either HISAT and STAR. Paired t-test was used to test for significant differences in expression. XTR and XAR show a significant increase, p-value < 0.05, in female expression for each tissue type. XTR and XAR additionally show a significant increase, p-value < 0.05, in male expression for liver and thyroid. PAR2 shows a significant increase, p-value < 0.05, in female liver expression. Additionally reported fold change in mean expression when using a sex chromosome complement informed compared to a default reference genome. The mean fold change in expression either increased or stayed the same ranging from 2.8 to 0.999 fold increase in expression. Finally, mean male over mean female expression was reported for each X chromosome region for each tissue. Mean male over mean female expression decreases for XTR when using a sex chromosome complement reference genome for each tissue.

**Additional file 12. Whole genome gene expression values per sample, aligner and reference genome used for alignment.** CPM values for male XY and female XX whole blood, brain cortex, breast, liver and thyroid samples when aligned to a default and sex chromosome complement informed reference genome for the whole genome (1-22, mtDNA, X, Y and non-chromosomal).

**Additional file 13. Gene expression for XY homologous genes.** X chromosome expression for 26 X and Y homologous genes (*AMELX, ARSD, ARSE, ARSF, CASK, GYG2, HSFX1, HSFX2, NLGN4X, OFD1, PCDH11X, PRKX, RBMX, RPS4X, SOX3, STS, TBL1X, TGIF2LX, TMSB4X, TSPYL2, USP9X, VCX, VCX2, VCX3A, VCX3B, ZFX)*. Difference in gene expression for when male XY and female XX samples were aligned to a default and sex chromosome complement informed reference genome for each tissue. Little to no difference in gene expression between default and sex chromosome complement informed reference genome alignment was observed for 25 of the 26 X and Y homologous genes for both male XY and female XX samples using either HISAT or STAR. The log_2_ fold increase in expression for *PCDH11X* when aligned using HISAT was 0.4, 0.28, 0.33, 0.16, and 0.16 for whole blood, brain cortex, breast, liver, and thyroid, respectively. The greatest increase in expression was observed for *PCDH11X* in female whole blood at a log_2_ fold increase of 0.4.

**Additional file 14. Differentially expressed genes between the sexes that were uniquely and jointly called between reference genomes.** Genes that are differentially expressed between the sexes, male XY and female XX, for whole blood, brain cortex, breast, liver, and thyroid samples. Differentially expressed genes that are uniquely called when using either the default or sex chromosome complement informed reference genome and differentially expressed genes that were jointly called between the reference genomes.

**Additional file 15. Gene expression differences between male XY and female XX samples.** Sex differences in gene expression for whole blood, brain cortex, breast, liver, and thyroid samples for when samples were aligned to a default reference genome and to a reference genome informed on the sex chromosome complement. Showing sex differences in gene expression between reference genomes used for alignment and for when samples were aligned using HISAT and STAR.

**Additional file 16. GO analysis of differentially expressed genes in female and male samples with HISAT aligner.** Gene enrichment analysis of genes that are more highly expressed in one sex versus the other sex for each tissue, whole blood, brain cortex, breast, liver and thyroid, when samples were aligned to a default or sex chromosome complement informed reference genome using HISAT.

**Additional file 17. GO analysis of differentially expressed genes in female and male samples with STAR aligner.** Gene enrichment analysis of genes that are more highly expressed in one sex versus the other sex for each tissue, whole blood, brain cortex, breast, liver and thyroid, when samples were aligned to a default or sex chromosome complement informed reference genome using STAR.

**Additional file 18. Sex chromosome complement informed transcriptome reference eliminates Y-linked expression in female XX samples. A)** Sex differences in gene expression, log_2_(CPM+0.25/L), between the sixteen samples from genetic males and females are shown when aligning all samples to the default Ensembl reference transcriptome (left) and a reference transcriptome informed on the sex chromosome complement (right) for brain cortex. Each point represents a gene. Genes that are differentially expressed, adjusted p-value < 0.01 are indicated in black for autosomal genes, blue for Y-linked genes, and red for X-linked genes. **B)** We show overlap between genes that are called as differentially expressed when all samples are pseudo-aligned to the default transcriptome, and genes that are called as differentially expressed when pseudo-aligned to a sex chromosome complement informed transcriptome reference. When samples were aligned to a reference transcriptome informed on the sex chromosome complement, 14 genes were called as differentially expressed between the sexes. *PLCXD1* was uniquely called as differentially expressed when aligned to a default reference genome.

**Additional file 18. Ensembl sex chromosome complement informed transcriptome reference eliminates Y-linked expression in female XX samples. A)** Sex differences in gene expression, log_2_(CPM+0.25/L), between the sixteen samples from genetic males and females are shown when aligning all samples to the default Ensembl reference transcriptome (left) and a reference transcriptome informed on the sex chromosome complement (right) for brain cortex. Each point represents a gene. Genes that are differentially expressed, adjusted p-value < 0.01 are indicated in black for autosomal genes, blue for Y-linked genes, and red for X-linked genes. **B)** We show overlap between genes that are called as differentially expressed when all samples are pseudo-aligned to the default transcriptome, and genes that are called as differentially expressed when pseudo-aligned to a sex chromosome complement informed transcriptome reference. When samples were aligned to a reference transcriptome informed on the sex chromosome complement, 14 genes were called as differentially expressed between the sexes. *PLCXD1* was uniquely called as differentially expressed when aligned to a default reference genome.

**Additional file 19. Gencode sex chromosome complement informed transcriptome reference eliminates Y-linked expression in female XX samples. A)** Sex differences in gene expression, log_2_(CPM+0.25/L), between the sixteen samples from genetic males and females are shown when aligning all samples to the default gencode reference transcriptome (left) and a reference transcriptome informed on the sex chromosome complement (right) for brain cortex. Each point represents a gene. Genes that are differentially expressed, adjusted p-value < 0.01 are indicated in black for autosomal genes, blue for Y-linked genes, and red for X-linked genes. **B)** We show overlap between genes that are called as differentially expressed when all samples are pseudo-aligned to the default transcriptome, and genes that are called as differentially expressed when pseudo-aligned to a sex chromosome complement informed transcriptome reference. When samples were aligned to a reference transcriptome informed on the sex chromosome complement, 17 genes were called as differentially expressed between the sexes. *ZBED1* was uniquely called as differentially expressed when aligned to a default reference genome.

**Additional file 20. 3 male XY and 3 female XX brain cortex and whole blood differential expression analysis.** Replicated analysis in a smaller sample size of 3 male XY compared to 3 female XX samples for whole blood and brain cortex tissue. Samples were randomly selected, and confirm the results from the larger sample size.

