## Supplementary figures and images for "Reference genome and transcriptome informed by the sex chromosome complement of the sample increases ability to detect sex differences in gene expression from RNA-Seq data"

### Additional file 2

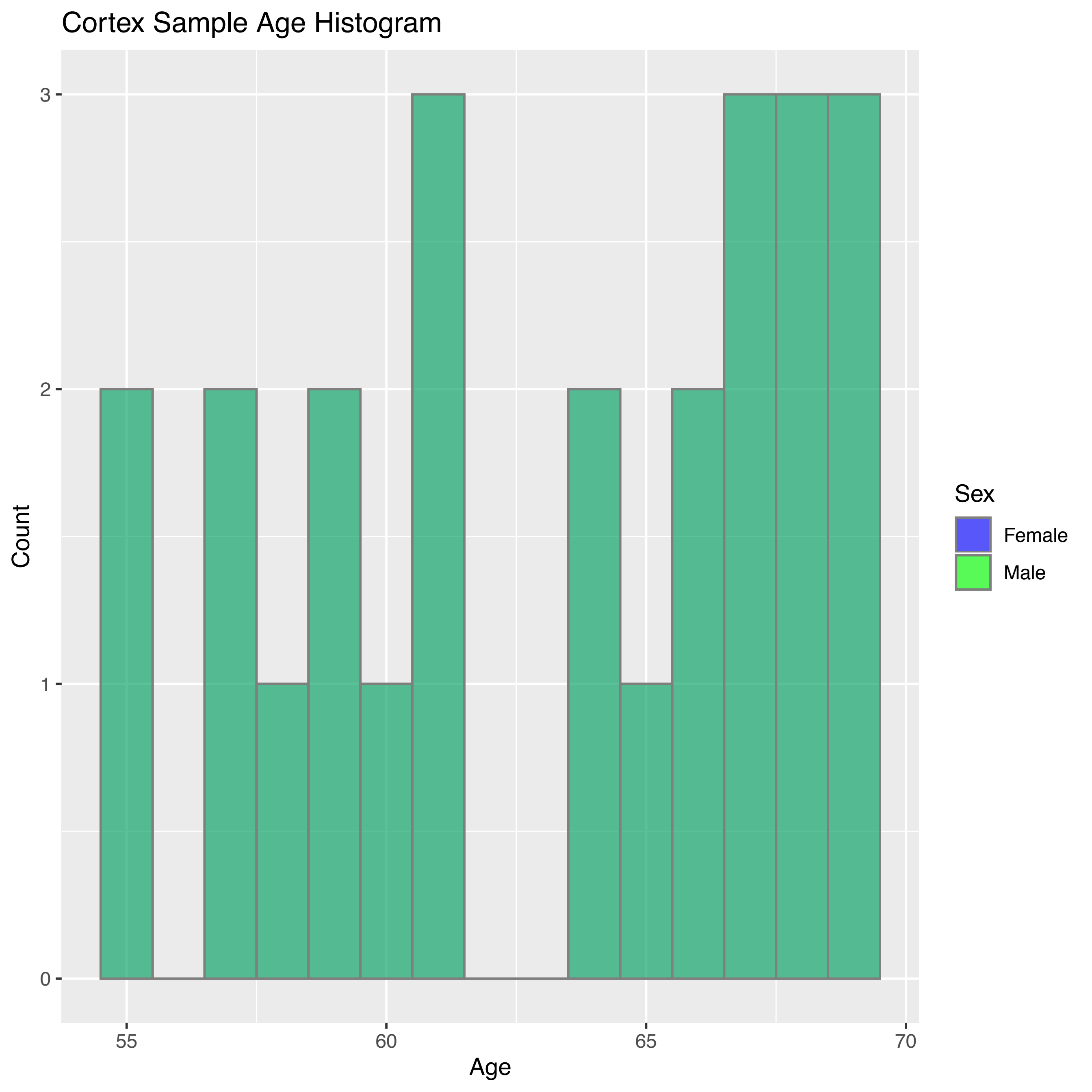

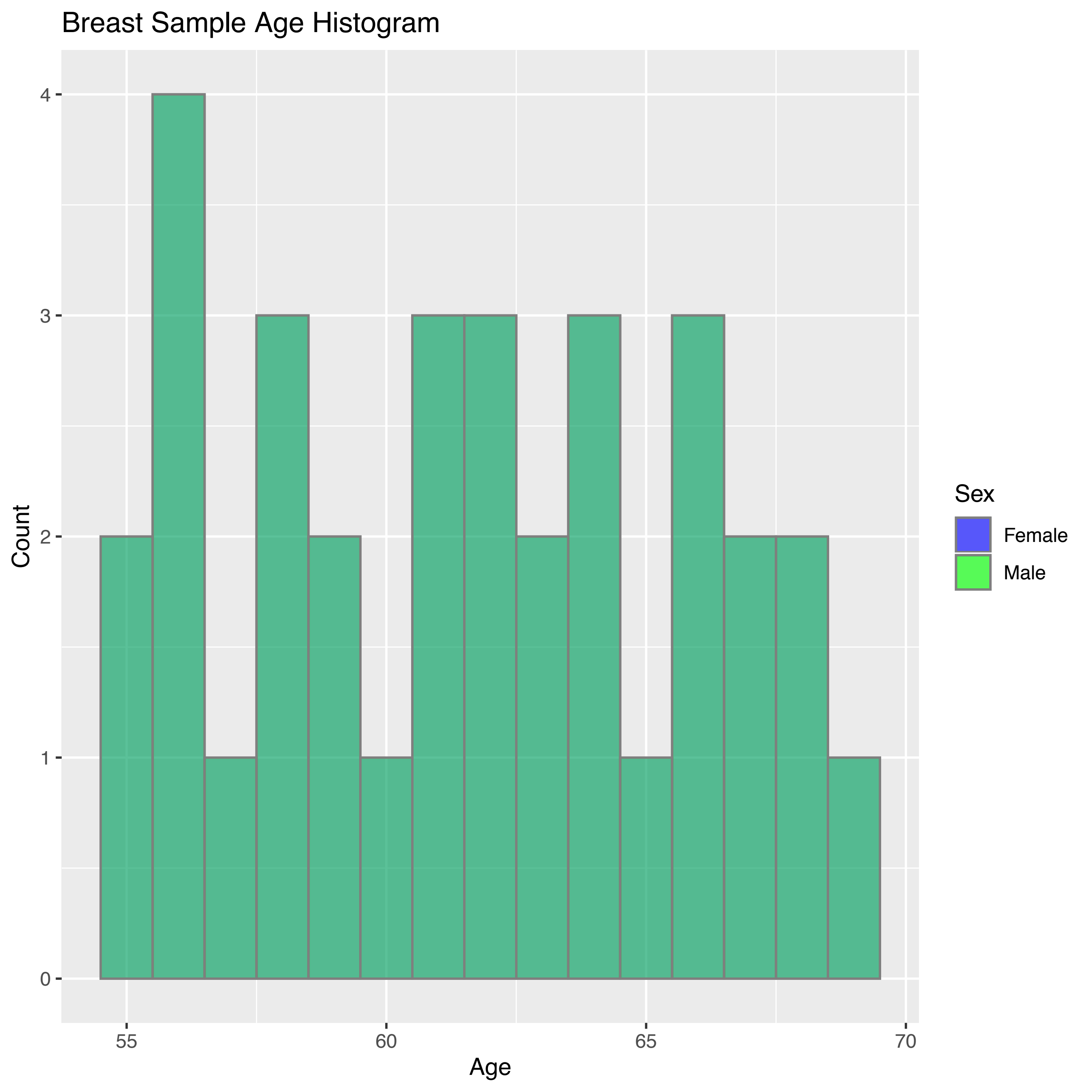

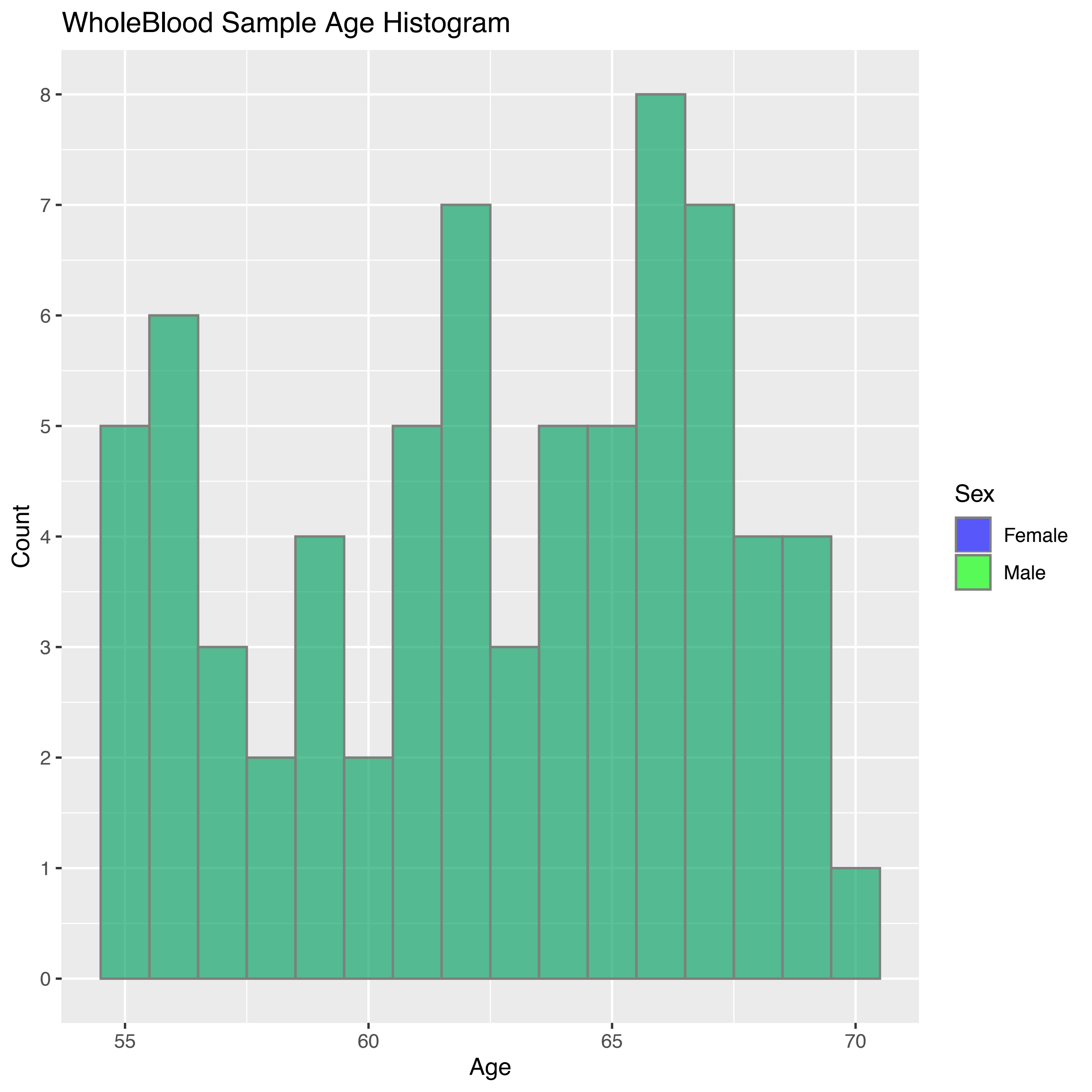

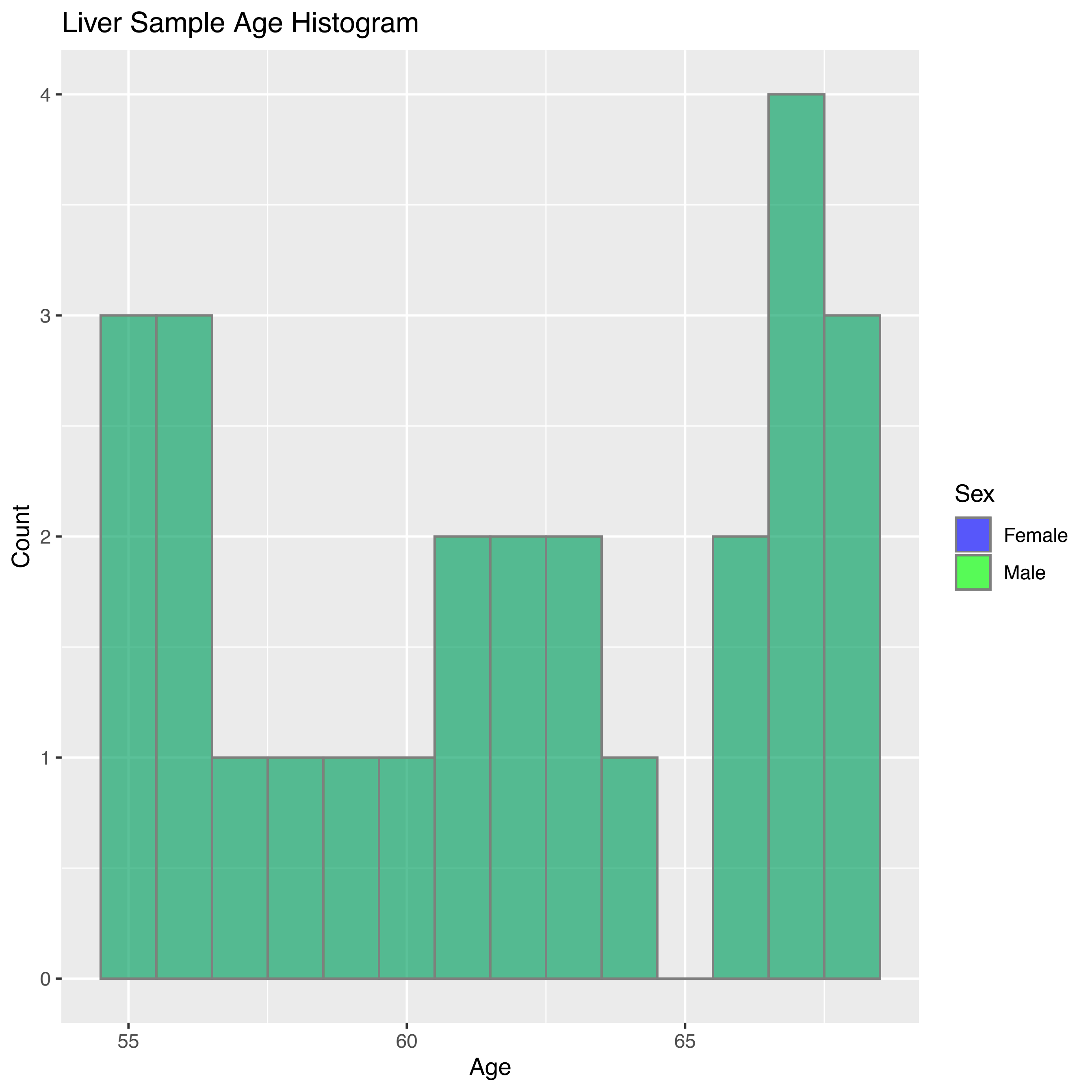

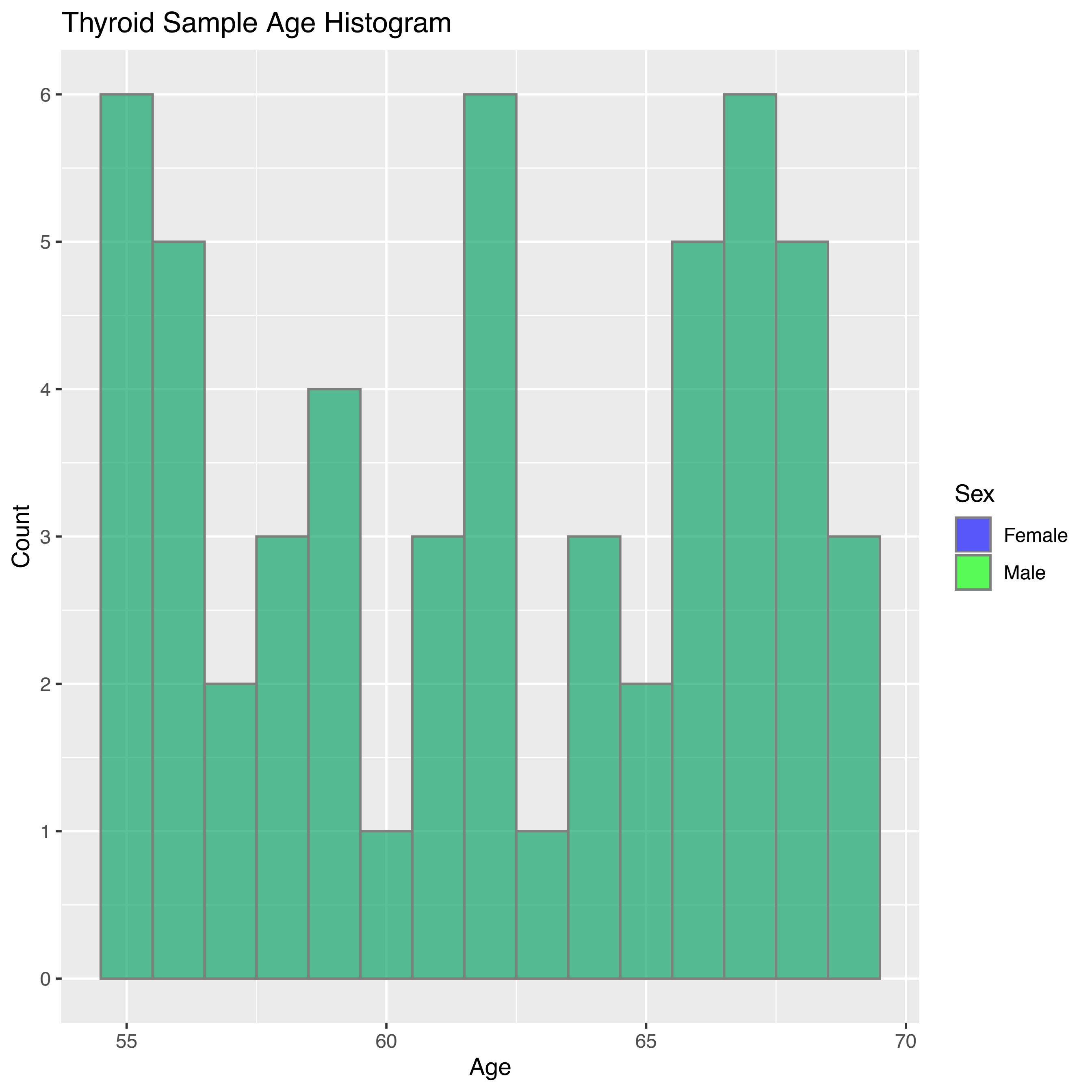

### Additional file 7

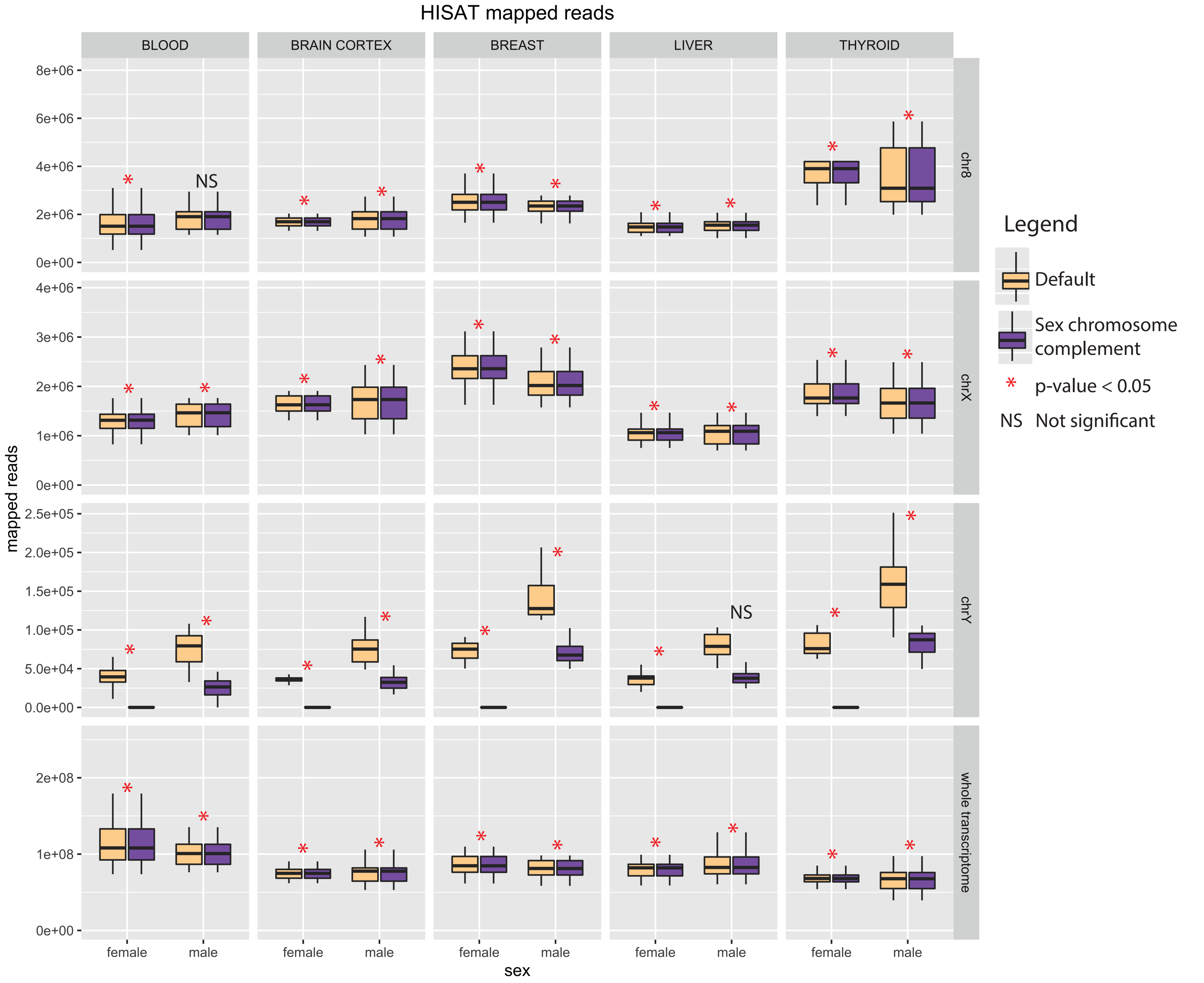

### Additional file 8

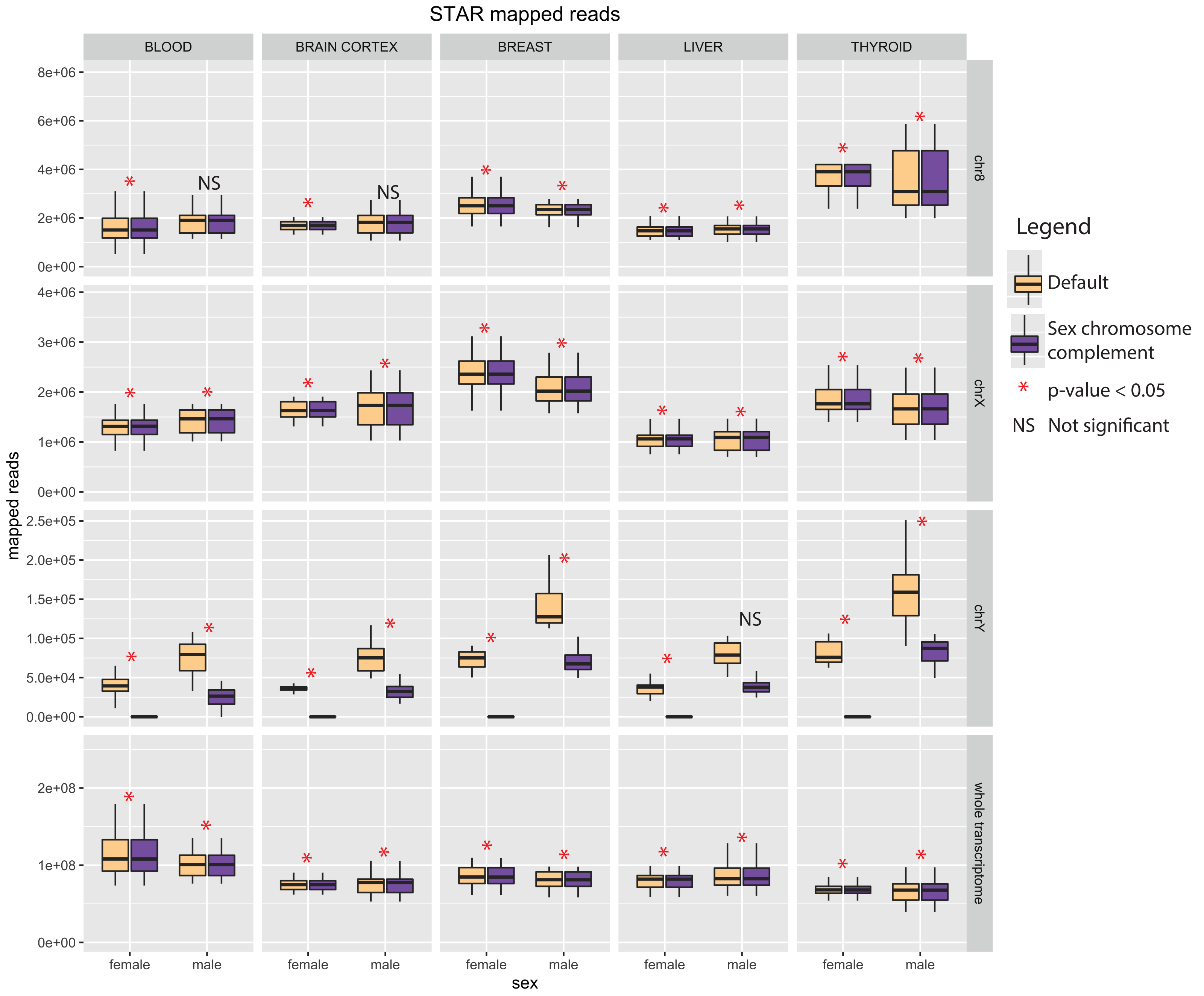
