## Additional file 3 for "Reference genome and transcriptome informed by the sex chromosome complement of the sample increases ability to detect sex differences in gene expression from RNA-Seq data"

A

All TISSUES aligned to STAR and default reference genome

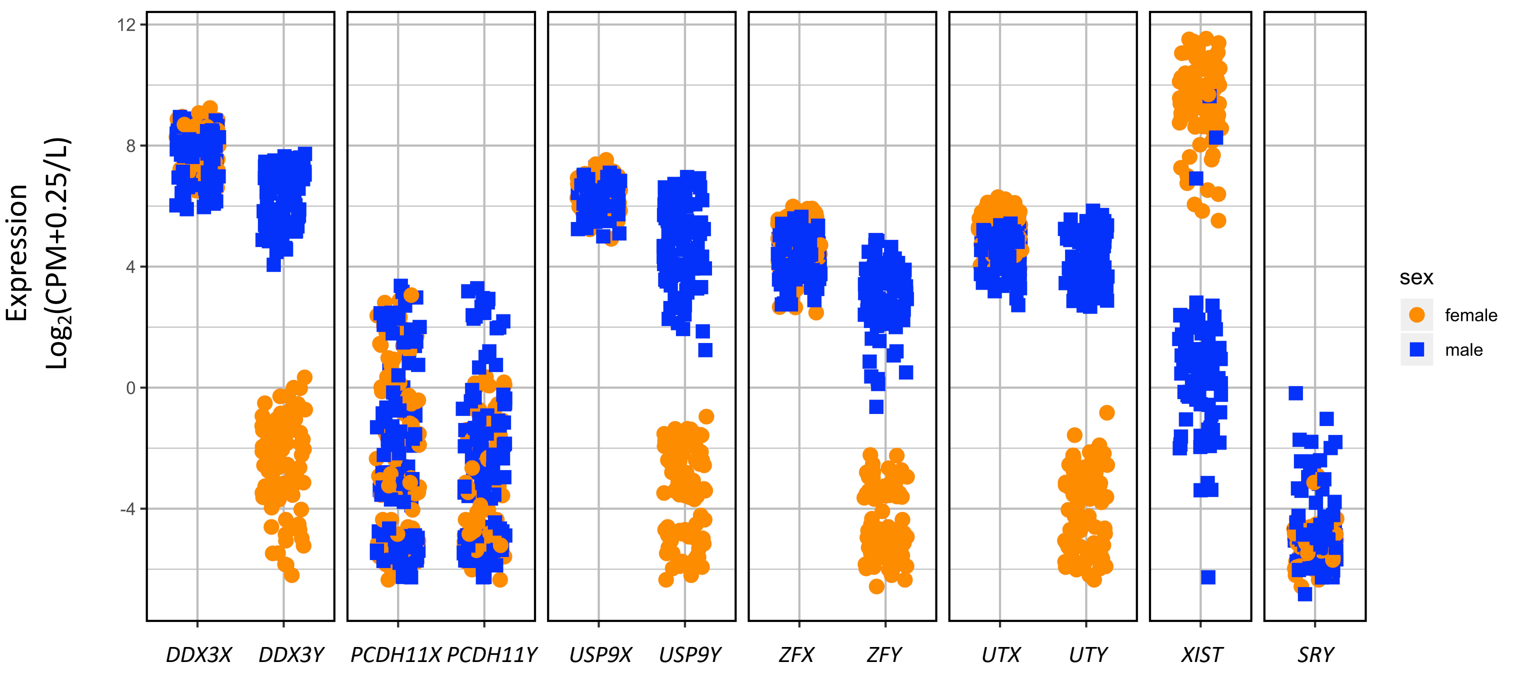


B

All TISSUES aligned to STAR and sex chromosome complement reference genome

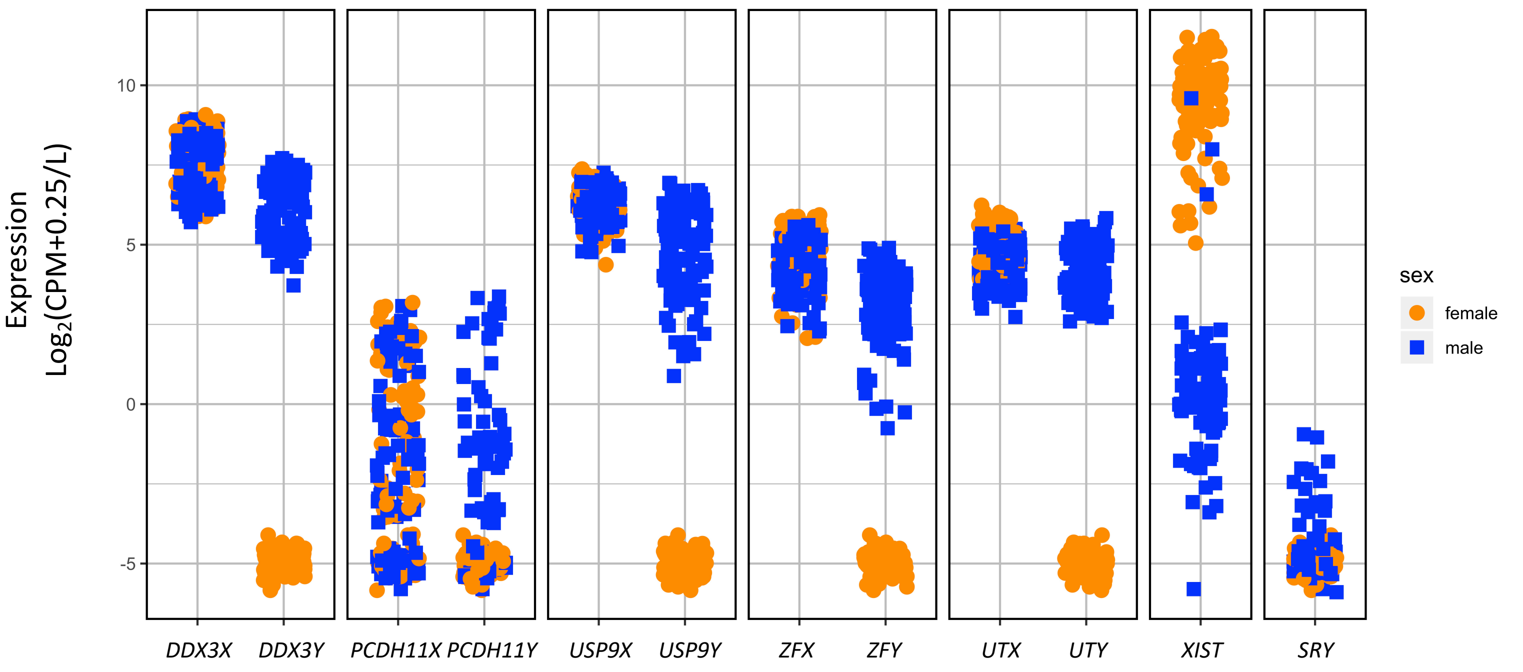
