## Additional file 4 for "Reference genome and transcriptome informed by the sex chromosome complement of the sample increases ability to detect sex differences in gene expression from RNA-Seq data"

A BLOOD aligned to HISAT and default reference genome

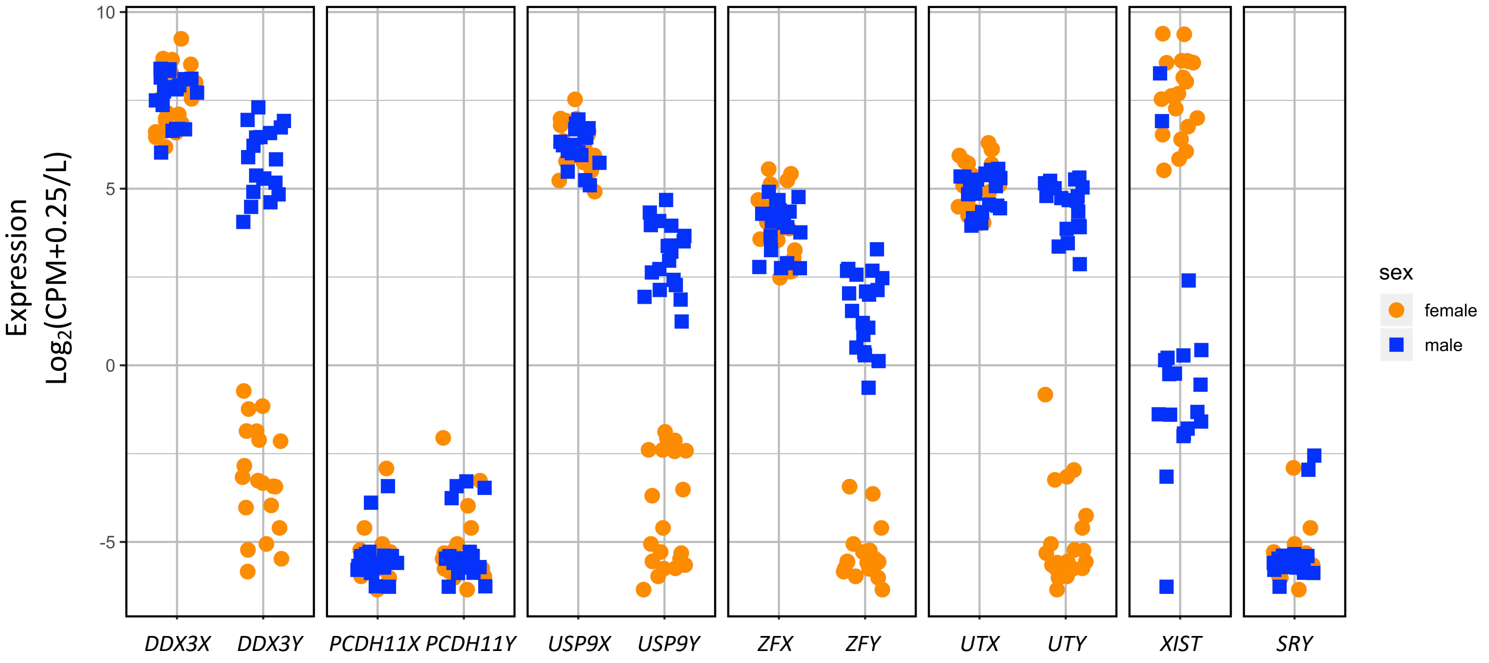


B BLOOD aligned to HISAT and sex chromosome complement reference genome

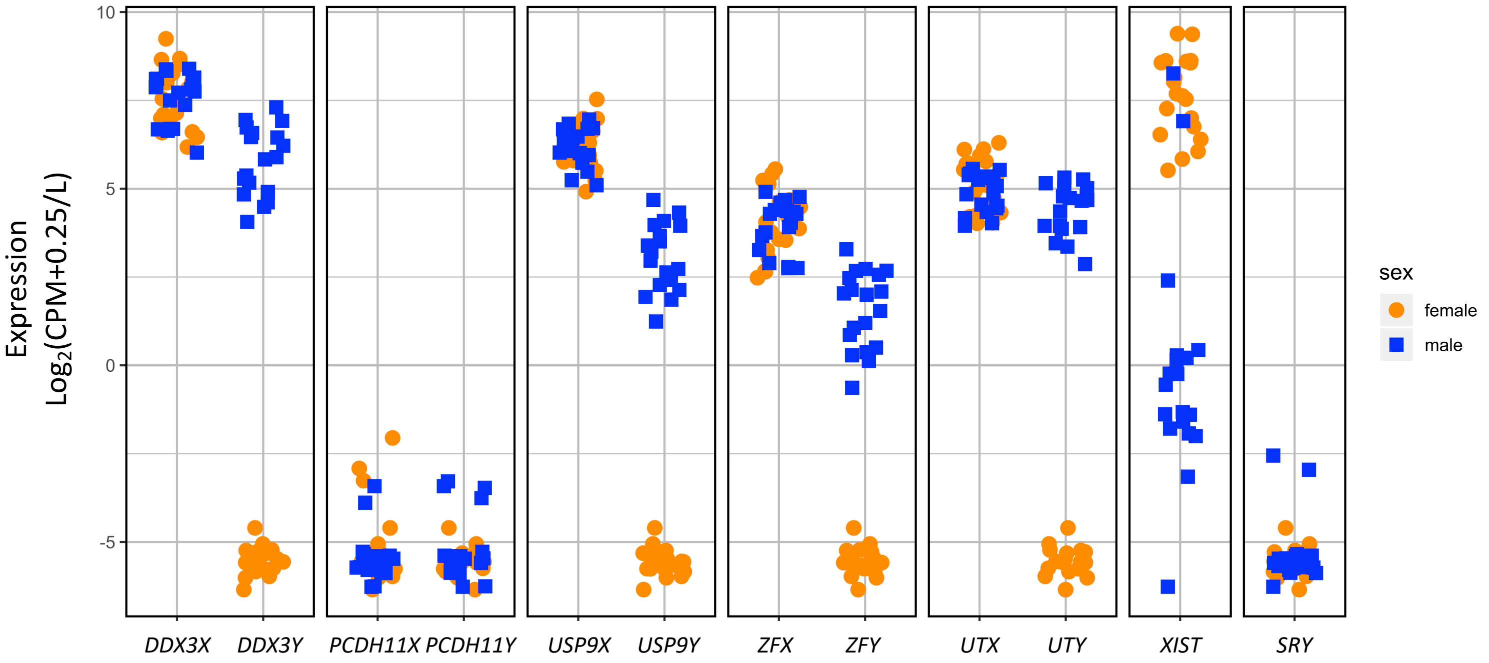

C BLOOD aligned to STAR and default reference genome

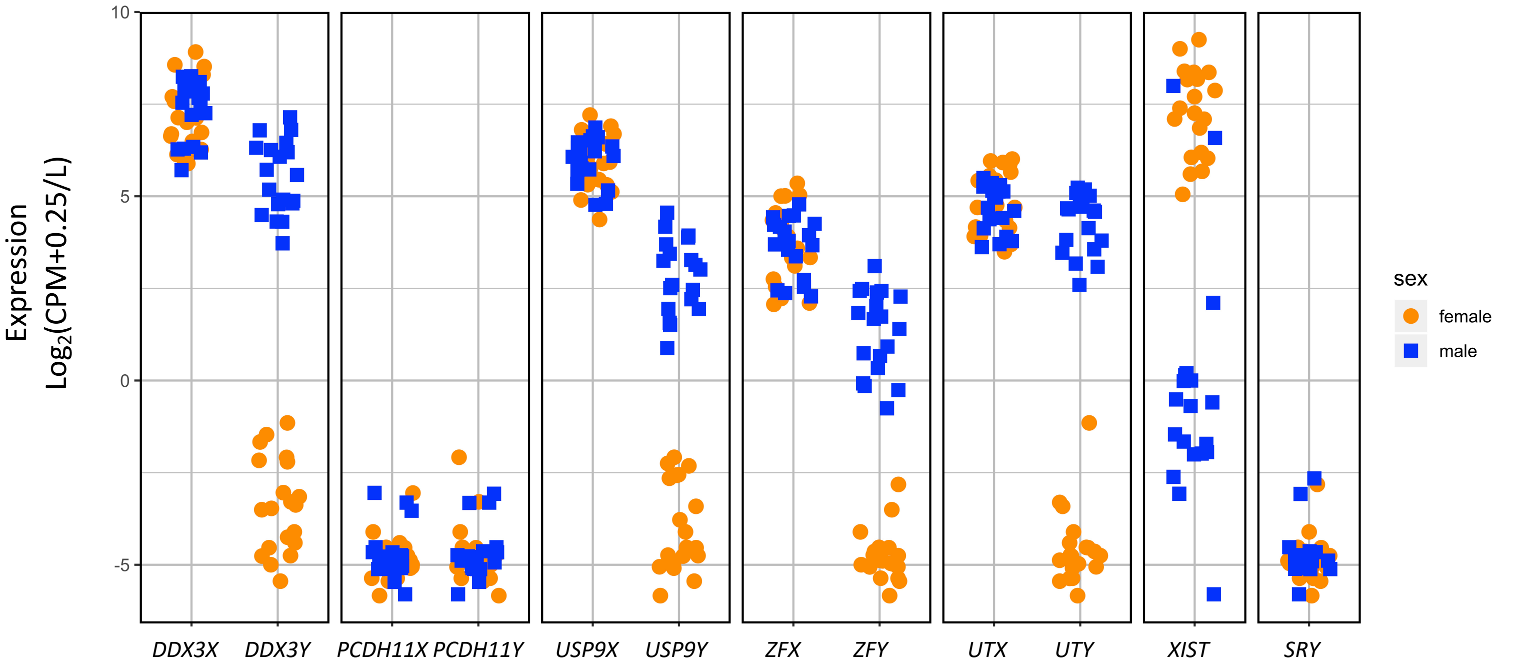


D BLOOD aligned to STAR and sex chromosome complement reference genome

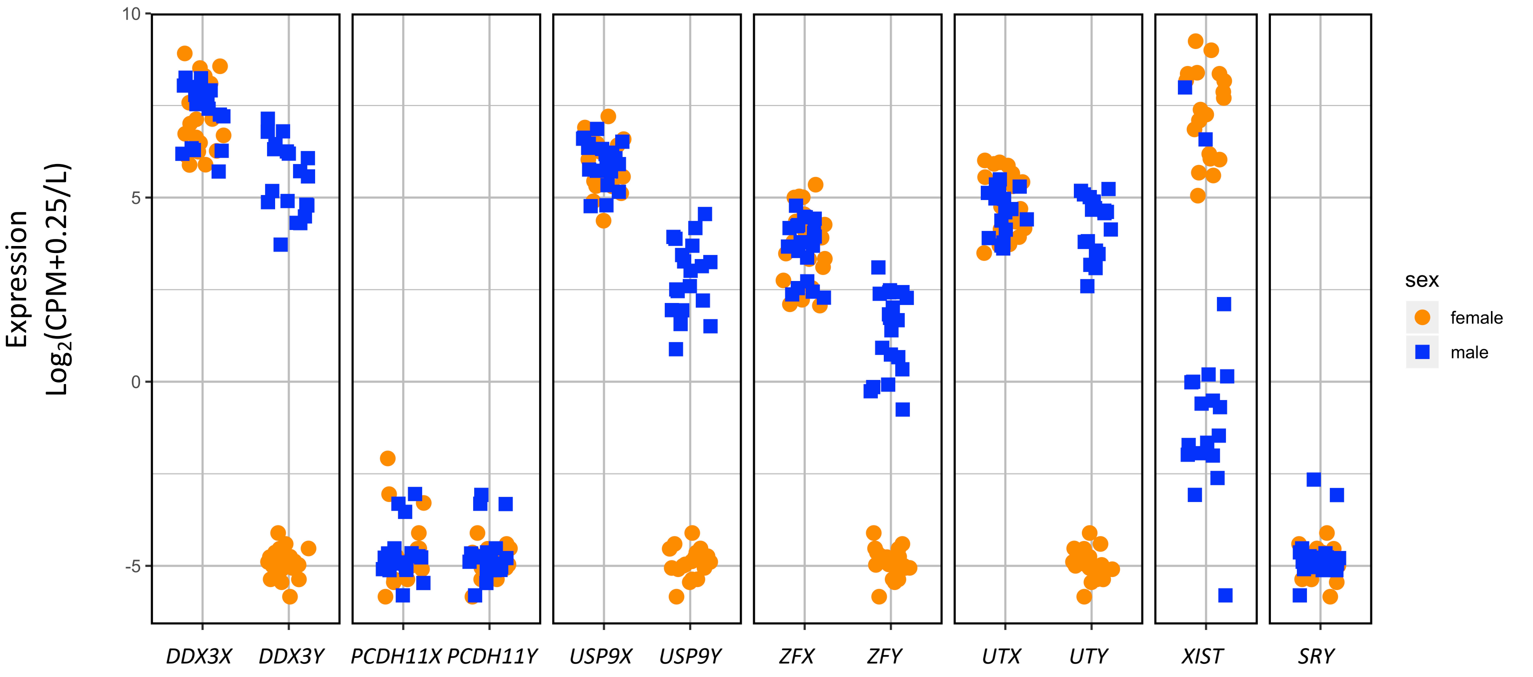

E BRAIN CORTEX aligned to HISAT and default reference genome

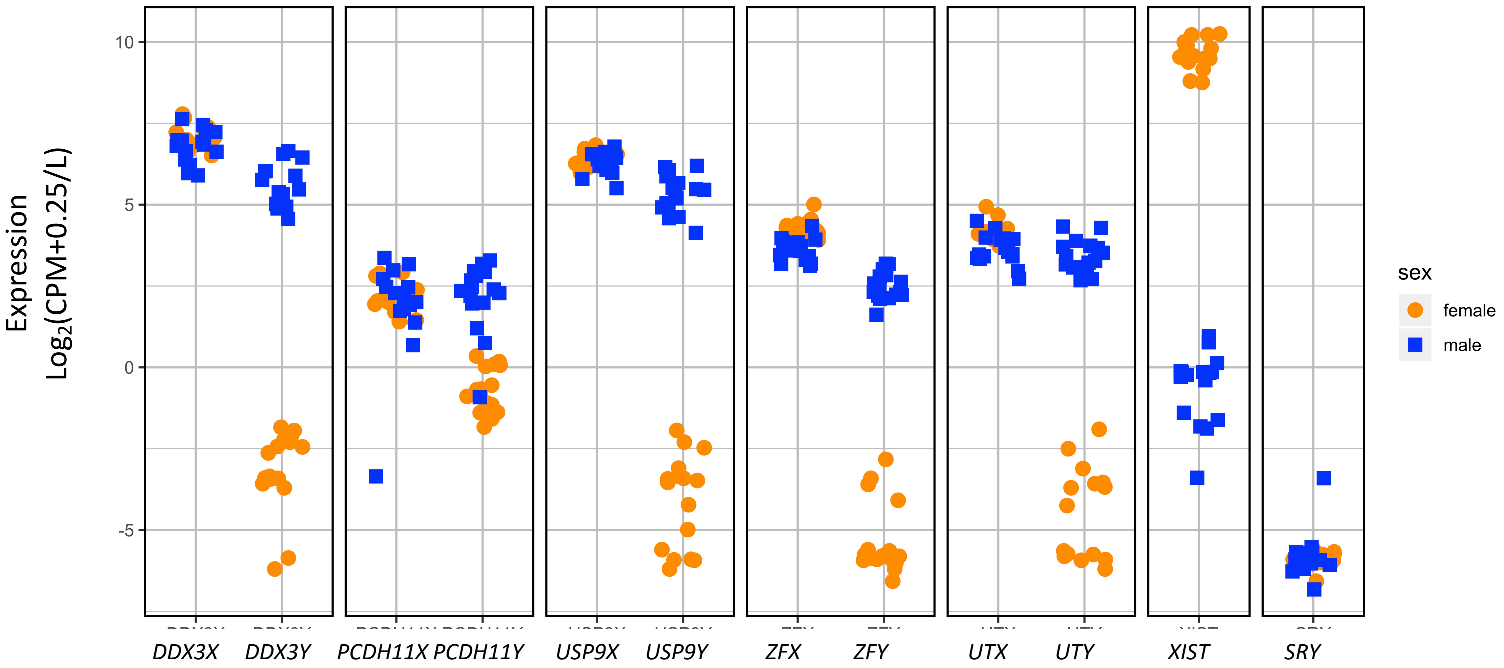


F BRAIN CORTEX aligned to HISAT and sex chromosome complement reference genome

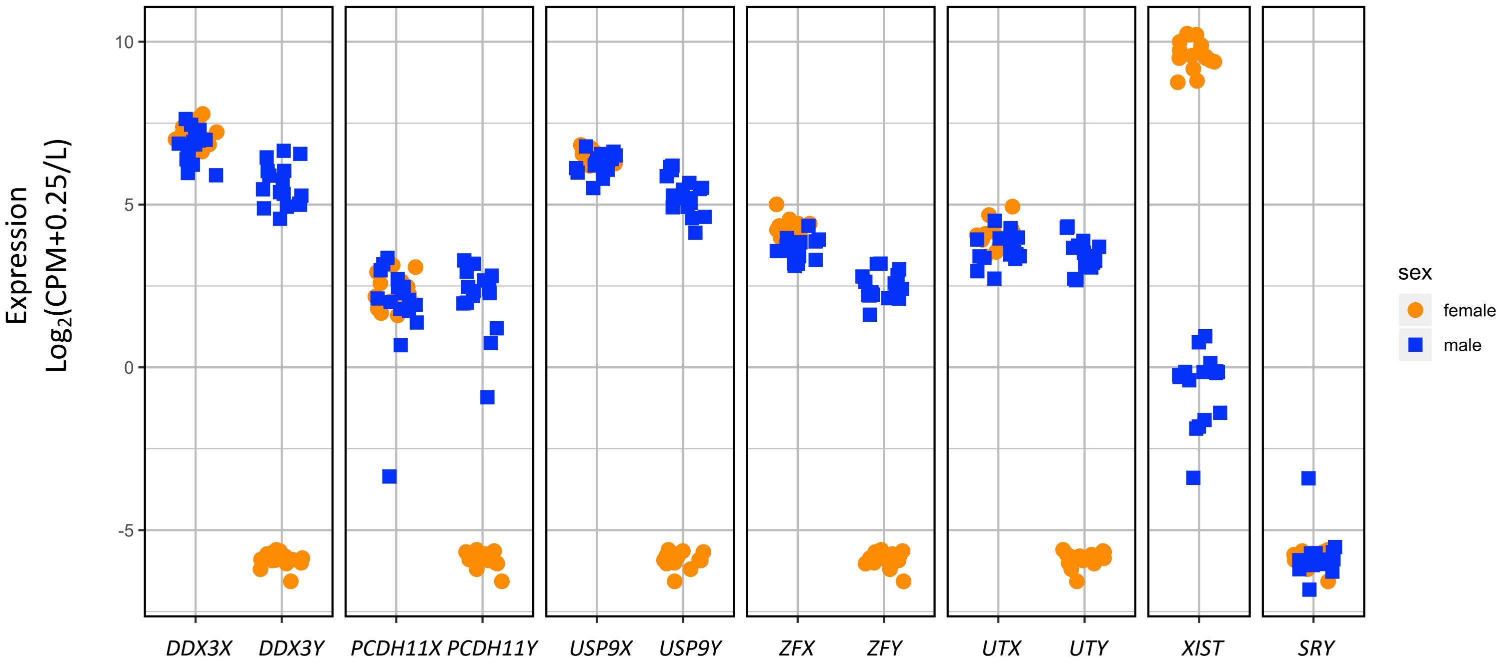

G BRAIN CORTEX aligned to STAR and default reference genome

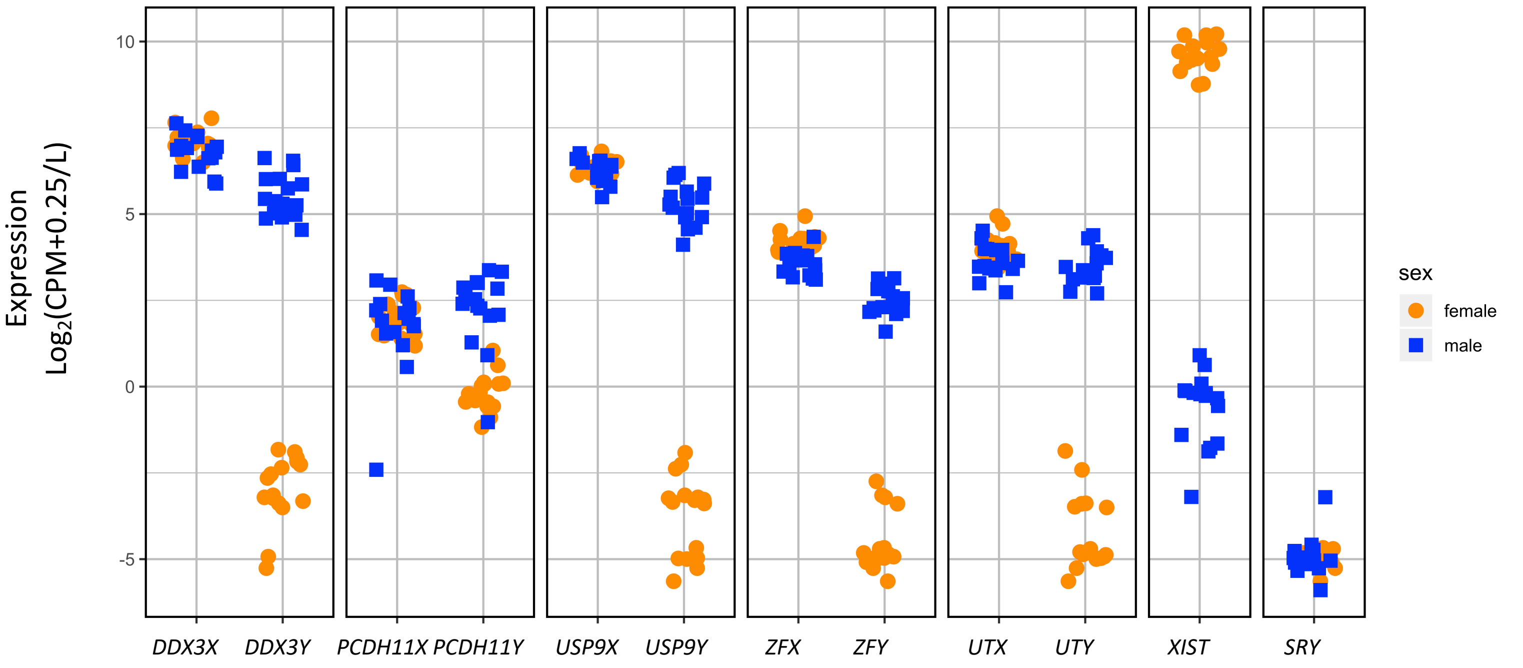


H BRAIN CORTEX aligned to STAR and sex chromosome complement reference genome

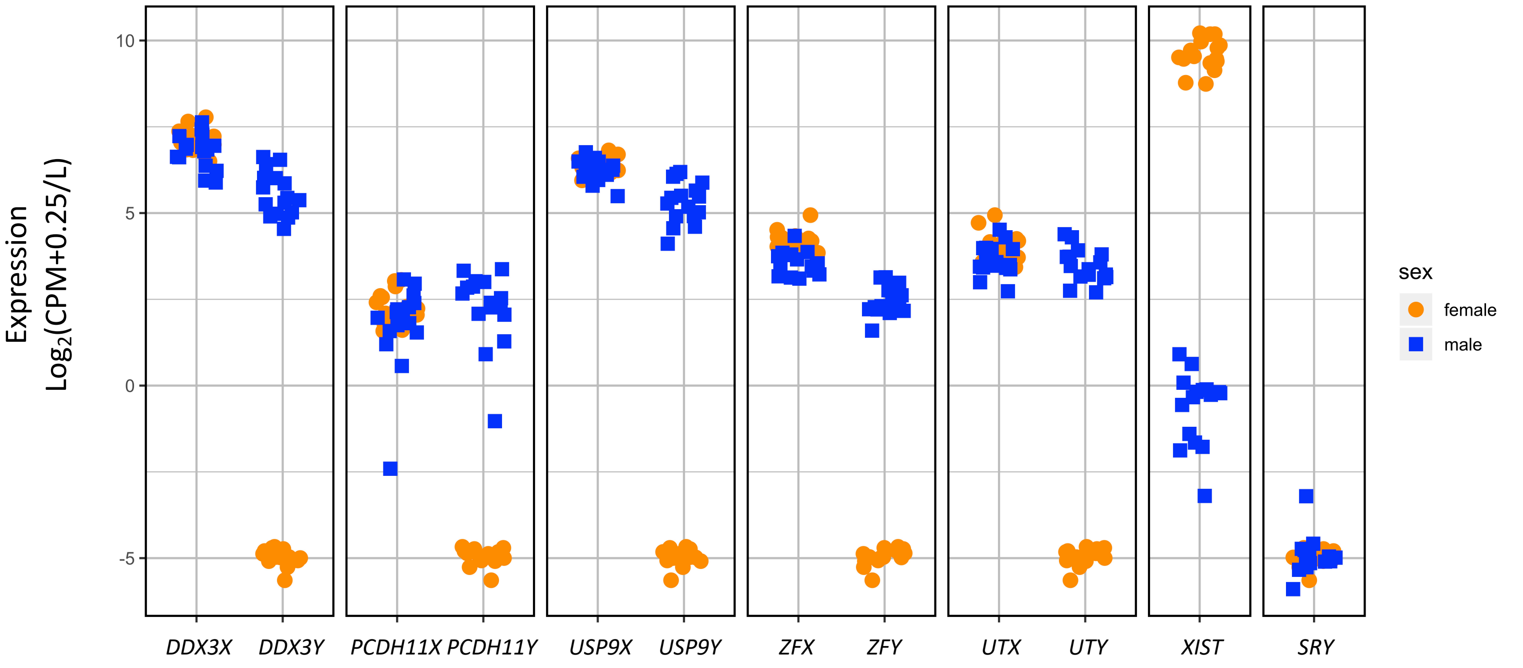

I

BREAST aligned to HISAT and default reference genome

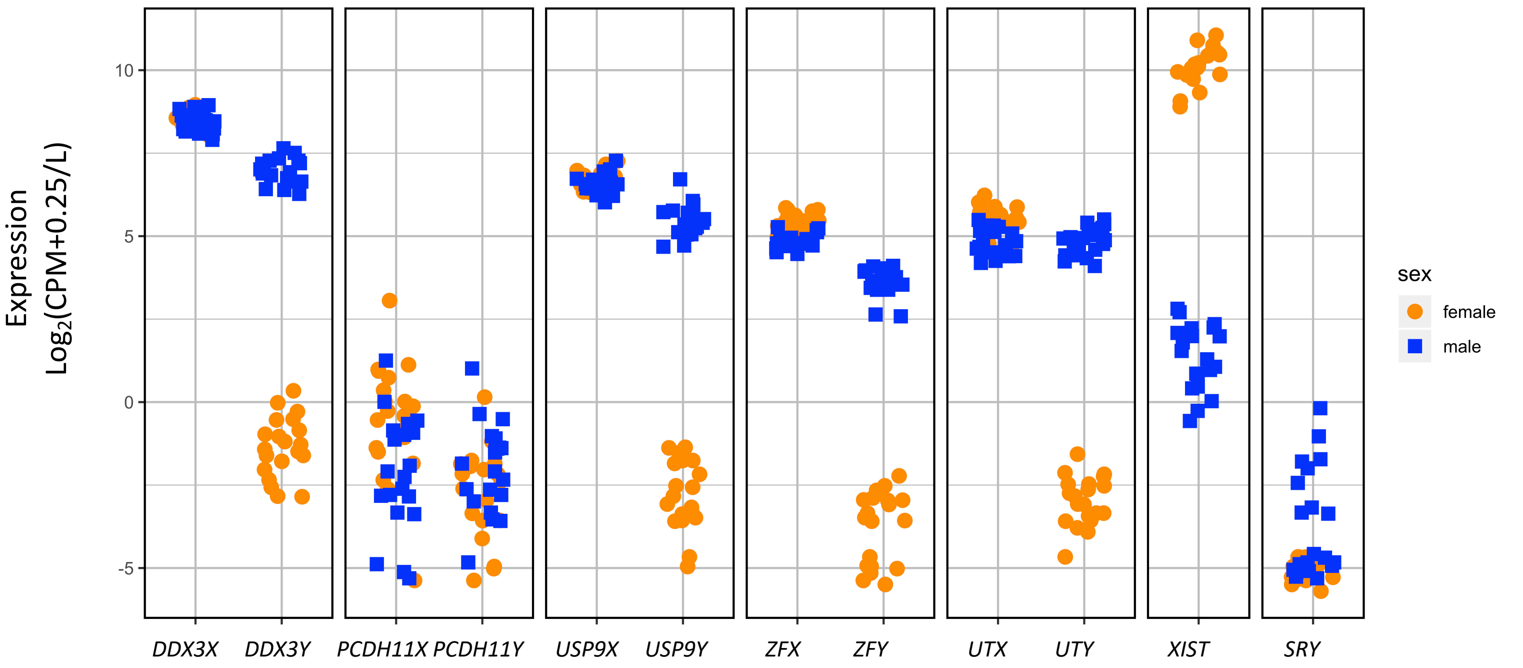


J

BREAST aligned to HISAT and sex chromosome complement reference genome

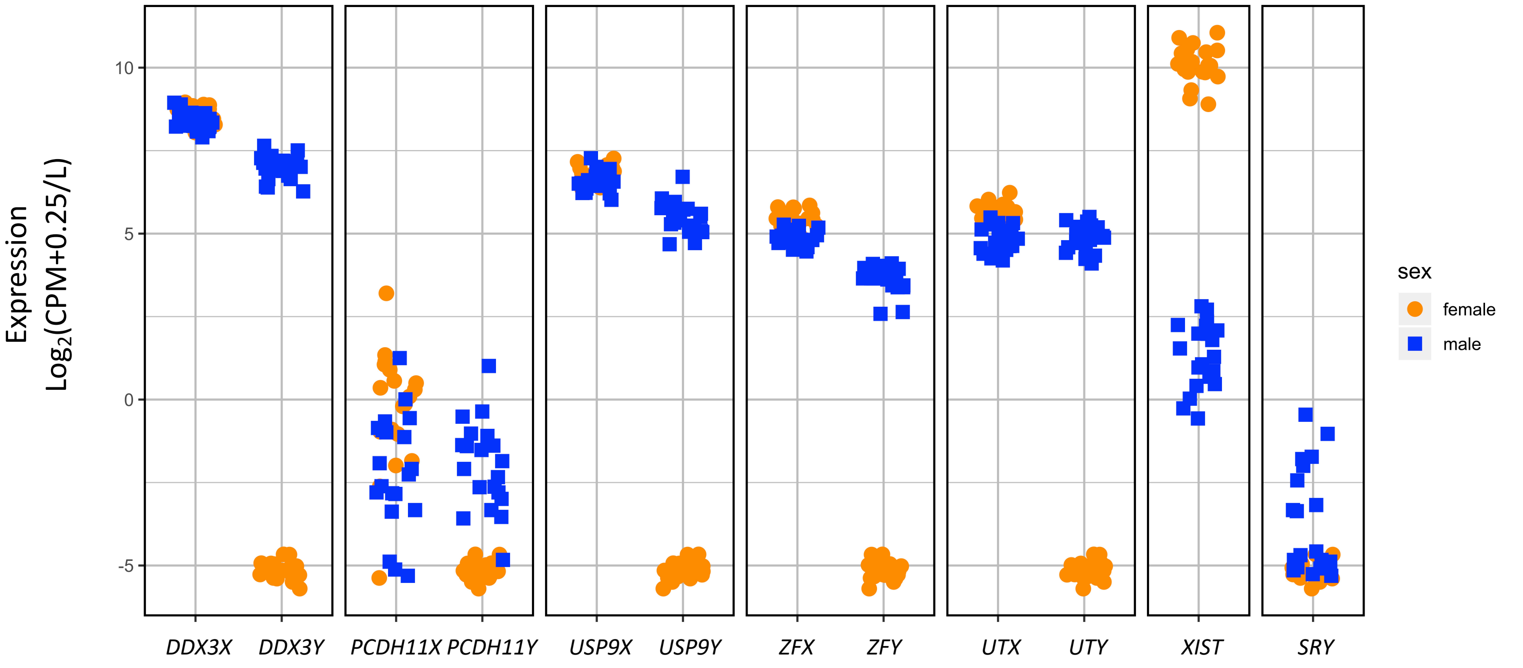

K

BREAST aligned to STAR and default reference genome

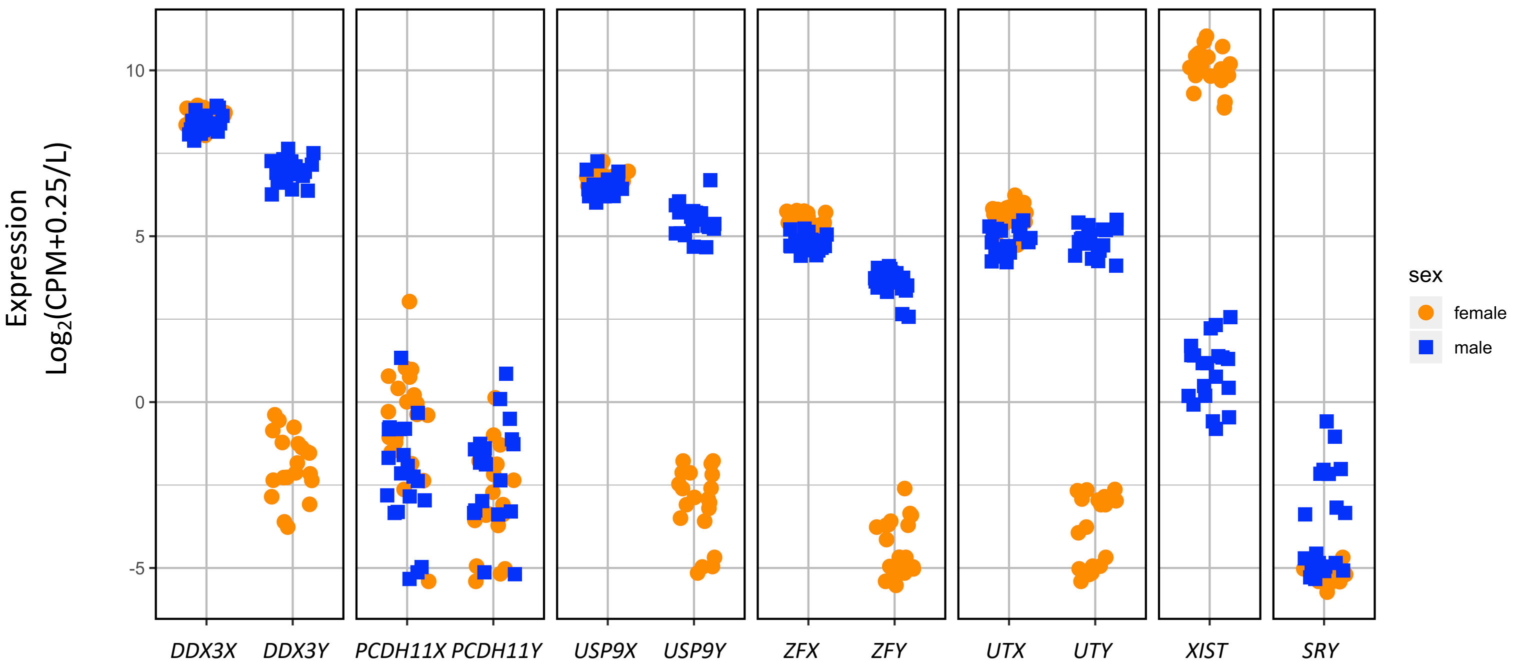


L

BREAST aligned to STAR and sex chromosome complement reference genome

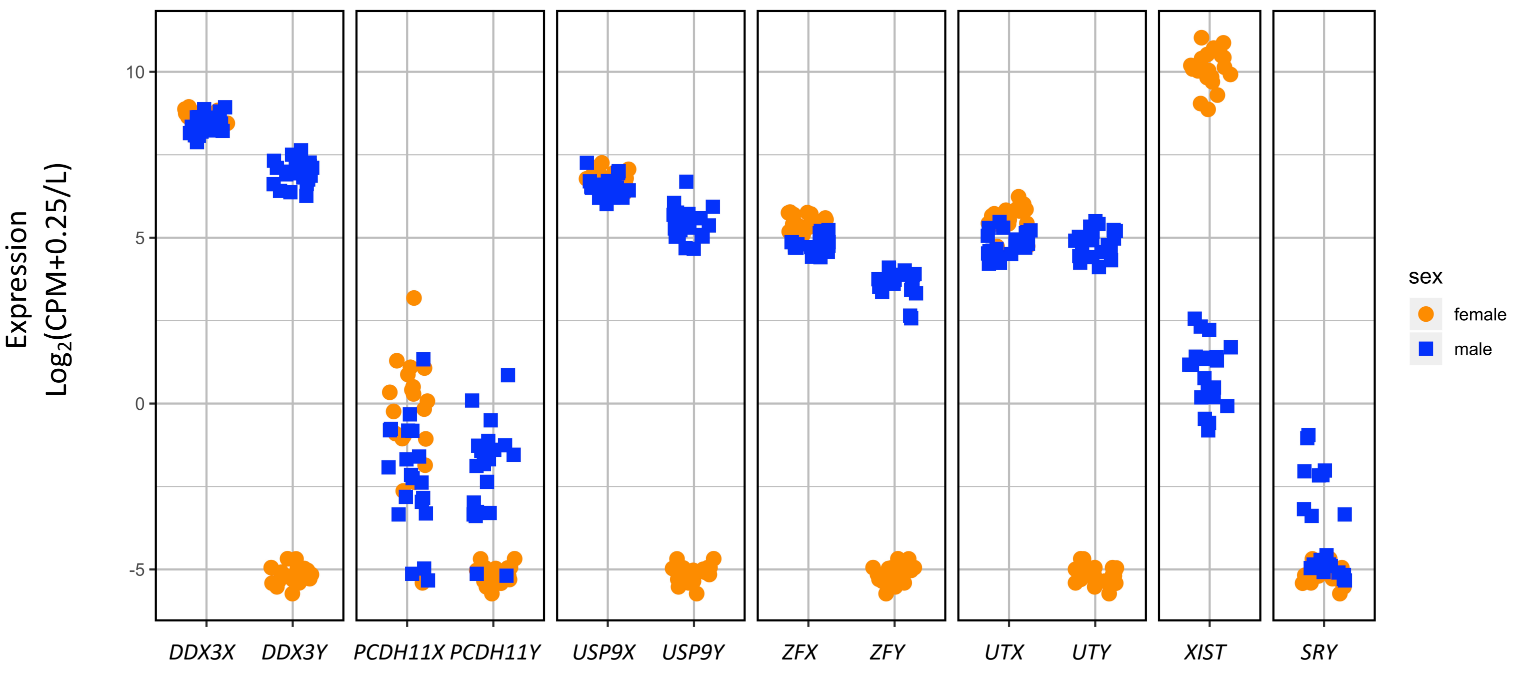

M LIVER aligned to HISAT and default reference genome

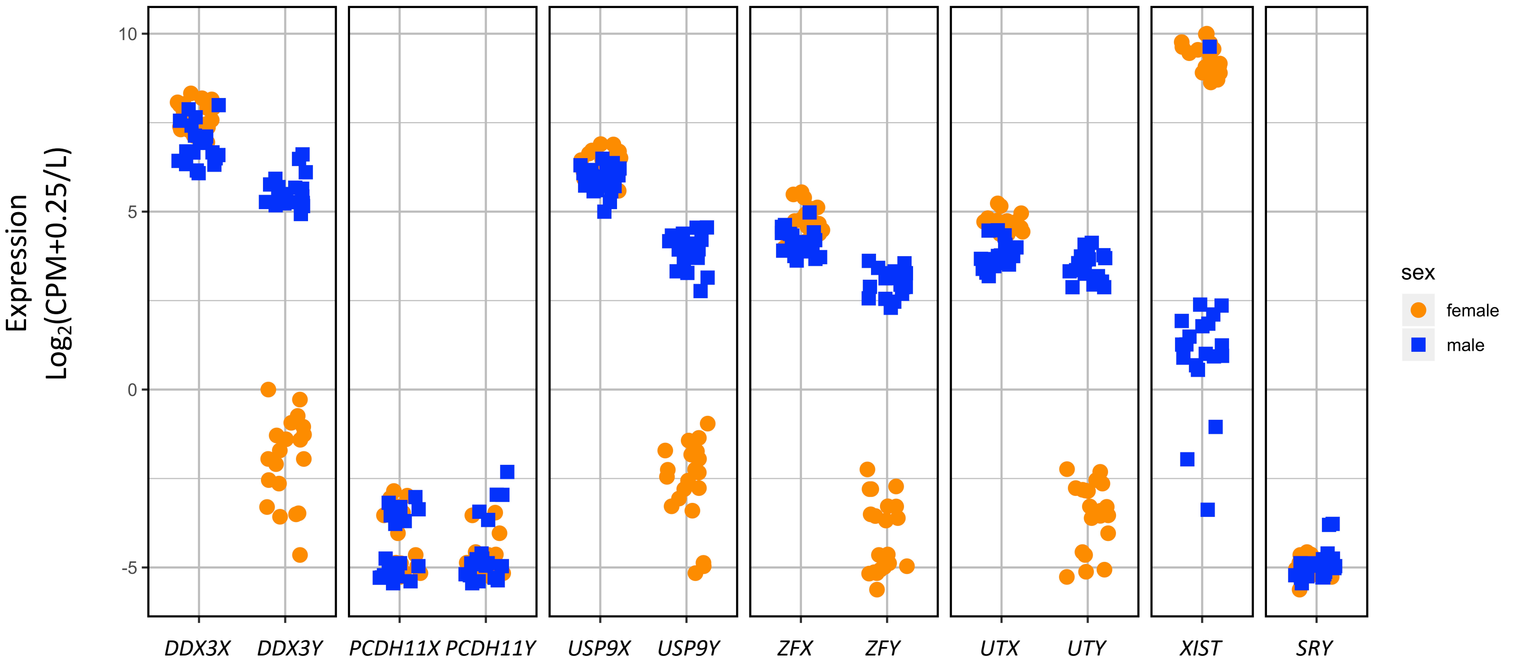


N LIVER aligned to HISAT and sex chromosome complement reference genome

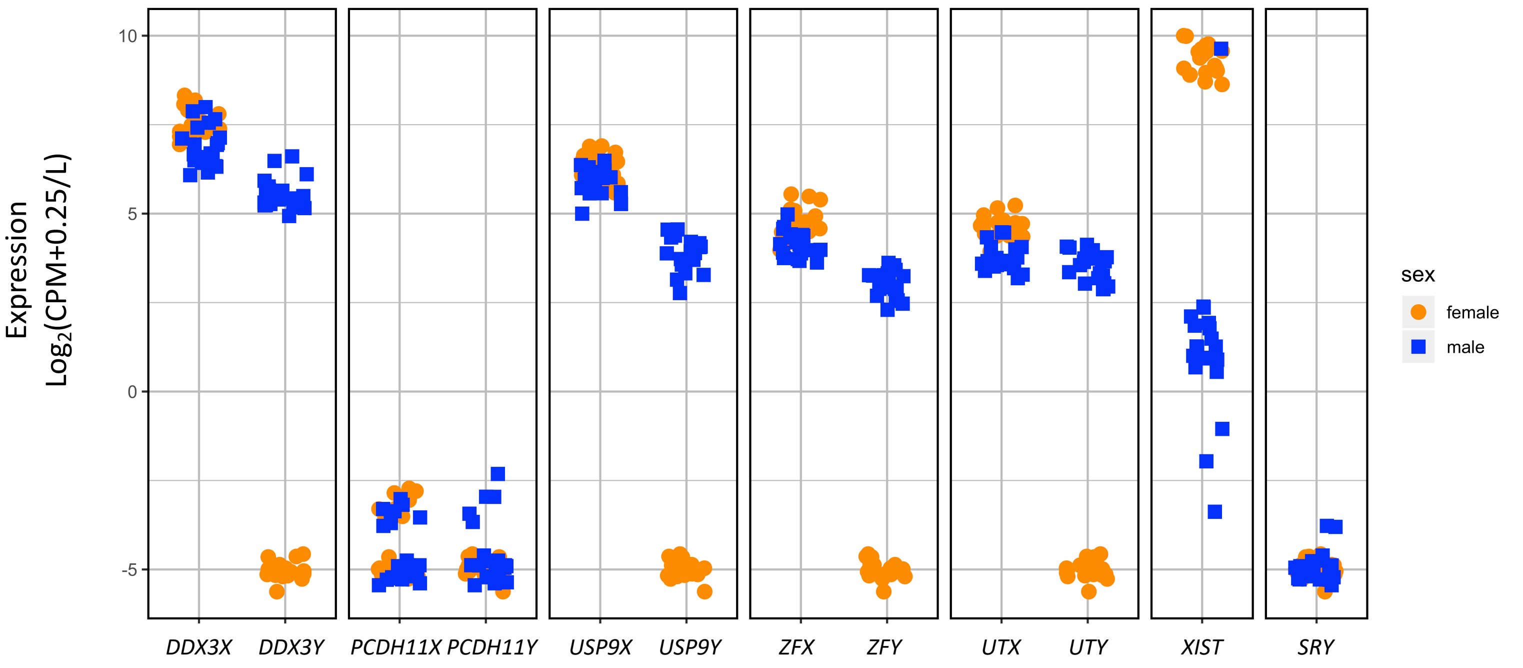

O LIVER aligned to STAR and default reference genome

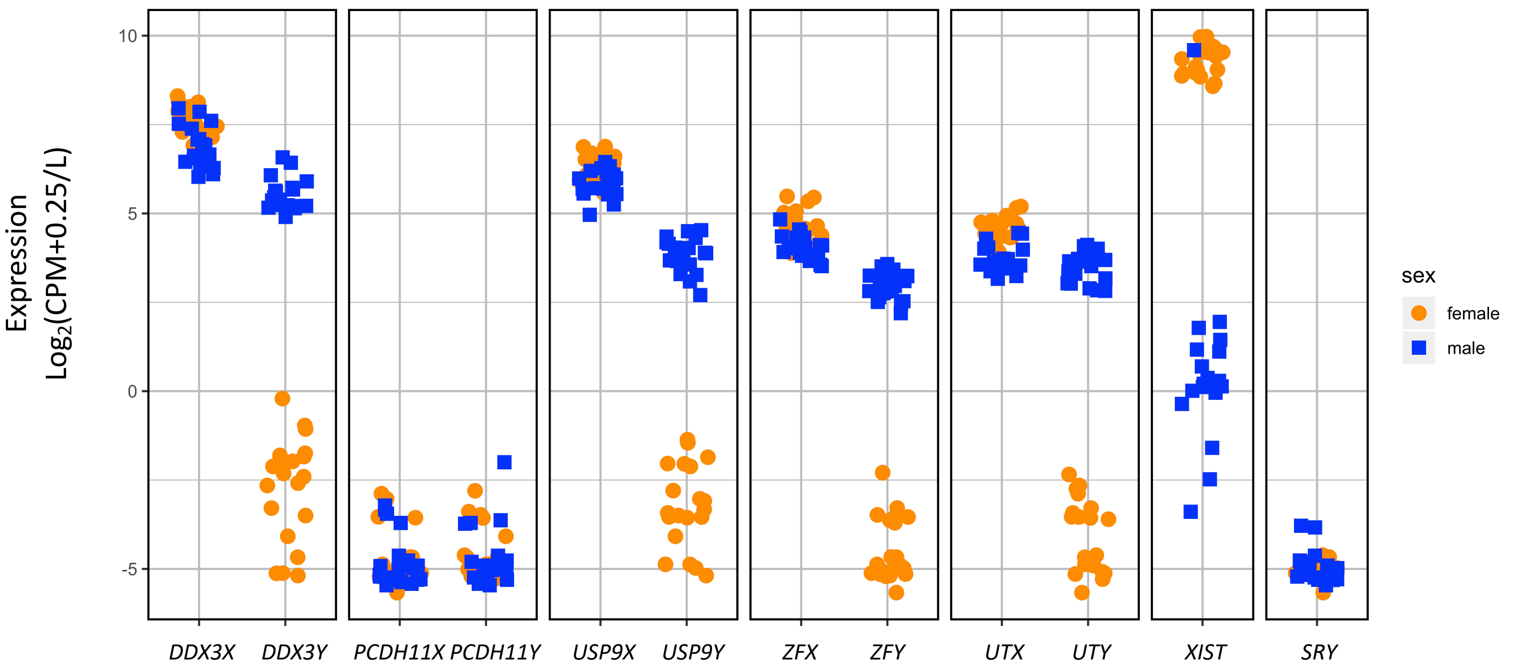


P LIVER aligned to STAR and sex chromosome complement reference genome

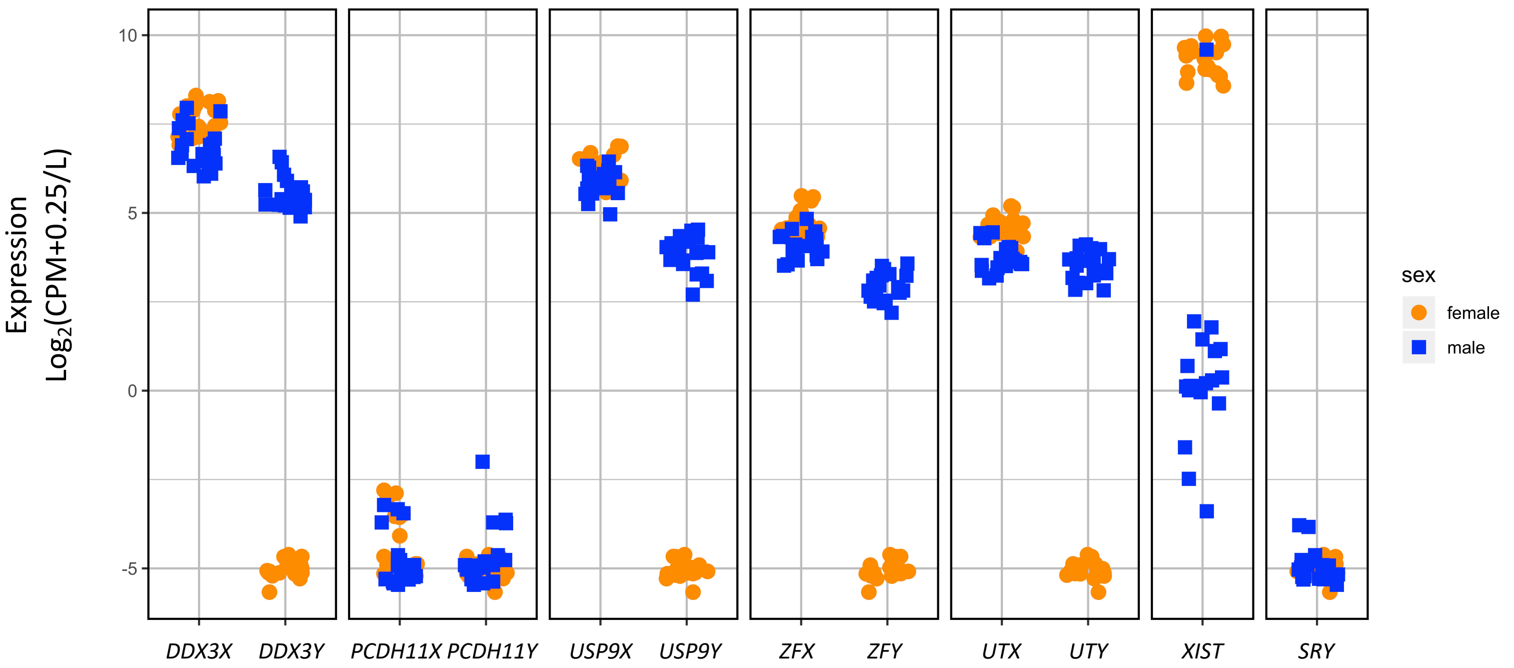

Q THYROID aligned to HISAT and default reference genome

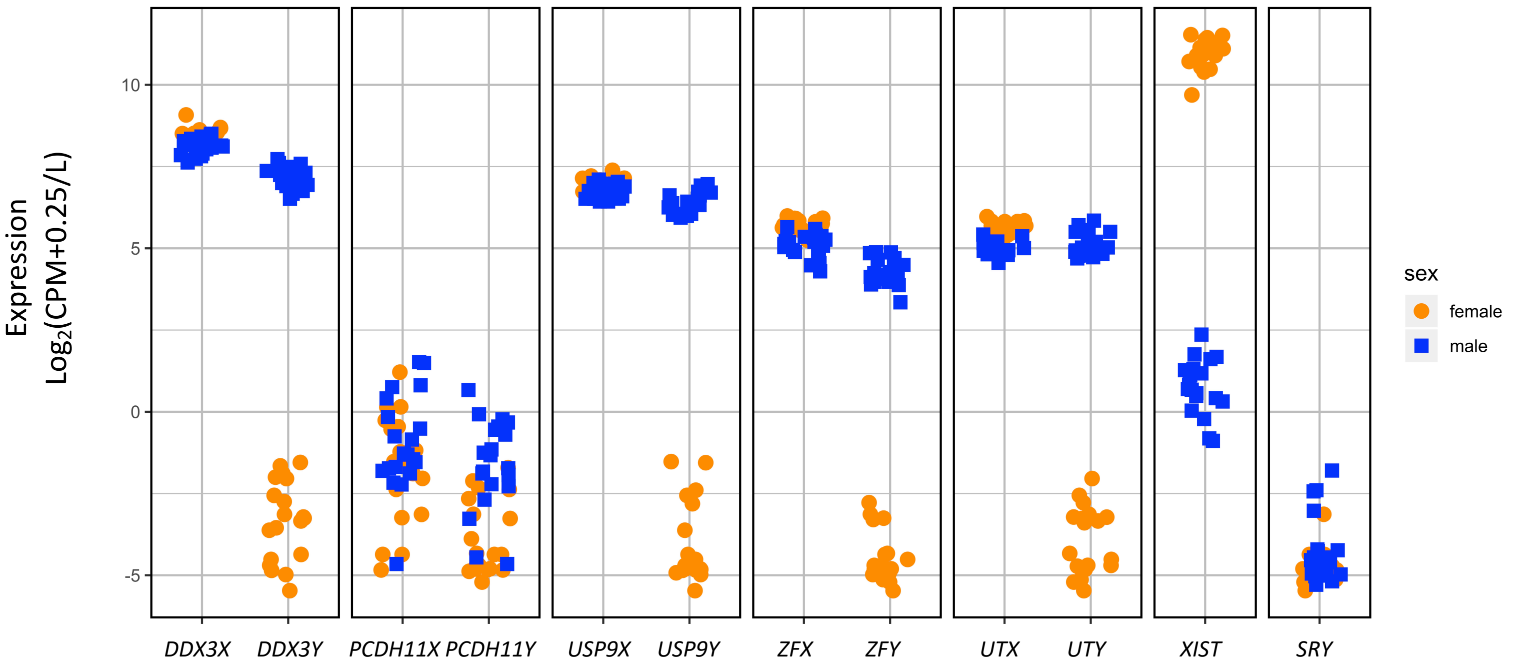


R THYROID aligned to HISAT and sex chromosome complement reference genome

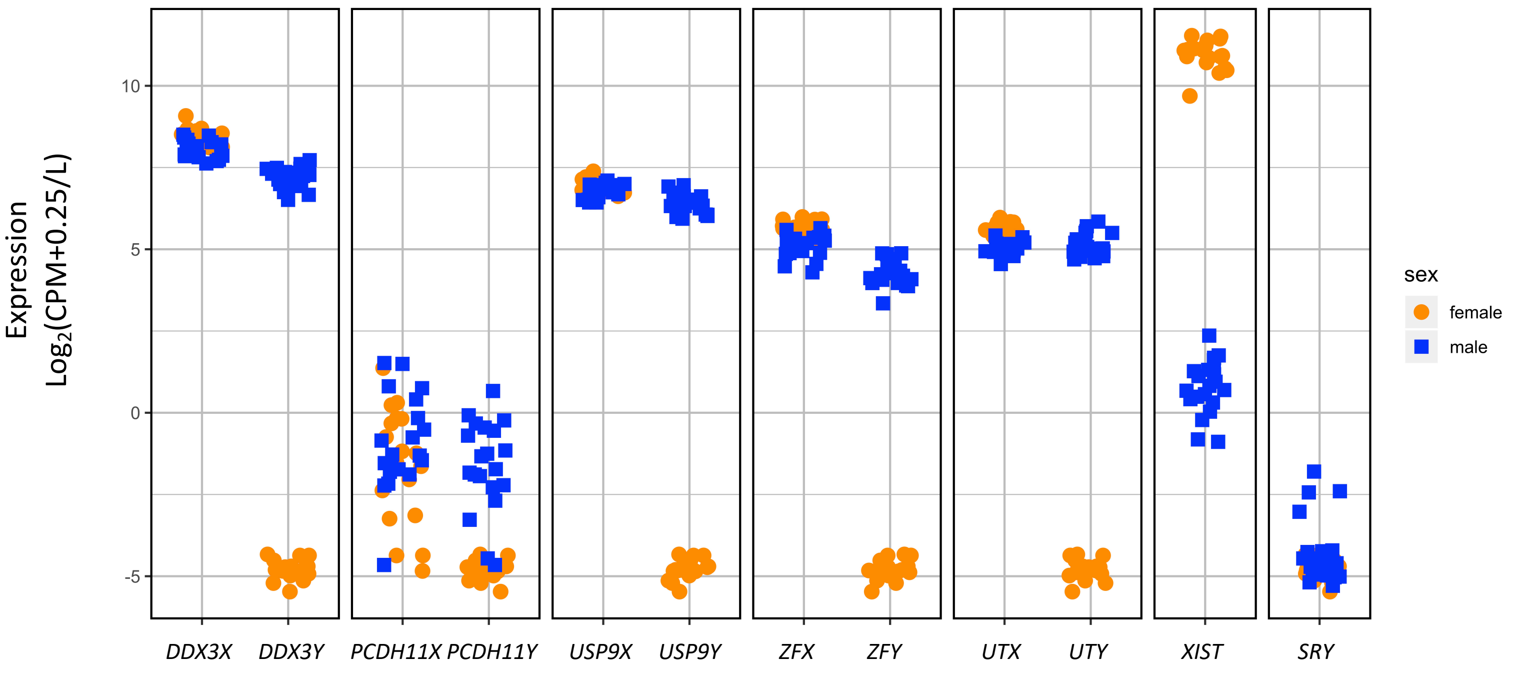

S THYROID aligned to STAR and default reference genome

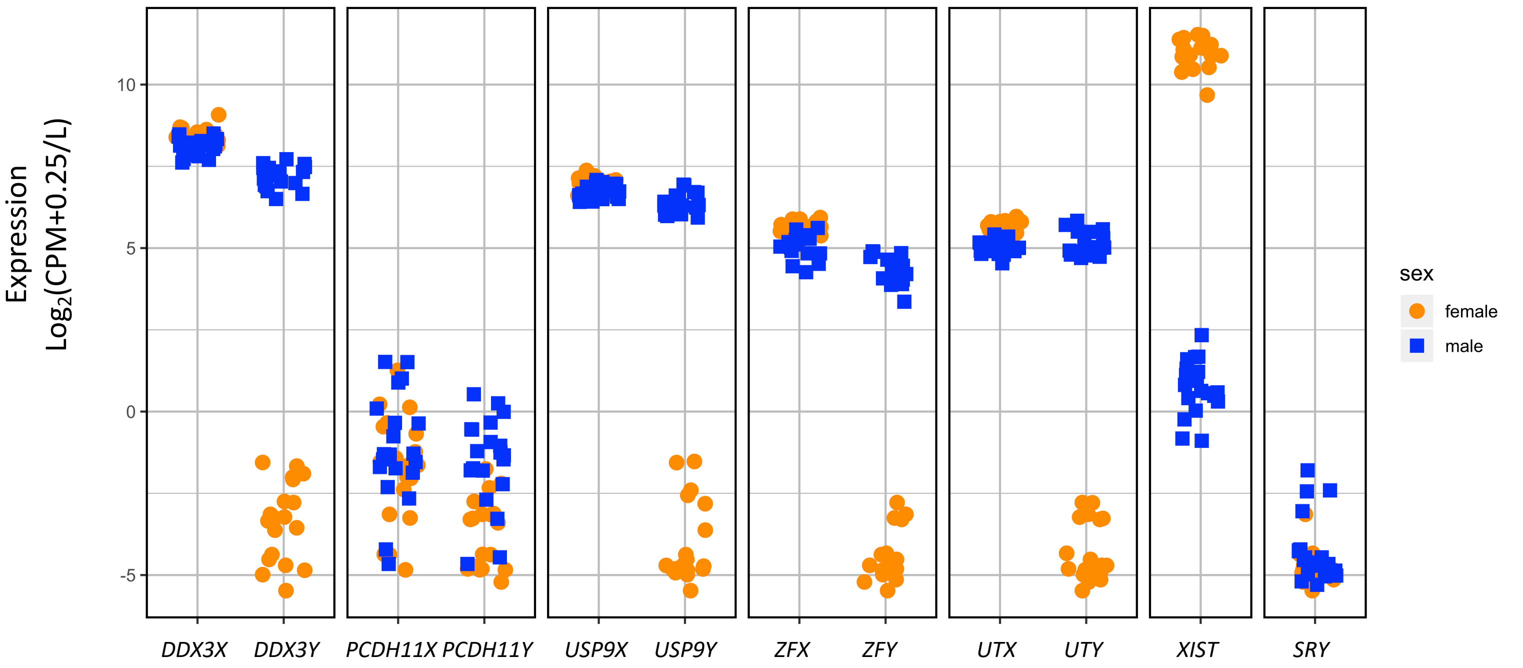


T THYROID aligned to STAR and sex chromosome complement reference genome

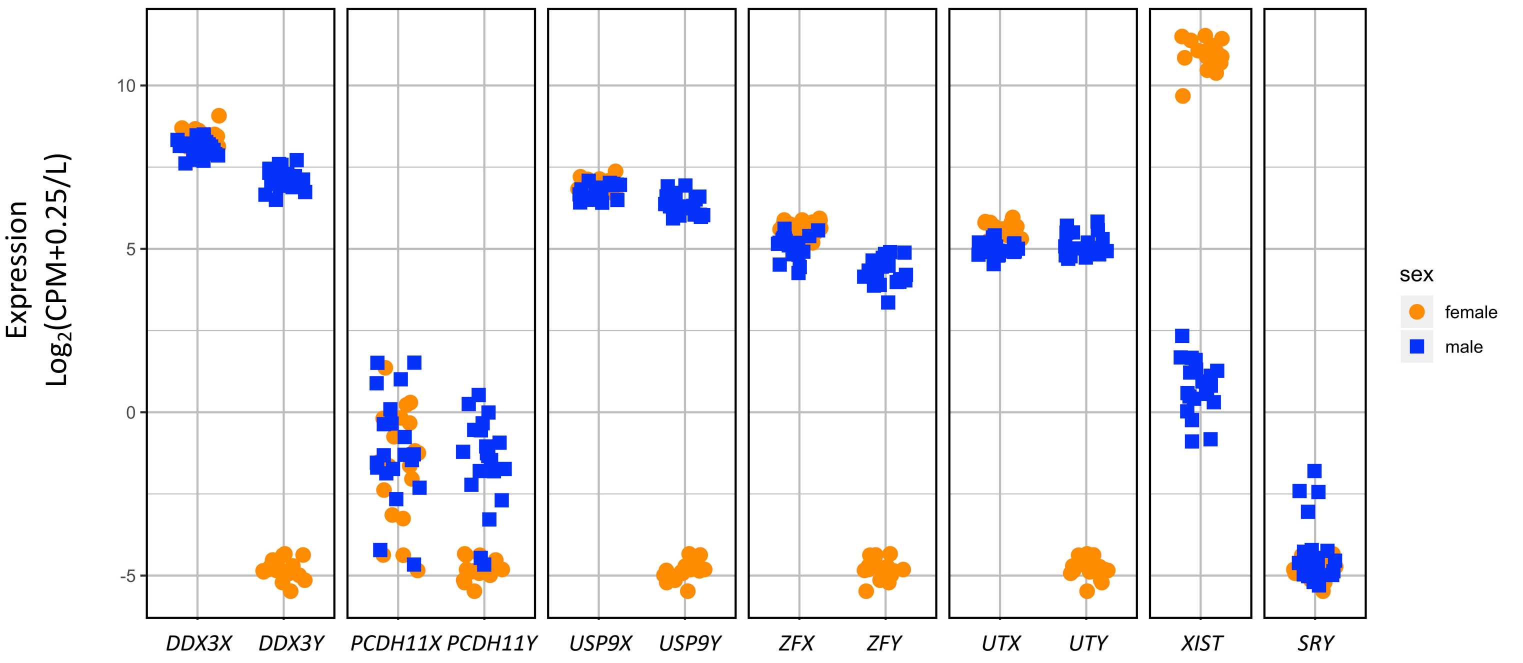
