## Additional file 6 for "Reference genome and transcriptome informed by the sex chromosome complement of the sample increases ability to detect sex differences in gene expression from RNA-Seq data"

A) Blood PC1 & PC2

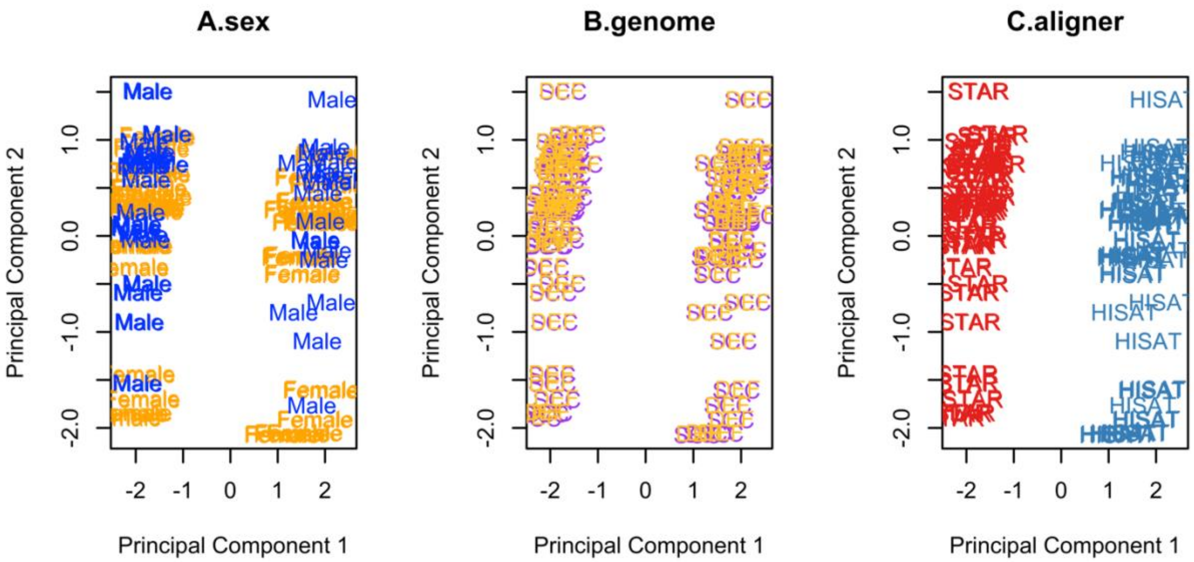


B) Blood PC2 & PC3

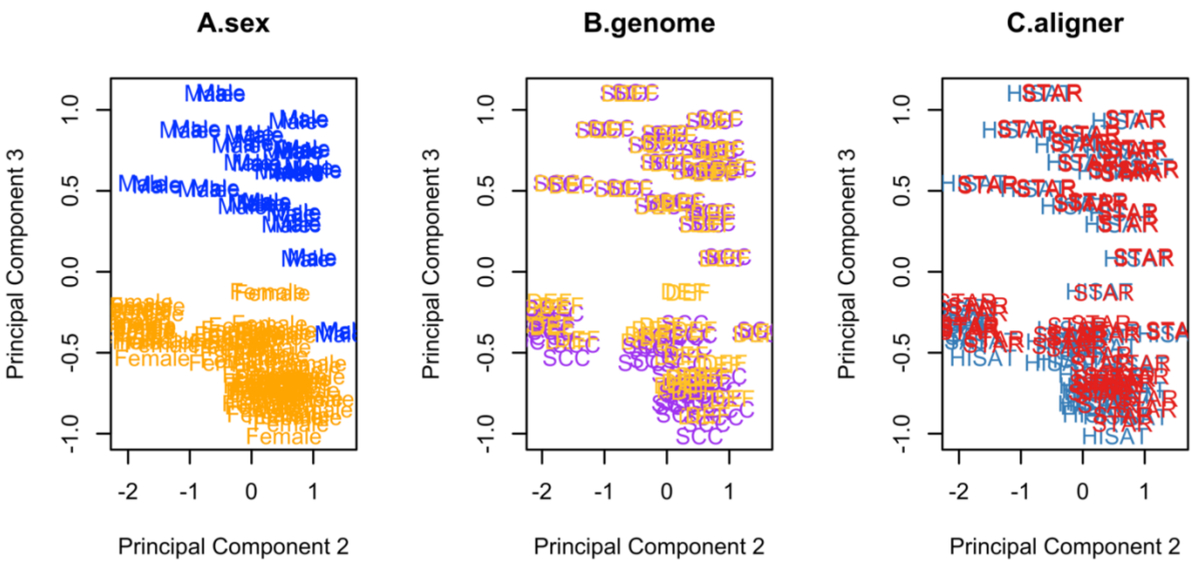

C) Brain cortex PC1 & PC2

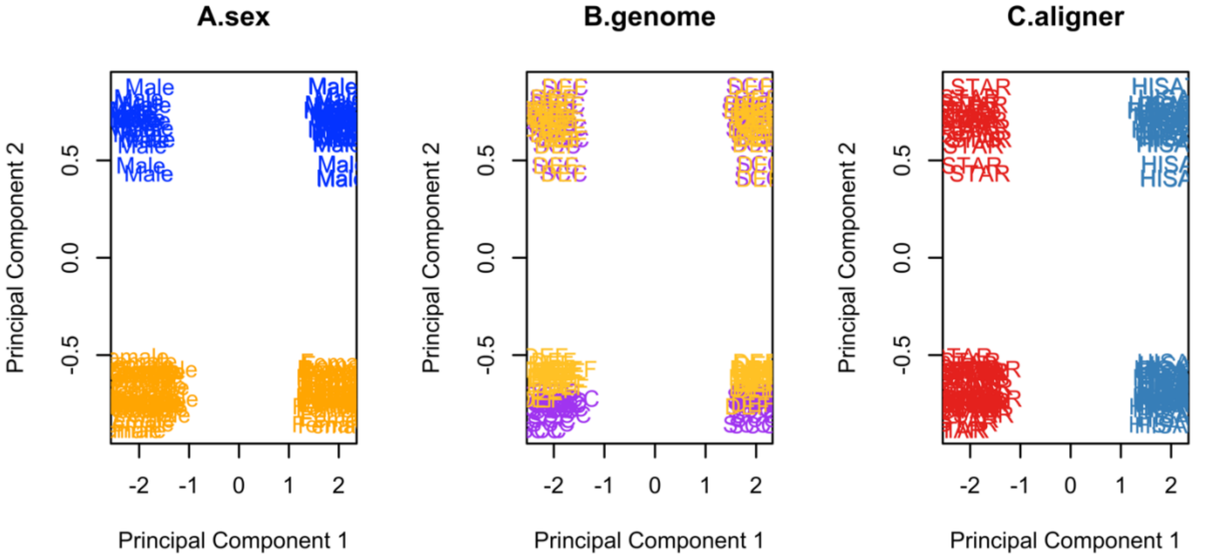


D) Brain cortex PC2 & PC3

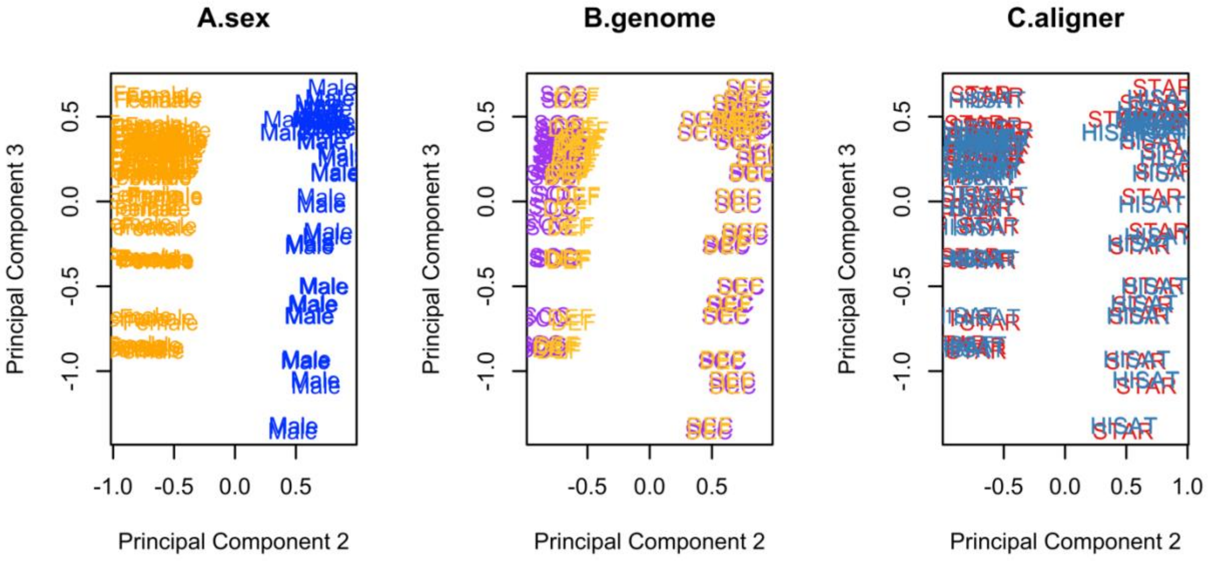

E) Breast PC1 & PC2

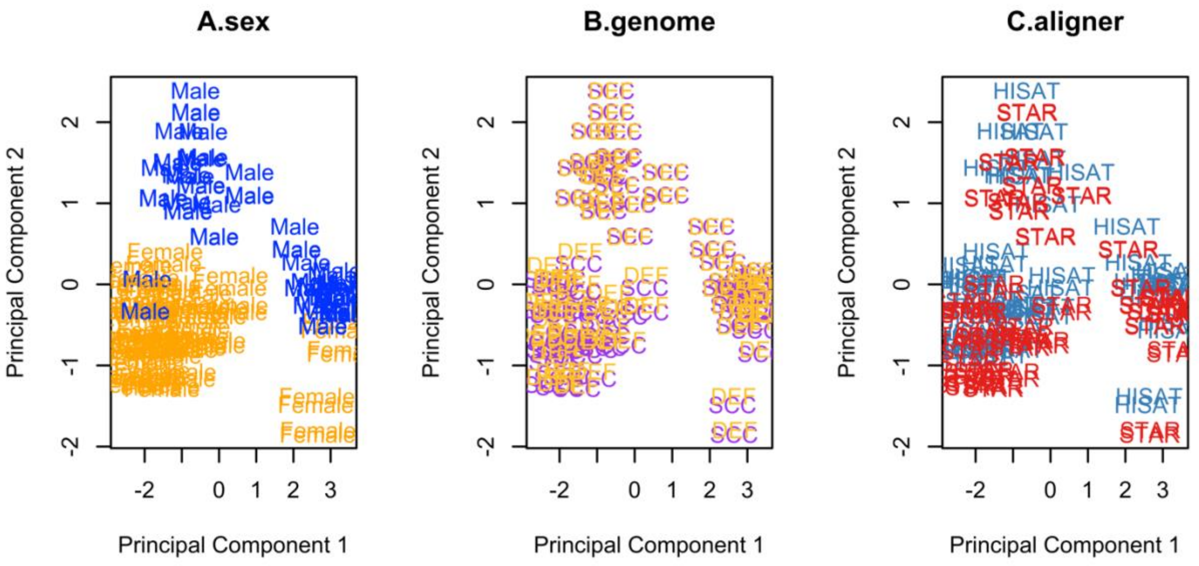


F) Breast PC2 & PC3

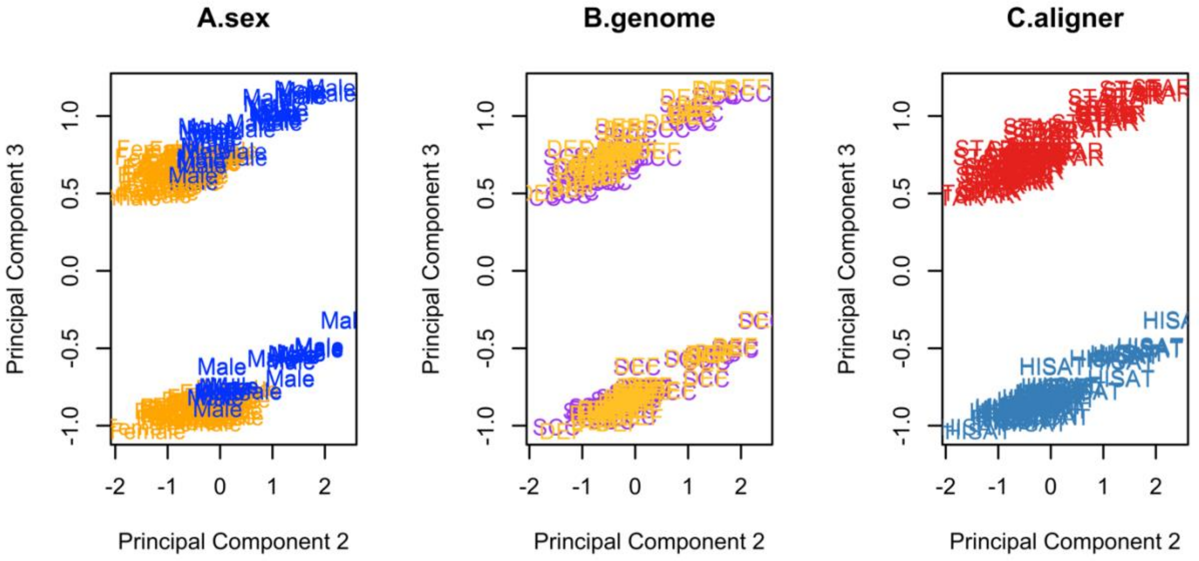

G) Liver PC1 & PC2

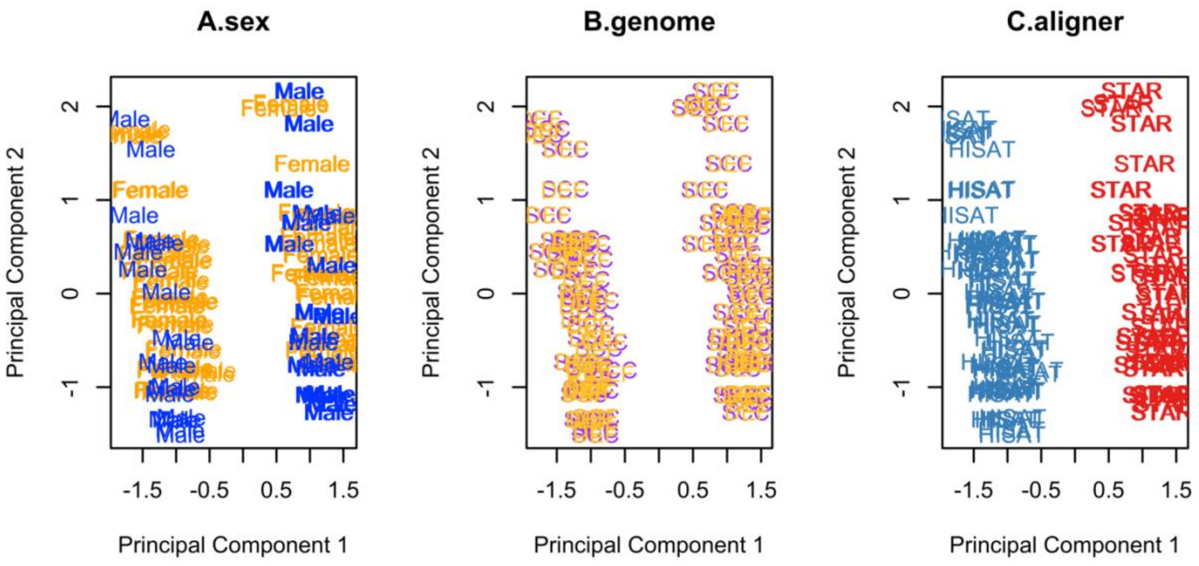


H) Liver PC2 & PC3

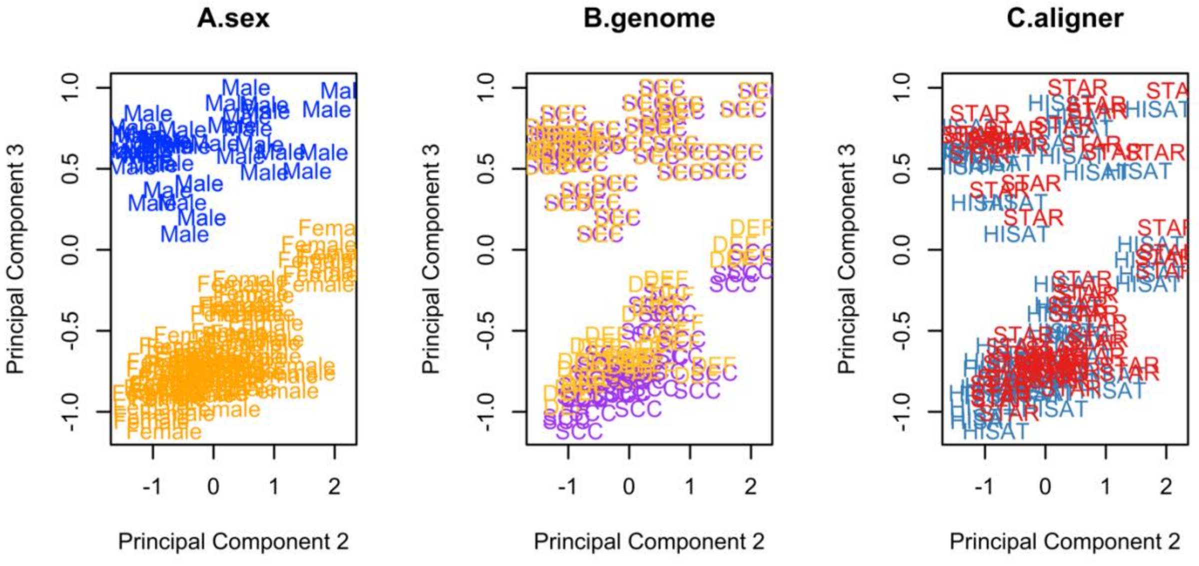

I) Thyroid PC1 & PC2

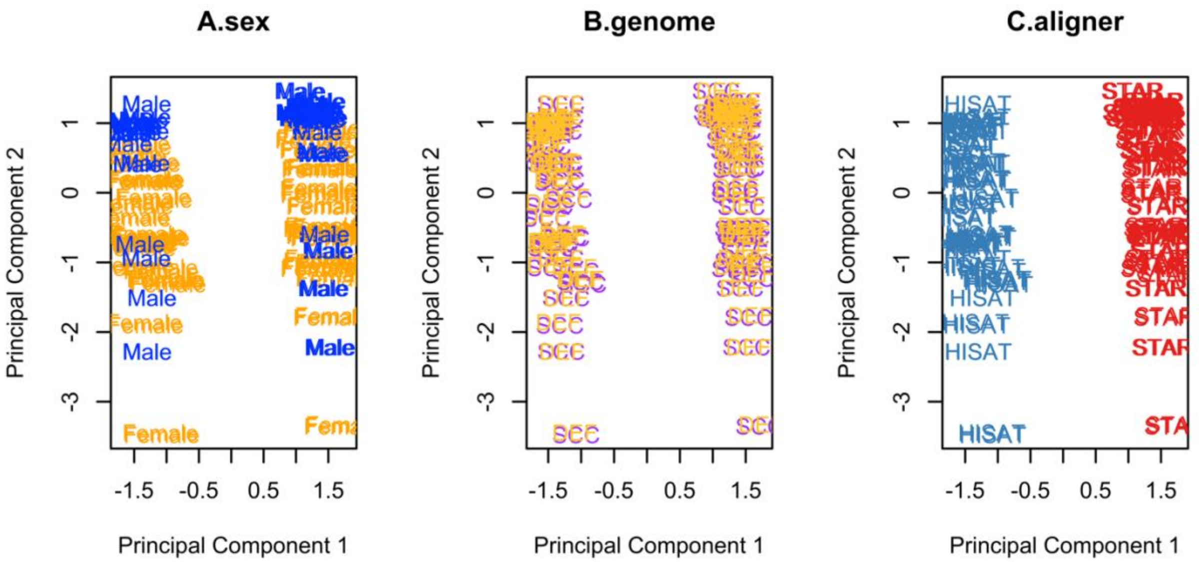


J) Thyroid PC2 & PC3

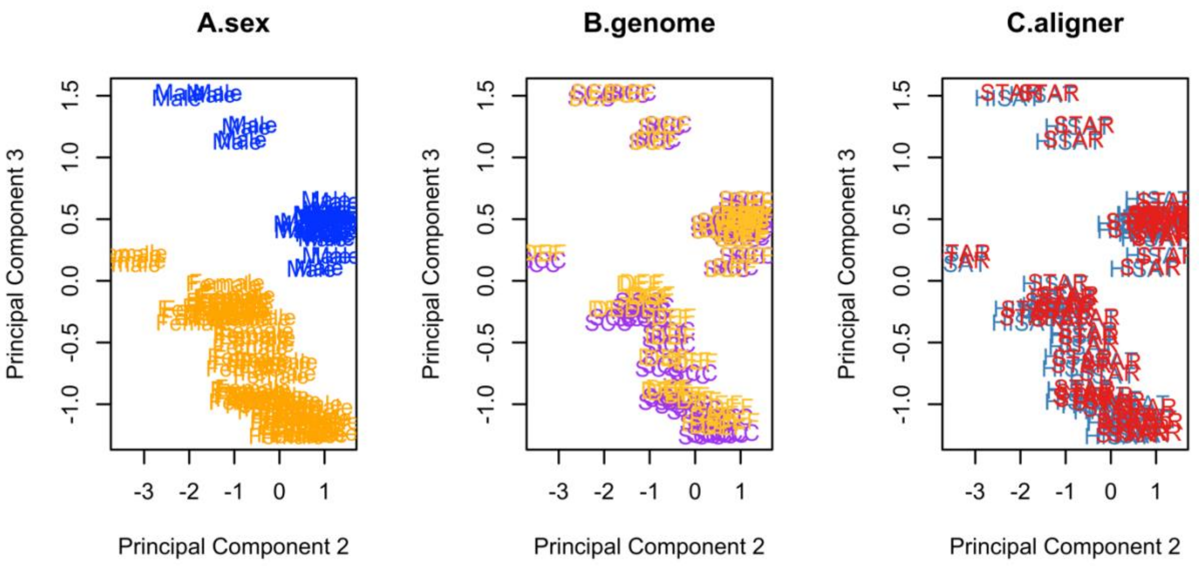
