## Additional file 10 for "Reference genome and transcriptome informed by the sex chromosome complement of the sample increases ability to detect sex differences in gene expression from RNA-Seq data"

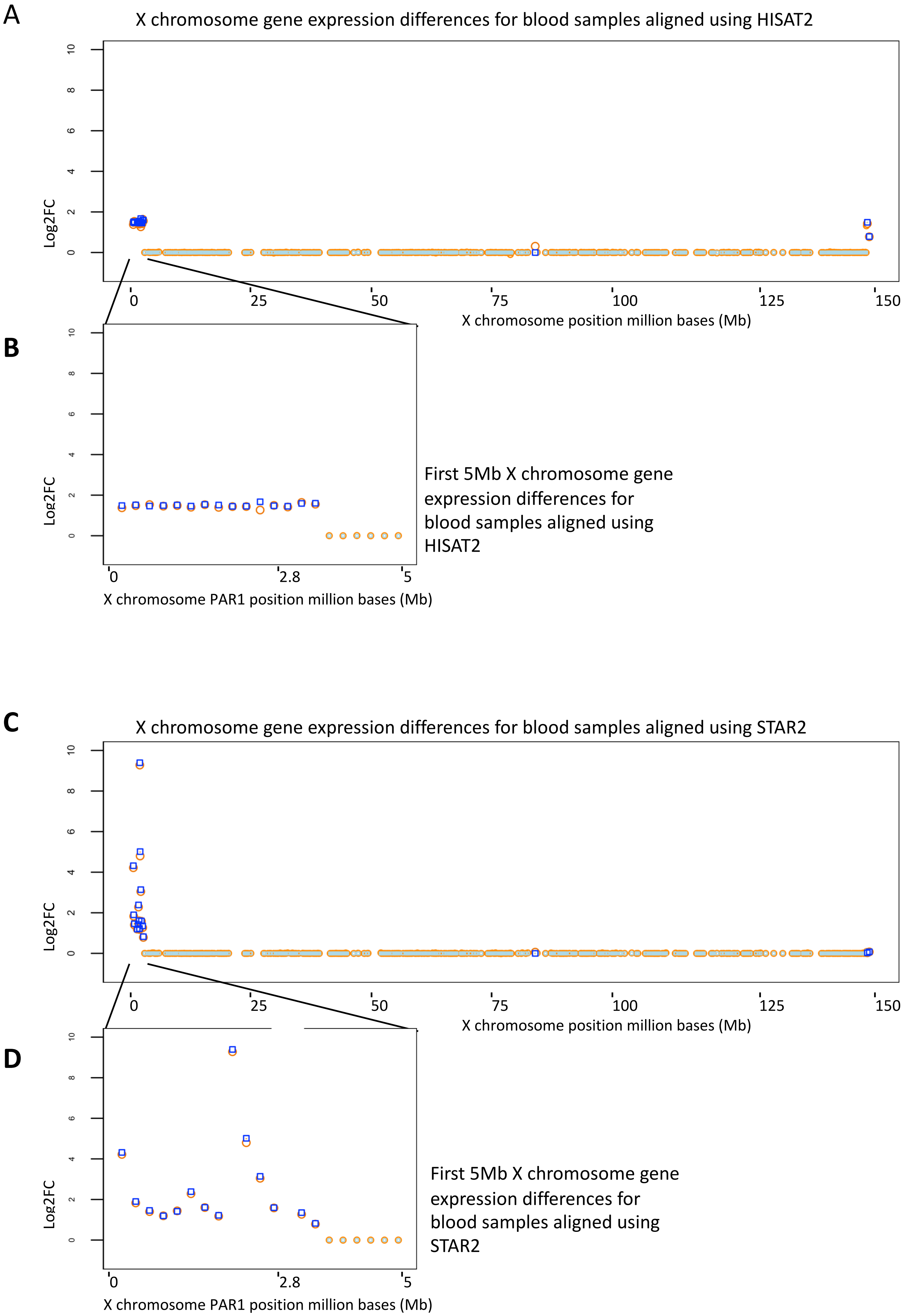
A
X chromosome gene expression differences for blood samples aligned using HISAT2
Log2FC
X chromosome position million bases (Mb)
B
First 5Mb X chromosome gene expression differences for blood samples aligned using HISAT2
X chromosome PAR1 position million bases (Mb)
C
X chromosome gene expression differences for blood samples aligned using STAR2
X chromosome position million bases (Mb)
D
First 5Mb X chromosome gene expression differences for blood samples aligned using STAR2
X chromosome PAR1 position million bases (Mb)

E

X chromosome gene expression differences for brain samples aligned using HISAT2

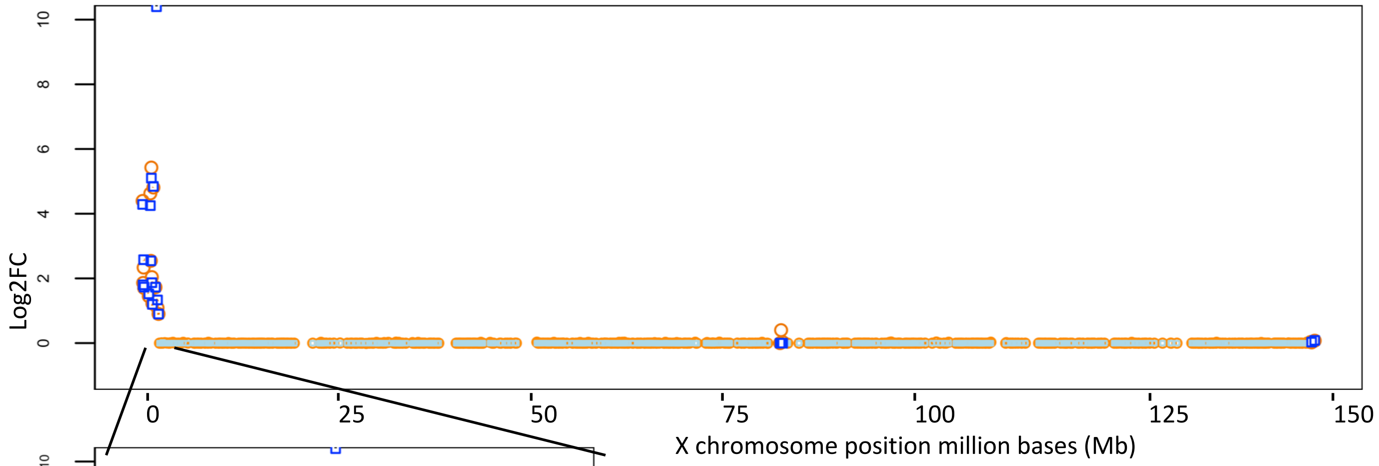


F

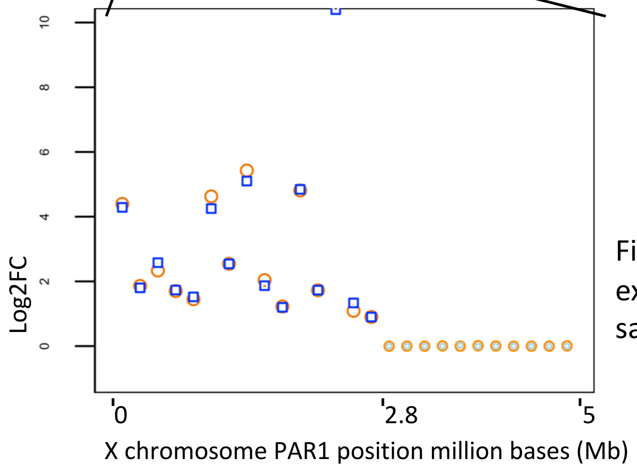


First 5Mb X chromosome gene expression differences for brain samples aligned using HISAT

G

X chromosome gene expression differences for brain samples aligned using STAR2

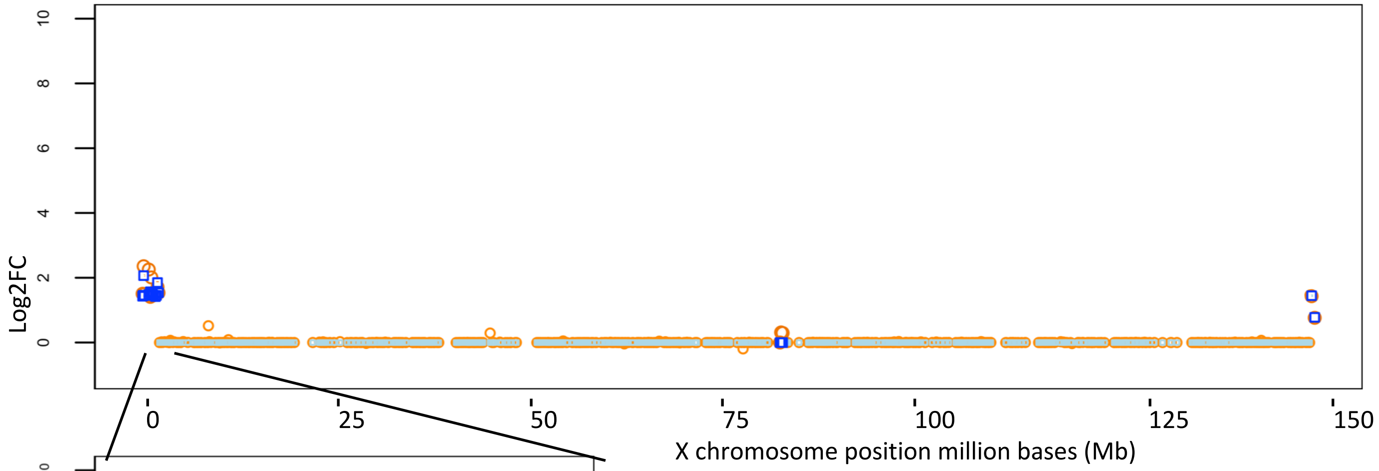


H

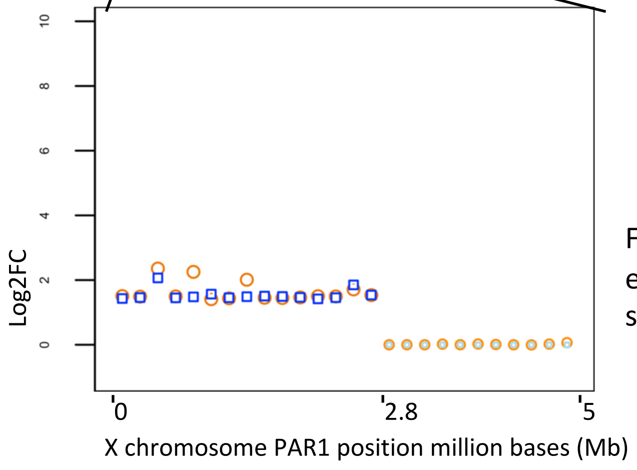


First 5Mb X chromosome gene expression differences for brain samples aligned using STAR2

I

X chromosome gene expression differences for breast samples aligned using HISAT2

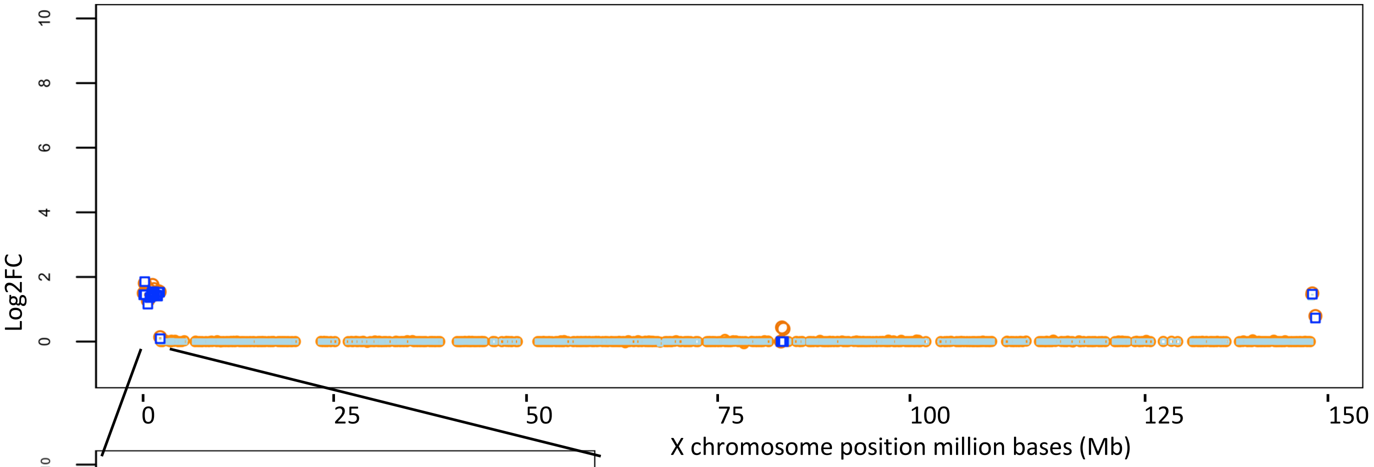


J

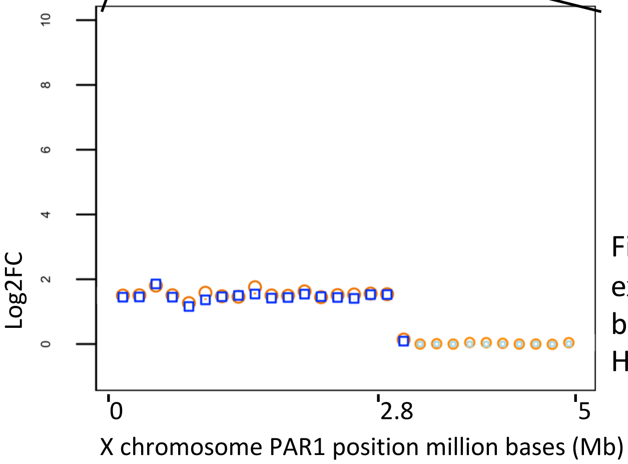


First 5Mb X chromosome gene expression differences for breast samples aligned using HISAT2

K

X chromosome gene expression differences for breast samples aligned using STAR2

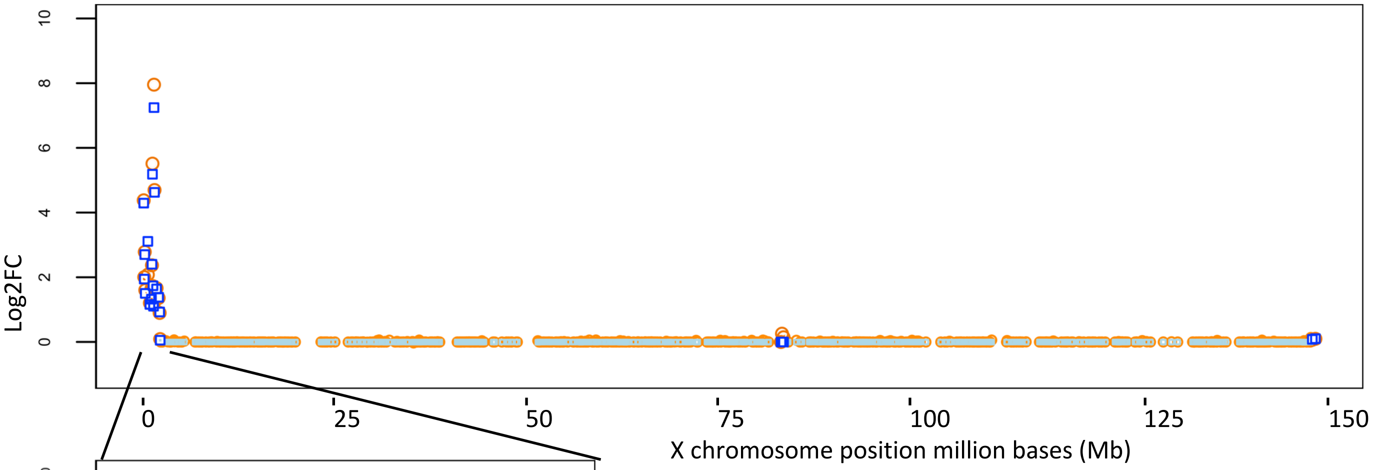


L

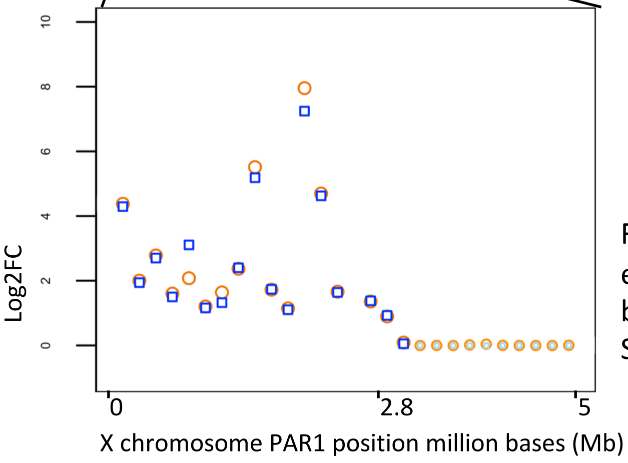


First 5Mb X chromosome gene expression differences for breast samples aligned using STAR2

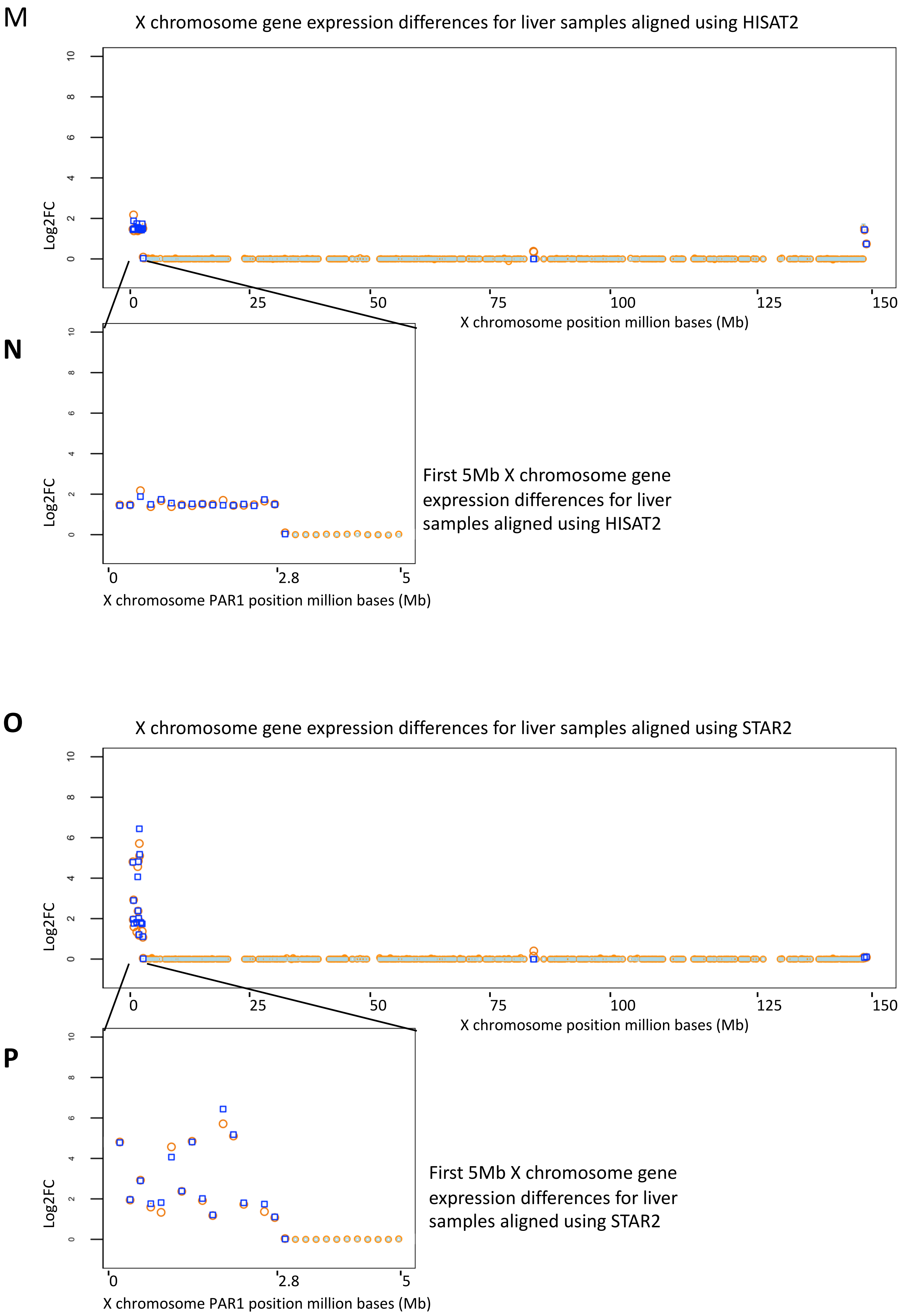

M
X chromosome gene expression differences for liver samples aligned using HISAT2
Log2FC
0
2
4
6
8
10
0
25
50
75
100
125
150
X chromosome position million bases (Mb)
N
Log2FC
0
2
4
6
8
10
0
2.8
5
X chromosome PAR1 position million bases (Mb)
First 5Mb X chromosome gene expression differences for liver samples aligned using HISAT2
O
X chromosome gene expression differences for liver samples aligned using STAR2
Log2FC
0
2
4
6
8
10
0
25
50
75
100
125
150
X chromosome position million bases (Mb)
P
Log2FC
0
2
4
6
8
10
0
2.8
5
X chromosome PAR1 position million bases (Mb)
First 5Mb X chromosome gene expression differences for liver samples aligned using STAR2

Q

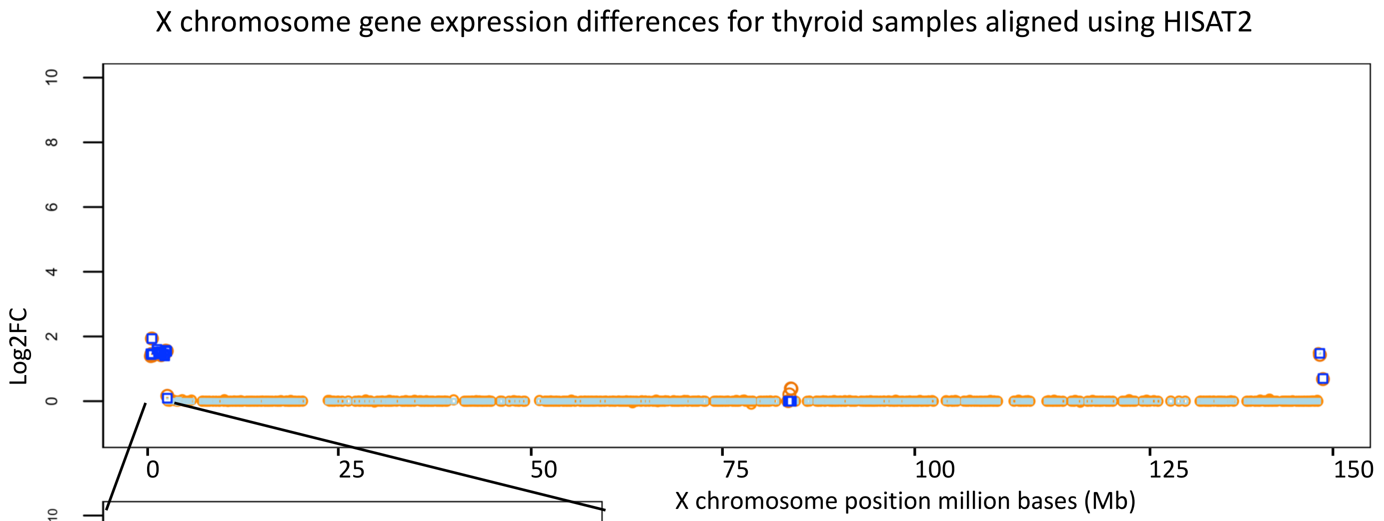
X chromosome gene expression differences for thyroid samples aligned using HISAT2
Log2FC
X chromosome position million bases (Mb)
0
25
50
75
100
125
150

R

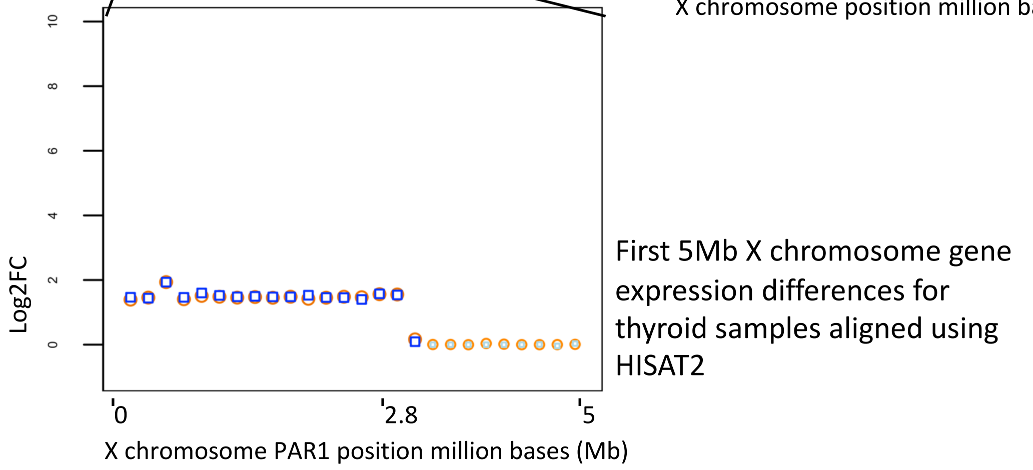
Log2FC
0
2.8
5
X chromosome PAR1 position million bases (Mb)
First 5Mb X chromosome gene expression differences for thyroid samples aligned using HISAT2

S

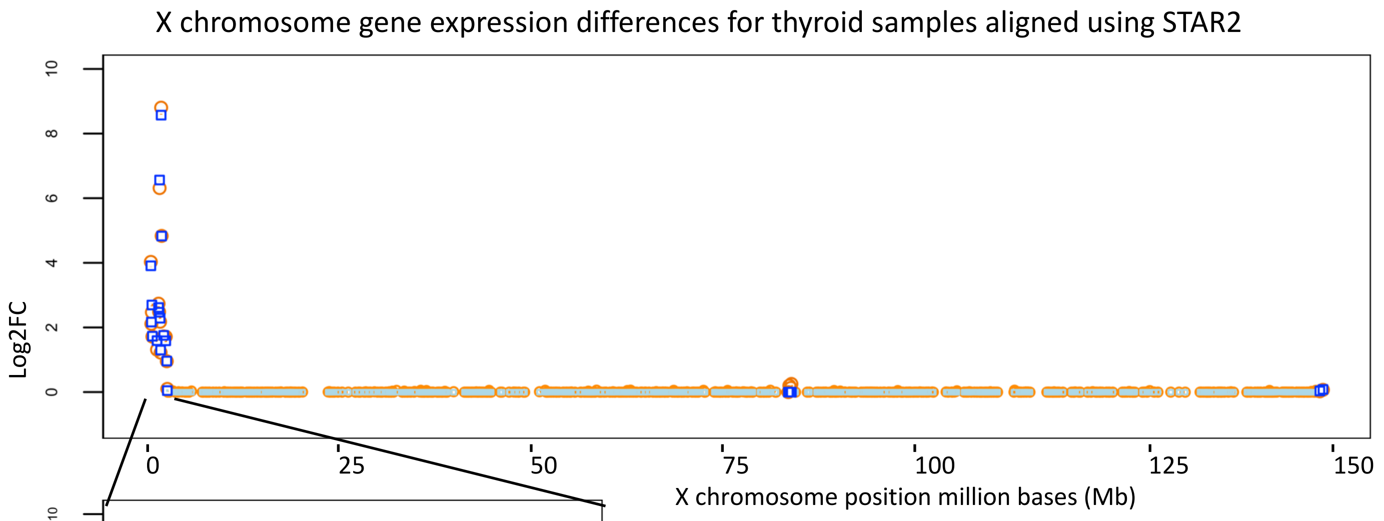
X chromosome gene expression differences for thyroid samples aligned using STAR2
Log2FC
X chromosome position million bases (Mb)
0
25
50
75
100
125
150

T

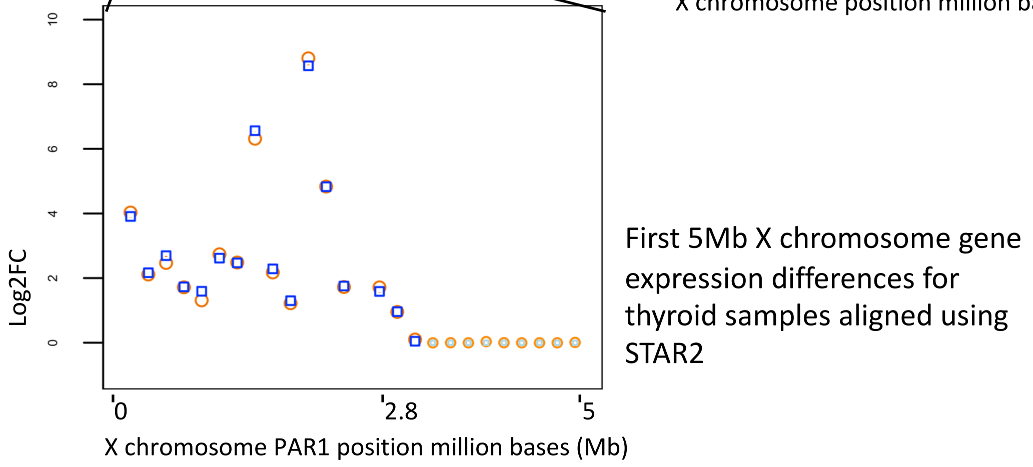
Log2FC
0
2.8
5
X chromosome PAR1 position million bases (Mb)
First 5Mb X chromosome gene expression differences for thyroid samples aligned using STAR2
