## Additional file 15 for "Reference genome and transcriptome informed by the sex chromosome complement of the sample increases ability to detect sex differences in gene expression from RNA-Seq data"

A Gene expression differences between male XY and female XX blood samples aligned using HISAT

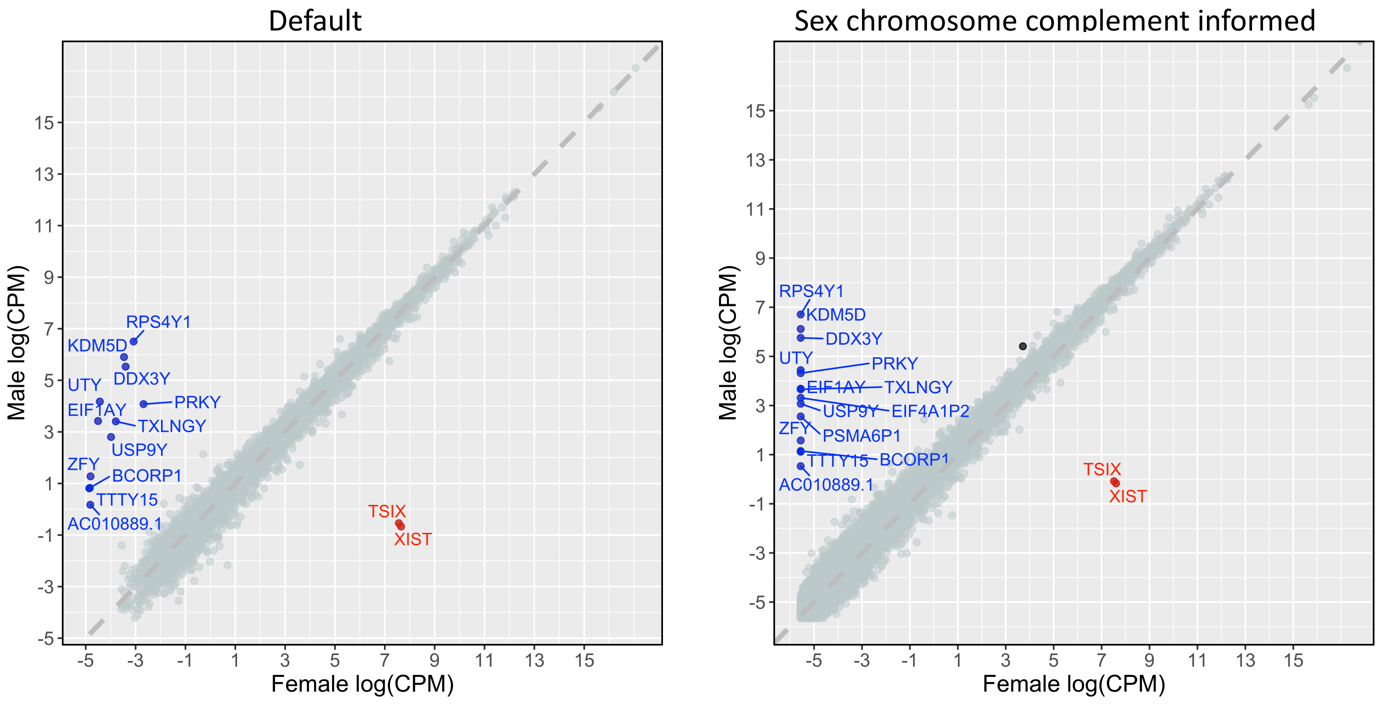


B Gene expression differences between male XY and female XX blood samples aligned using STAR

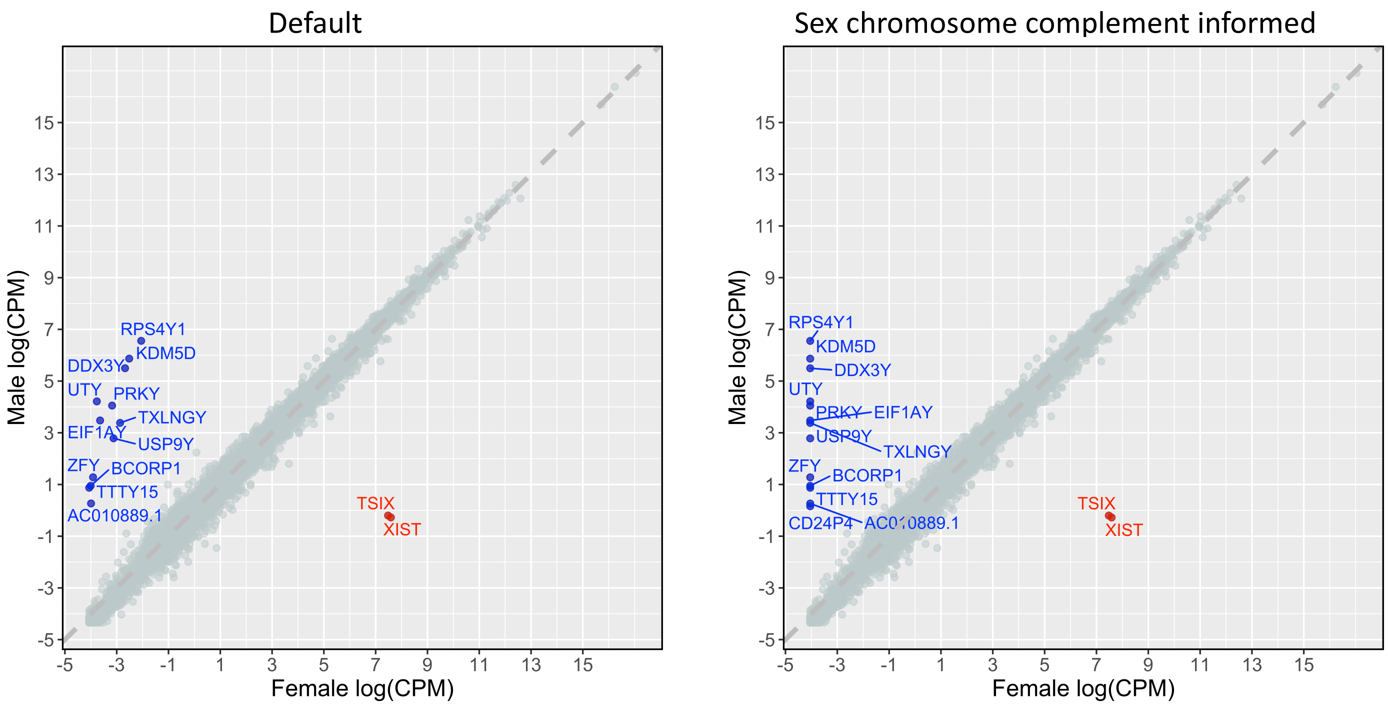

C Gene expression differences between male XY and female XX brain samples aligned using HISAT

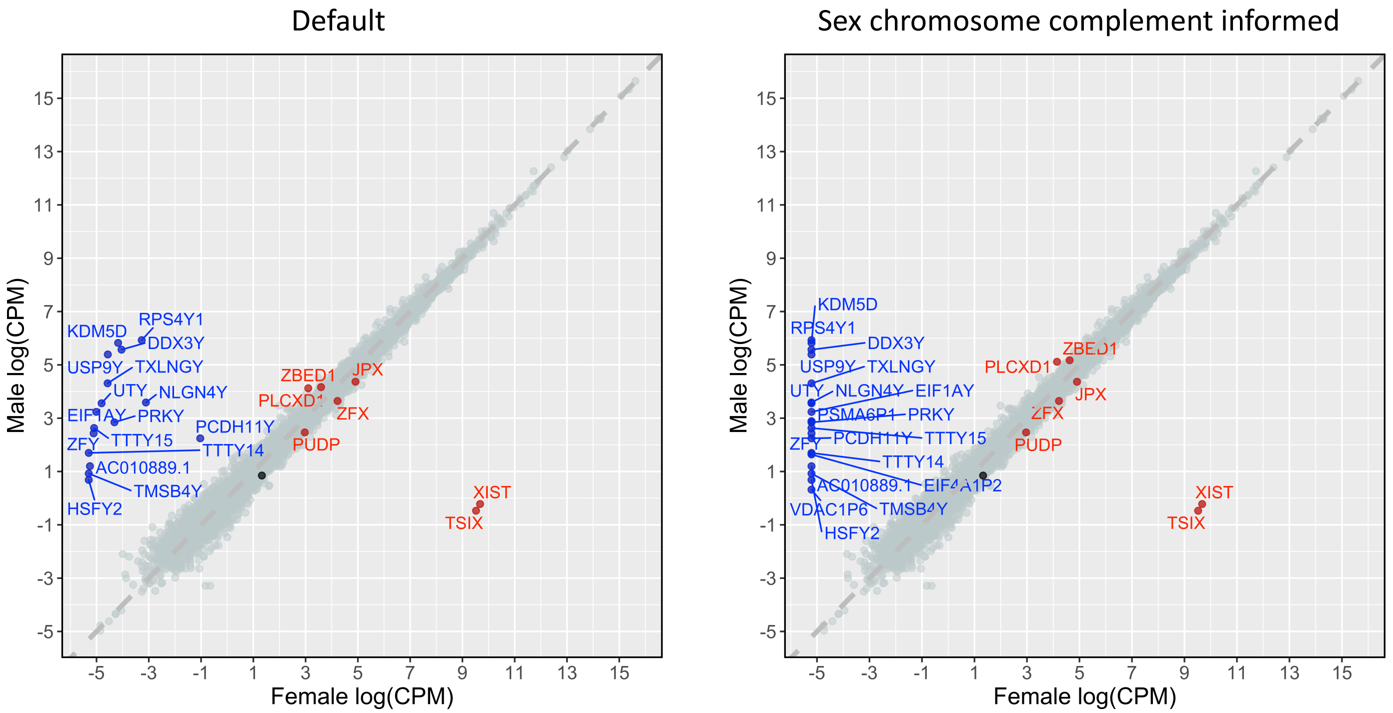


D Gene expression differences between male XY and female XX brain samples aligned using STAR

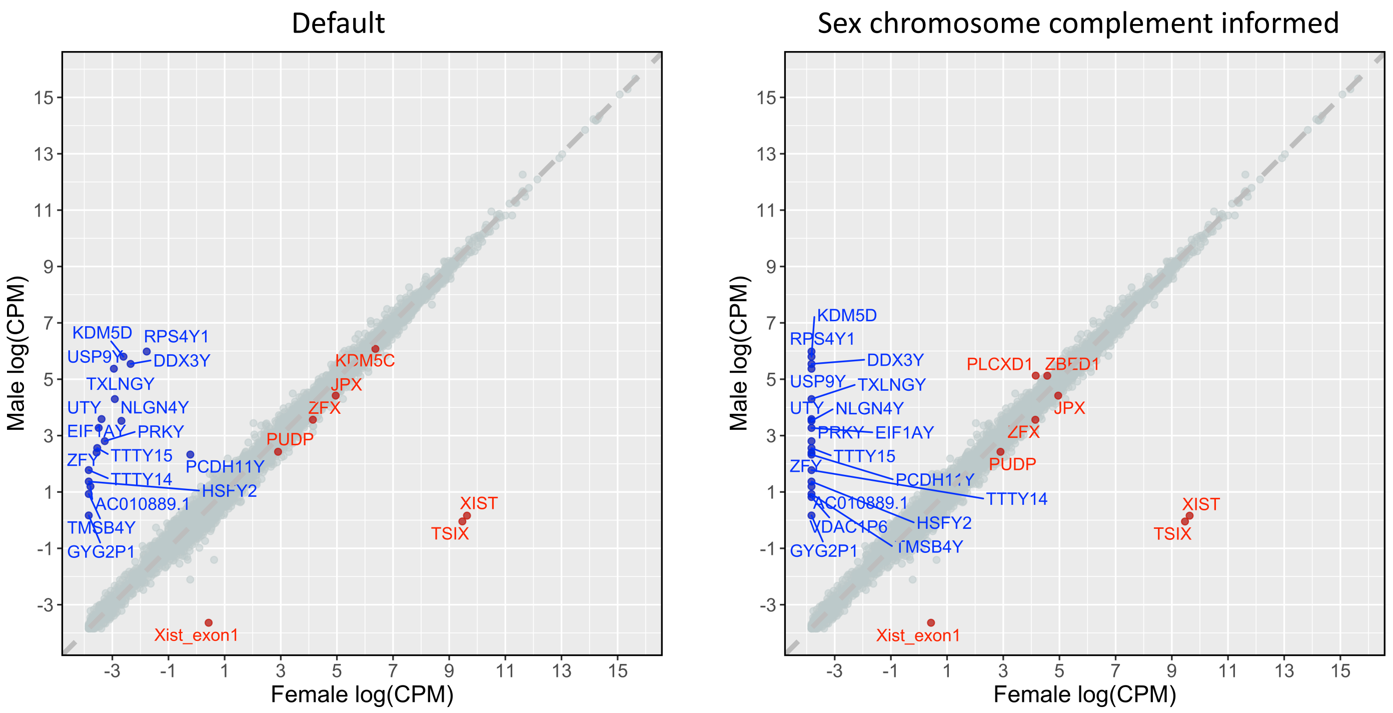

E Gene expression differences between male XY and female XX breast samples aligned using HISAT

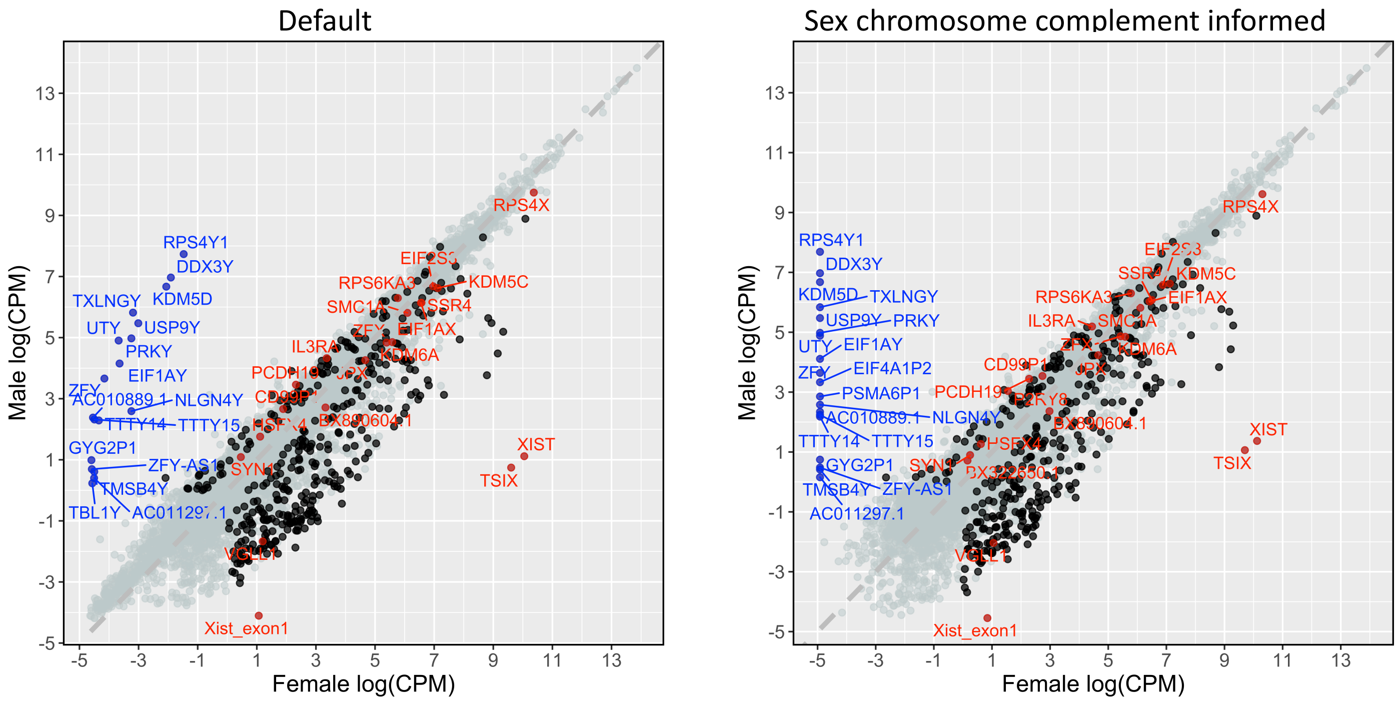


F Gene expression differences between male XY and female XX breast samples aligned using STAR

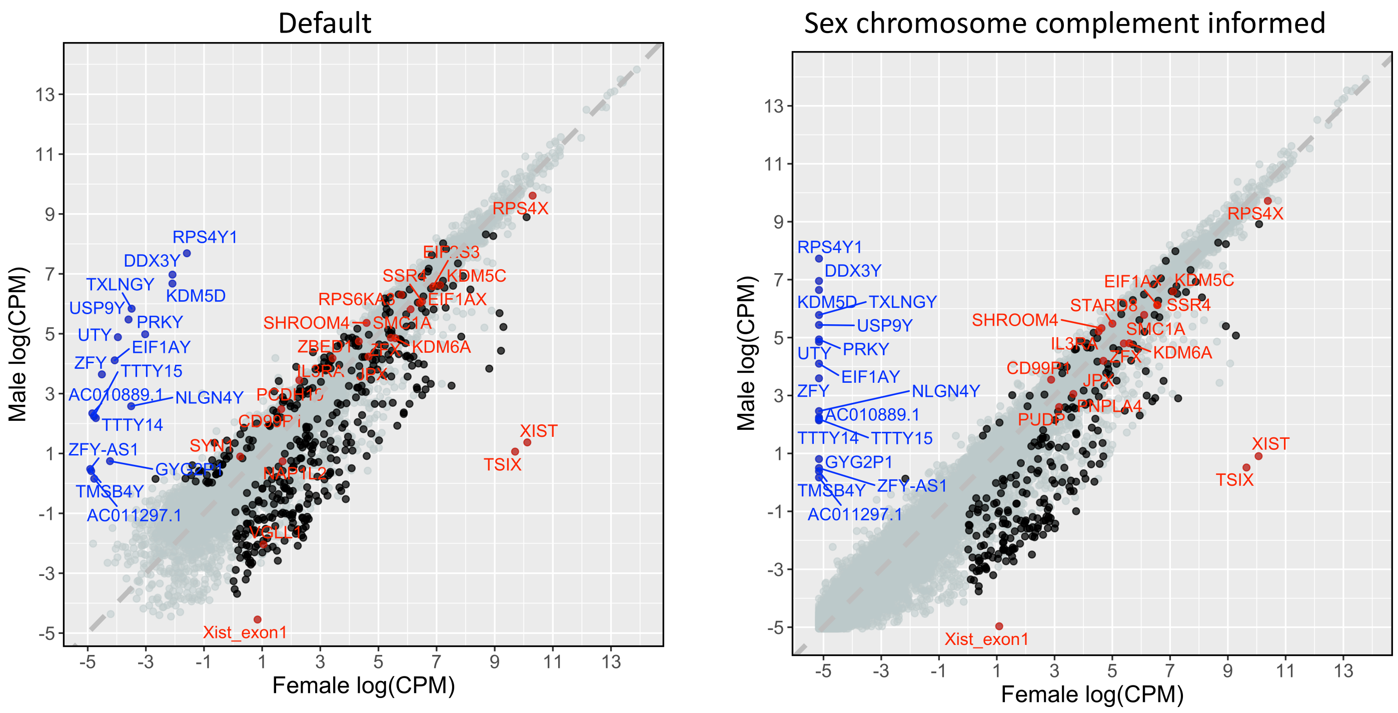

G Gene expression differences between male XY and female XX liver samples aligned using HISAT

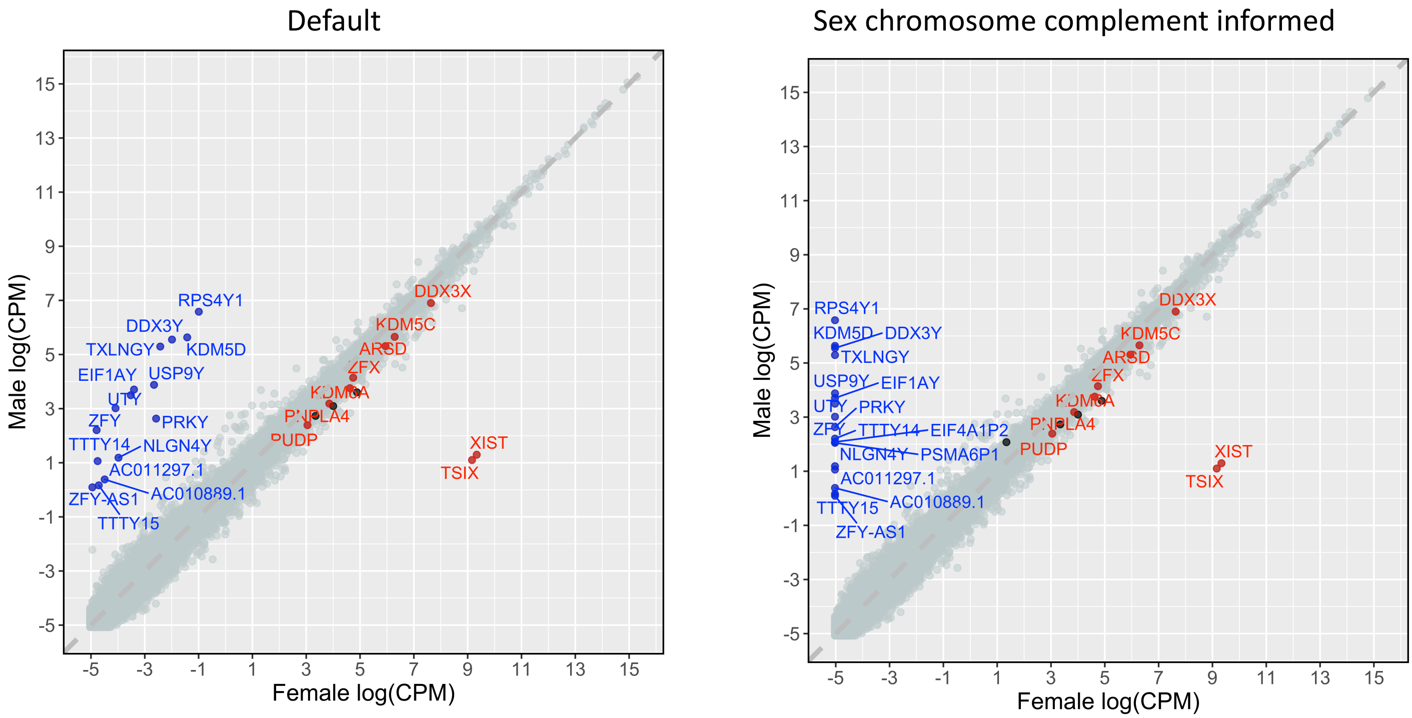


H Gene expression differences between male XY and female XX liver samples aligned using STAR

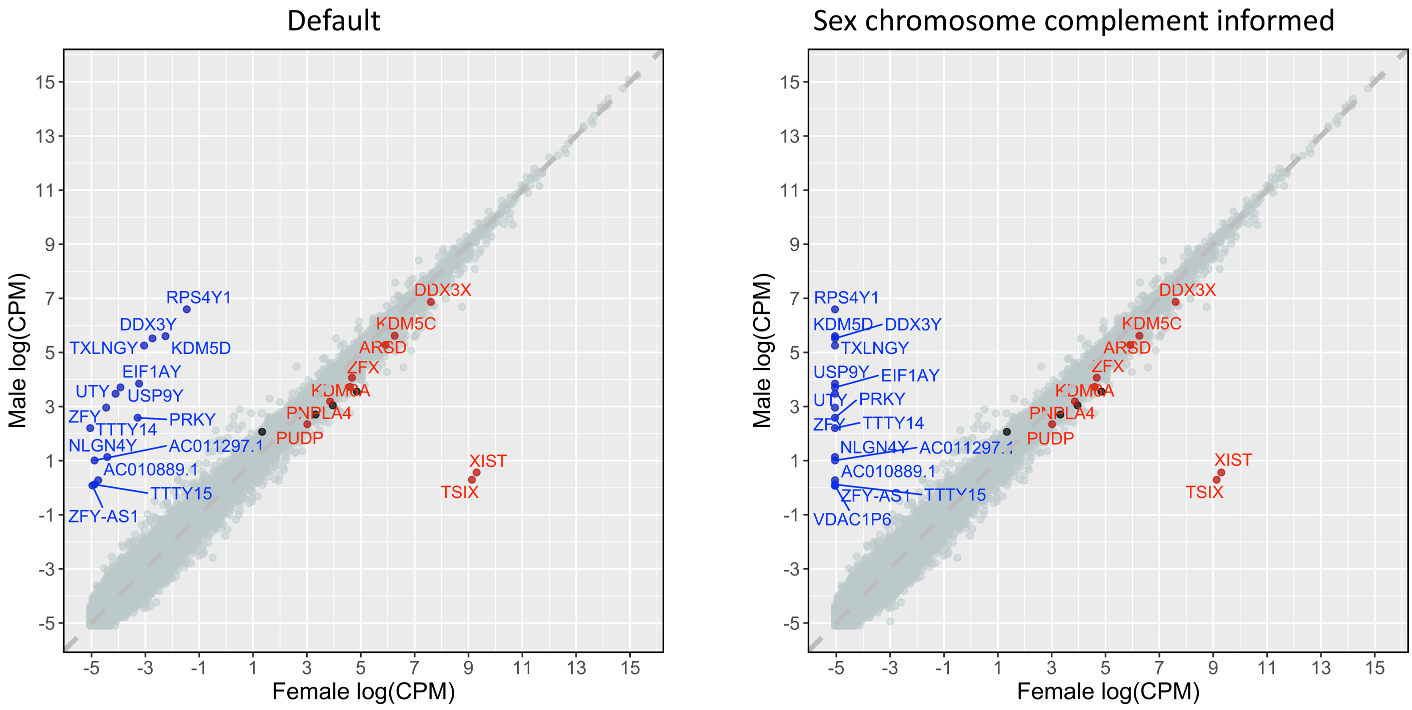

I Gene expression differences between male XY and female XX thyroid samples aligned using HISAT

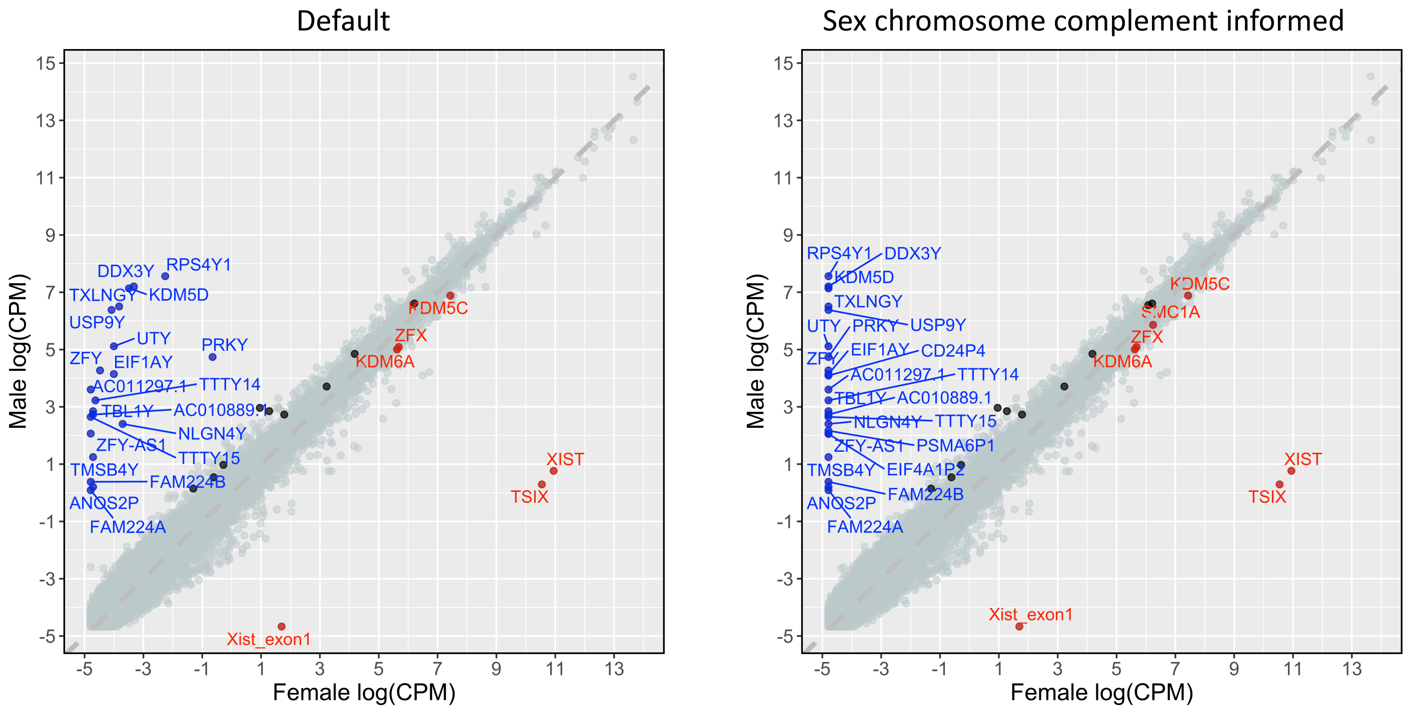


J Gene expression differences between male XY and female XX thyroid samples aligned using STAR

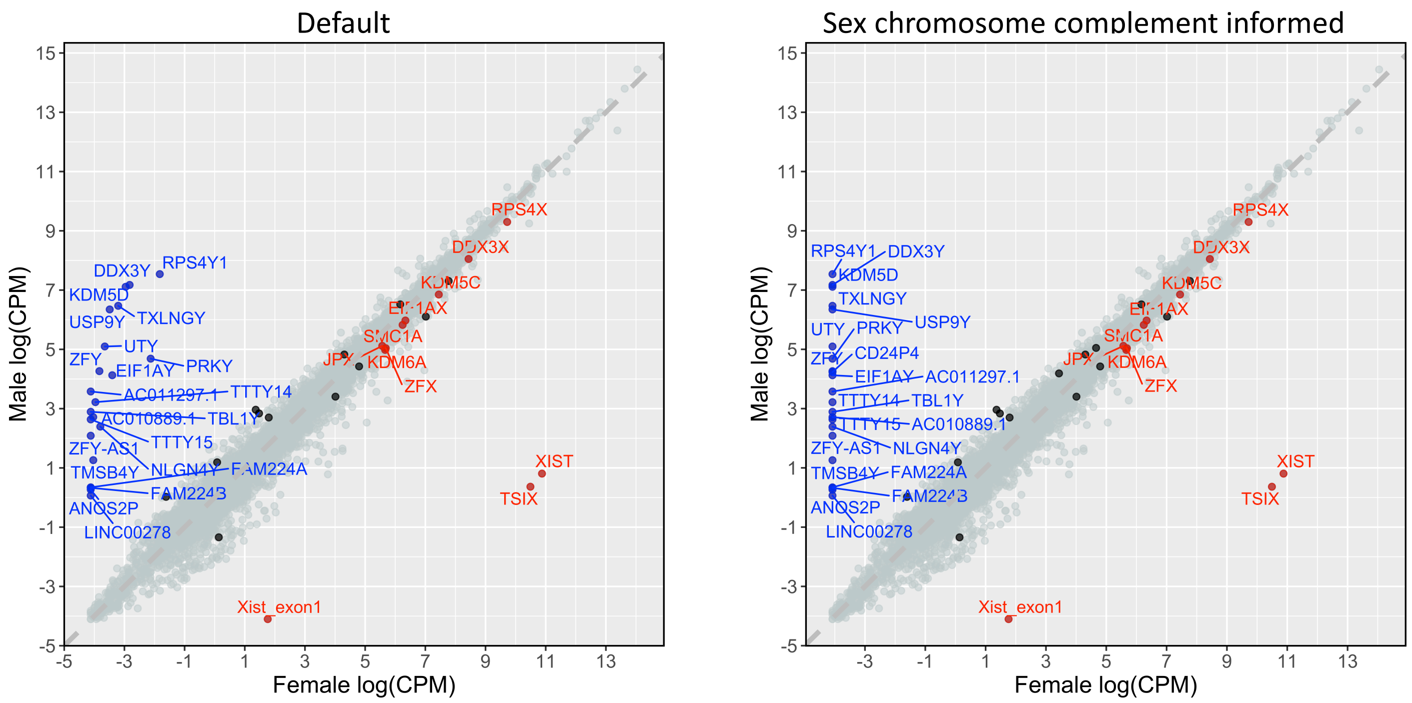
