## Additional file 19 for "Reference genome and transcriptome informed by the sex chromosome complement of the sample increases ability to detect sex differences in gene expression from RNA-Seq data"

A

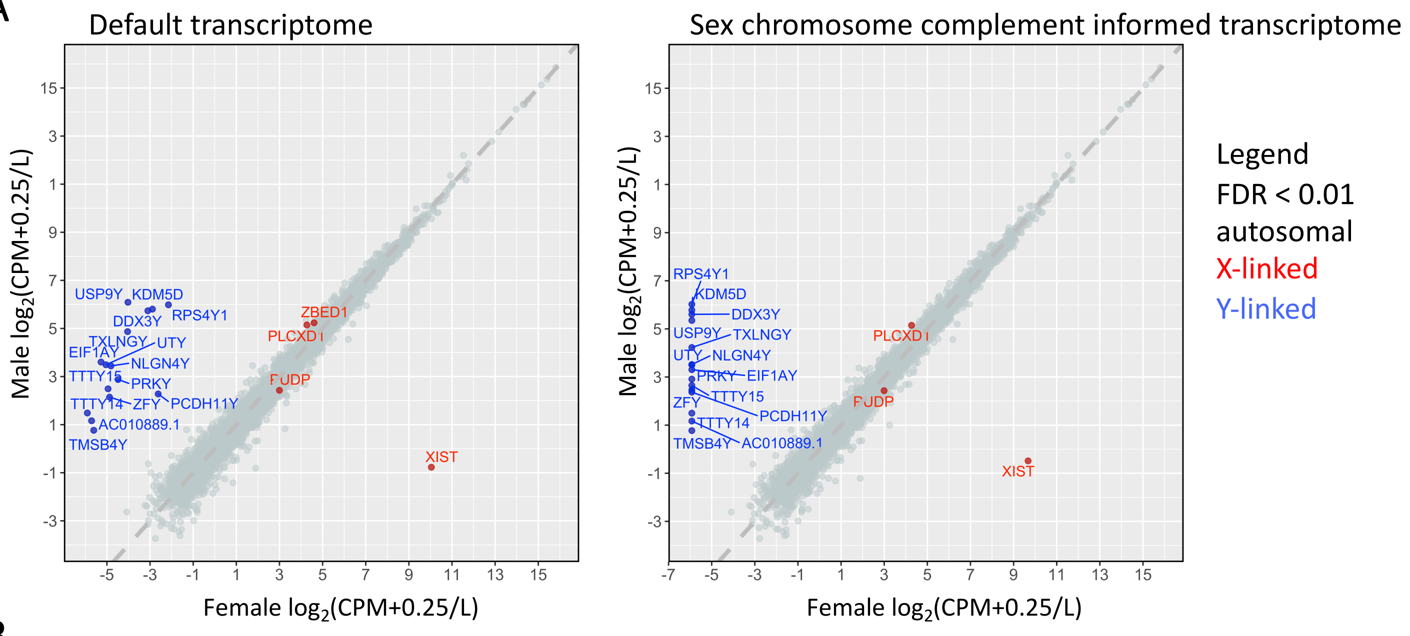


B

| Genes | Total | Unique |
|---|---|---|
| Autosomal | 0 | 0 |
| X-linked | 2 | 1 |
| Y-linked | 15 | 0 |

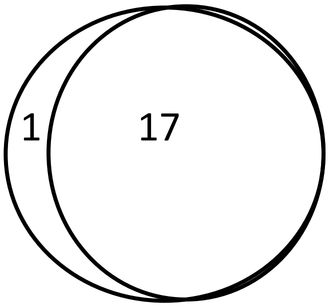


| Genes | Total | Unique |
|---|---|---|
| Autosomal | 0 | 0 |
| X-linked | 2 | 0 |
| Y-linked | 15 | 0 |
