## Additional file 20 for "Reference genome and transcriptome informed by the sex chromosome complement of the sample increases ability to detect sex differences in gene expression from RNA-Seq data"

### **Methods**

*RNA-Seq samples*

From the Genotyping-Tissue Expression (GTEx) Project data, we downloaded SRA files for brain cortex and whole blood from 3 genetic female (46, XX) and 3 genetic male (46, XY) individuals [[19,35]](https://paperpile.com/c/pGSVvm/yQP9r+DqyNW) table 1. The samples were processed in the exact same way as the samples in the main manuscript.

### **Results**

*RNA-Seq reads aligned to autosomes do not vary much between reference genomes*

Reads aligning to chromosome 8 didn’t vary tremendously or at all between aligning to the default versus sex chromosome complement informed reference for both brain cortex and whole blood tissues. Specifically, chromosome 8 increased from 1,479,544 reads to 1,479,584 on average across both male and female samples in brain cortex when mapped to a sex chromosome complement informed using HISAT (Table 2). A female XX brain cortex sample showed the largest chromosome 8 mapped read increase when aligned to a sex chromosome complement informed reference genome (334 more reads with HISAT; 608 more reads with STAR; Table 2). The sample that showed the highest decrease in mapped reads was a male XY brain cortex sample, which showed 5 fewer reads on chromosome 8 when aligned to a reference genome informed on the sex chromosome complement using HISAT read aligner. There was no decrease in chromosome 8 mapped reads when aligned to a sex chromosome complement informed reference genome using STAR read aligner for male or female whole blood samples.

*Reads aligned to the X chromosome increase in both XX and XY samples when using a sex chromosome complement informed reference genome*

We found that when reads were aligned to a reference genome informed by the sex chromosome complement for both male XY and female XX brain cortex and whole blood samples, reads on the X chromosome increased by ~0.13%. On average there was 1,627 more reads aligning to the X chromosome when using a sex chromosome complement informed reference genome. Reads on the Y chromosome decreased 100% (46,852 reads on average) in female XX samples and by ~63.43% (52,327 reads on average) in male XY samples for brain cortex and whole blood when aligned using HISAT (Table 3). Similar increases in X chromosome and decreases in Y chromosome reads when aligned to a sex chromosome complement informed reference was observed for when STAR was used as the read aligner for both male XY and female XX brain cortex and whole blood samples (Table 3).

*Aligning to a sex chromosome complement informed reference genome increases the X chromosome PAR1 and PAR2 expression*

There is an increase in X chromosome PAR1 and PAR2 expression when reads were aligned to a reference genome informed on the sex chromosome complement. This is true for both male XY and female XX samples from the brain cortex (Figure 1) and whole blood samples using either HISAT or STAR as the read aligner (Figure 1). We found an average of 2.83 fold increase in expression in PAR1 expression for female XX brain cortex samples and a 2.71 fold increase in expression for male XY brain cortex samples using HISAT read aligner (Table 4). XTR in female XX brain cortex samples showed a 1.49 fold increase in expression and no change in male XY brain cortex samples. PAR2 showed an average of 2.47 fold increase for female XX brain cortex samples, while there was a 2.19 fold increase in PAR2 for male XY brain cortex samples using HISAT read aligner. Similar results were obtained using STAR read aligner (Table 4).

*X and Y homologous genes showed little to no increase in expression when mapped to a sex chromosome complement reference genome compared*

X and Y homologous genes showed little to no increase in expression when mapped to a sex chromosome complement reference genome compared to mapping to a default reference genome, with the exception of *PCDH11X* gene (Table 5). *PCDH11X* showed a 1.50 fold increase (0.58 log_2_ fold increase) in female XX brain cortex samples, with a similar increase of 1.47 fold (0.55 log_2_ fold increase) in female XX whole blood samples using HISAT read aligner. A similar increase in expression for *PCDH11X* was observed for both female XX brain cortex and whole blood samples when STAR was used as the read aligner (Table 5). *PCDH11X* showed no differences in expression between reference genomes for male XY brain cortex and whole blood samples when using either HISAT or STAR (Table 5).

*A sex chromosome complement informed reference genome increases the ability to detect sex differences in gene expression*

At an adjusted p-value of 0.05 and aligning with HISAT, we find 12 new genes (all on the Y chromosome) that are only called as differentially expressed between the sexes in the brain cortex when aligned to reference genomes informed on the sex chromosome complement (Figure 4). We find 49 genes (48 autosomal and 1 X-linked) that are called as differentially expressed between the sexes in the brain cortex that are no longer called when using a sex chromosome complement for aligning reads. We observed similar trends in changes for differential expression between male XY and female XX whole blood samples using either HISAT or STAR as the aligner (Figure 2; Table 6).

*Using sex-linked genes alone is inefficient for determining the sex chromosome complement of a sample*

Using the default reference genome for aligning male XY and female XX brain cortex and whole blood samples, we observed a small number of reads aligning to the Y-linked genes in female XX samples, but also observed clustering by sex for *DDX3Y* and *XIST* gene expression (Figure 2). Male XY samples showed expression for *DDX3X* and *DDX3Y* (greater than 5 log_2_(CPM+2/L)). Female XX samples showed expression for *XIST* greater than 2.5 log_2_(CPM+2/L), and male XY samples showed little to no expression for XIST (less than 0 log_2_(CPM+2/L)). In contrast to the default reference genome, when aligned to a sex chromosome complement informed reference genome, samples cluster distinctly by sex for *DDX3Y*, *PCDH11Y*, *USP9Y*, *ZFY*, *UTY*, and *XIST*, all showing at least a 2.5 log_2_(CPM+2/L) difference between the sexes (Figure 3).

### **Discussion**

*Increase in X chromosome expression for both male and female samples*

Measurements of X chromosome expression increase for both male XY and female XX brain cortex and whole blood samples when aligned to a sex chromosome complement informed reference genome (Figure 1). There was a minimum 1.6 fold increase in expression for the genes in PAR1 and PAR2 for all samples (male and female) when aligned to a sex chromosome complement informed reference genome (Figure 1; Table 4). We see that the XTR has a minimum 1.35 fold increase in expression for female brain cortex and whole blood samples when aligned to a sex chromosome complement informed reference genome in comparison to a default reference genome. However, there was no change observed in males.

*Gene enrichment analysis alters depending on choice of reference genome*

GO enrichment analysis of genes that are more highly expressed in brain cortex male samples than females, when samples were aligned to a default reference genome, were genes involved in germline cell cycle switching and mitotic to meiotic cell cycle (Table 7). However, when these samples were aligned to a sex chromosome complement informed reference genome, genes upregulated in males were enriched for positive regulation of transcription from RNA polymerase II promoter in response to heat stress and GO component of specific granule lumen, which are secretory vesicles of the immune system.

### **Figures legends**

**Figure 1. X chromosome expression differences between default and sex chromosome complement informed alignment.** Differences in X chromosome expression between reference genomes for male XY and female XX samples aligned using HISAT for the whole X chromosome and the first 5Mb are shown for the brain cortex (**A** and **B**, respectively), and the whole blood (**C** and **D**, respectively). Differences in X chromosome expression between reference genomes for male XY and female XX samples aligned using STAR for the whole X chromosome and the first 5Mb are shown for the brain cortex (**E** and **F,** respectively), and the whole blood (**G** and **H**, respectively).

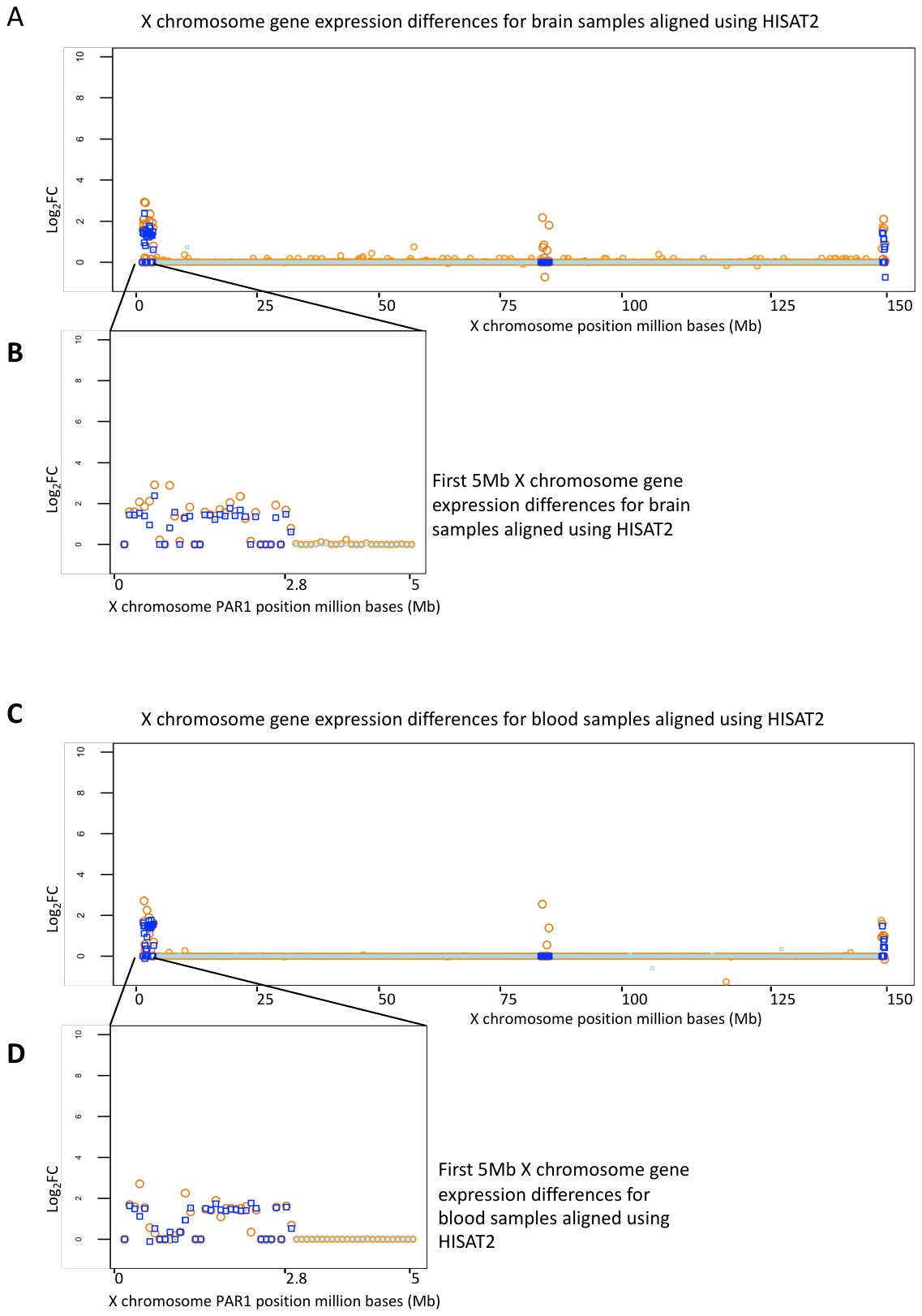
**Figure 2.** **Gene expression differences between male XY and female XX samples.** Sex differences in gene expression for brain cortex and whole blood samples for when samples were aligned to a default reference genome and a to a reference genome informed on the sex chromosome complement. Showing sex differences in gene expression between reference genomes used for alignment and for when samples were aligned using HISAT and STAR.

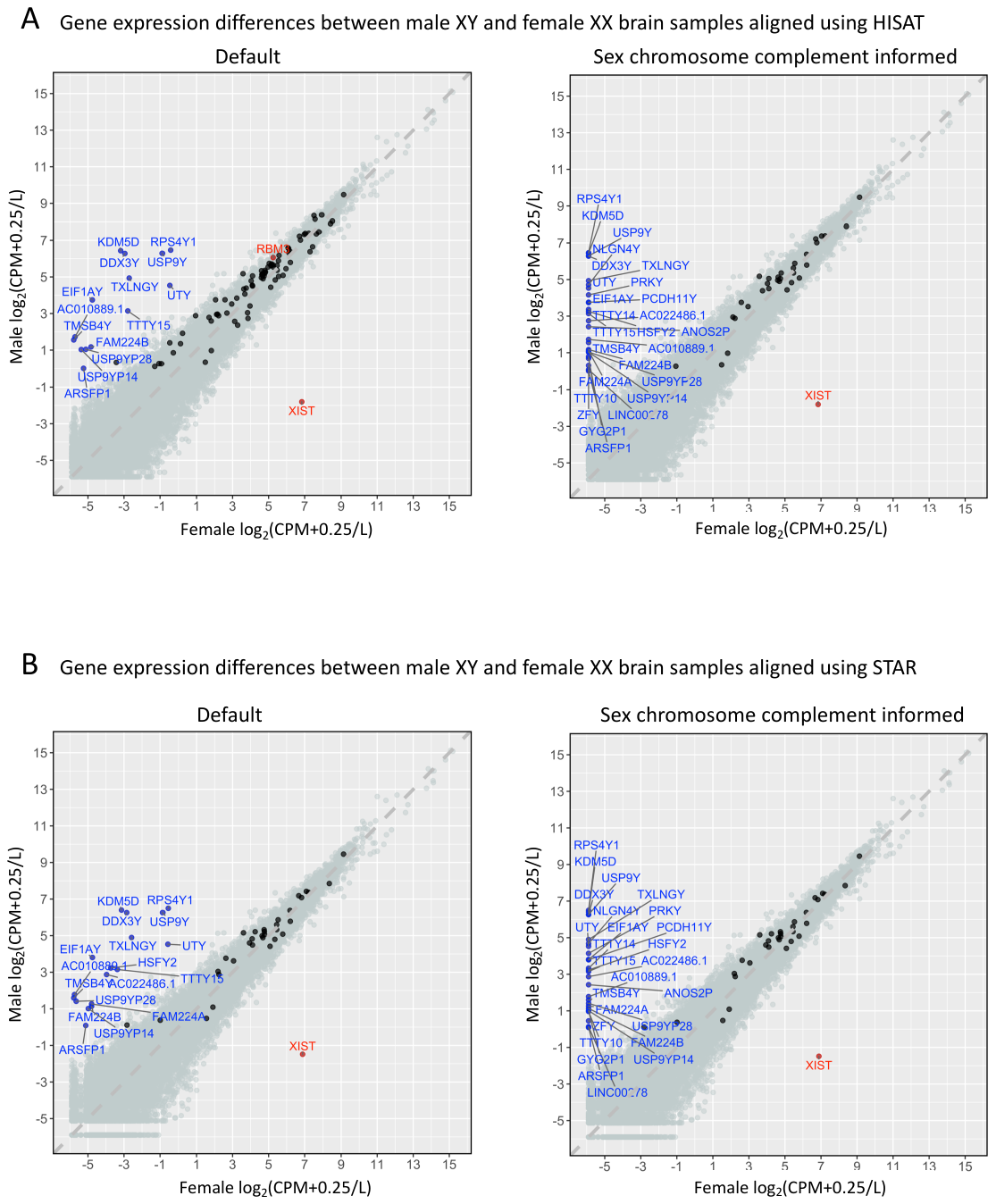
**Figure 3. Genetic sex of RNA-Seq samples.** Gene expression log2(CPM+0.25/L) for select XY homologous genes and XIST and SRY for when reads were aligned to a default reference genome A) and C) using HISAT and STAR, respectively, and for when reads were aligned to a sex chromosome complement informed reference genome B) and D) using HISAT and STAR, respectively. Male XY brain cortex and whole blood samples are shown in blue squares and female XX brain and blood samples shown in red circles.

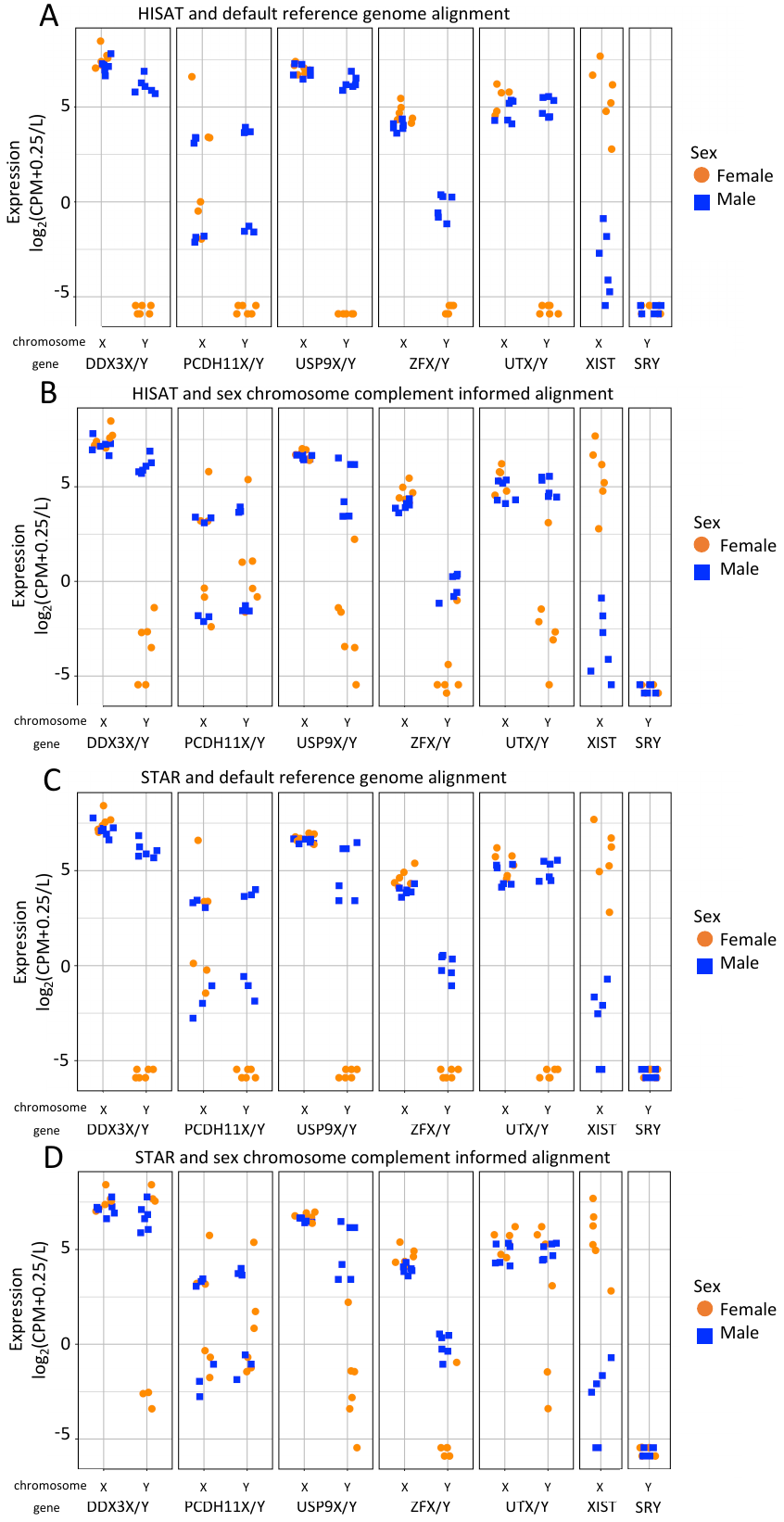

### **Table legends**

| **Sex** | **Individual_SRR** | **Brain_SRR** | **Blood_SRR** |
| --- | --- | --- | --- |
| Female | 678158 | SRR1475168 | SRR614011 |
| Female | 590962 | SRR602927 | SRR1351166 |
| Female | 590921 | SRR612575 | SRR809283 |
| Male | 721013 | SRR1081741 | SRR1074646 |
| Male | 847758 | SRR1081741 | SRR1090907 |
| Male | 984961 | SRR1310008 | SRR1345266 |

**Table 1. Sample IDs.** RNA-Seq brain cortex and whole blood tissue samples from 3 genetic female (46, XX) and 3 genetic male (46, XY) individuals were downloaded from the Genotype-Tissue Expression (GTEx) project [[19,31]](https://paperpile.com/c/pzklmk/pp3H9+Y6DzG) for a total of 12 RNA-Seq tissue samples.

**Table 2. Table of mapped read statistics for default and sex chromosome complement informed alignment.** Total reads mapped in brain cortex and whole blood female XX and male XY samples for when reads were aligned to a default reference genome and for when reads were aligned to a reference genome informed on the sex chromosome complement. Total mapped reads for HISAT and STAR as the read aligner. Additional file 18A

**Table 3. X chromosome gene expression values per sample, aligner and reference genome used for alignment.** CPM values for male XY and female XX brain cortex and whole blood samples when aligned to a default and sex chromosome complement informed reference genome for chromosome X. A text format of CPM values is available on GitHub, https://github.com/SexChrLab/XY_RNAseq.

**Table 4. X chromosome regions mean and median expression values.** X chromosome regions PAR1, PAR2, XTR, XDG, XAR, and XCR mean and median CPM expression for male XY and female XX brain cortex and whole blood samples when aligned to a default or sex chromosome complement informed reference genome using HISAT or STAR.

**Table 5. Gene expression for XY homologous genes.** X chromosome expression for 26 X and Y homologous genes. Difference in gene expression for when male XY and female XX brain cortex and whole blood samples were aligned to a default and sex chromosome complement informed reference genome. Little to no difference in gene expression between default and sex chromosome complement informed reference genome alignment was observed for 25 of the 26 X and Y homologous genes for both male XY and female XX brain cortex and whole blood samples using either HISAT or STAR. PCDH11X showed a 1.50 and 1.47 fold increase in expression for brain cortex and whole blood, respectively, in female XX samples using HISAT read aligner with similar results for STAR. XY male brain cortex and whole blood samples showed little to no differences in gene expression between reference genomes for the 26 X and Y homologous genes using either HISAT or STAR.

**Table 6**. Differentially expressed genes between the sexes that were uniquely and jointly called between reference genomes. Genes that are differentially expressed between the sexes, male XY and female XX, for brain cortex and whole blood samples. Differentially expressed genes that are uniquely called when using either the default or sex chromosome complement informed reference genome and differentially expressed genes that jointly called between the reference genomes.

**Table 7. Gene enrichment analysis.** Gene enrichment analysis of genes that are more highly expressed in one sex verses the other sex for when samples were aligned to a default or sex chromosome complement informed reference genome using either HISAT or STAR.
